# Multi-scale dissection, compaction and derivatization of mammalian developmental enhancers

**DOI:** 10.64898/2026.04.20.719625

**Authors:** Jean-Benoît Lalanne, Tony Li, Emma A.N. Kajiwara, Chau Huynh, Tiffany V. Do, Beth K. Martin, Samuel G. Regalado, Jay Shendure

## Abstract

Gene expression during mammalian development is orchestrated by non-coding *cis*-regulatory DNA elements (CREs) such as distal enhancers^1–3^. Despite their fundamental importance, and notwithstanding recent progress in predictive modeling^4–9^, many high-level properties of enhancer ‘grammar’ remain unresolved. How does the length of an autonomously active CRE constrain its activity? How robust are CREs to mutations or rearrangements of transcription factor binding sites (TFBSs)? And how much epistasis exists among these sites? As predictive models solely trained on endogenous CREs are unlikely to resolve these questions^10^, we subjected several endogenous CREs to intensive sequence-level perturbation. Specifically, we assayed >35,000 variants of 5 parietal endoderm enhancers, with variants organized into four perturbation classes, designed to probe: (i) the functional sufficiency of sub-fragments via dense multi-size tiling, (ii) local epistasis via multi-hit saturation mutagenesis, (iii) activity-size tradeoffs via model-guided compaction, or (iv) functional resilience via sequence derivatization anchored on key TFBSs, including random deposition, reconstitution, and synthetic thripsis. This multi-scale dissection revealed rich phenomena. Sub-tiling uncovered sharp non-additivity between activity and fragment size, highlighting strongly synergistic TFBS clusters. Compaction showed that natural CREs lie far from the activity-size Pareto front, and that model-guided deletions can yield shorter yet stronger elements. Mutational scanning exposed a spectrum of CRE robustness, from tolerant to fragile, together with rare but consequential epistasis between individual TFBSs. Finally, TFBS-anchored derivatization demonstrated that ‘background’ sequence can influence activity on par with TFBS arrangement. Strikingly, a substantial fraction of CRE derivatives exceeded the activity of their endogenous progenitors. Taken together, these results reveal both ‘soft’ and ‘stiff’ directions in regulatory sequence space, advancing a quantitative phenomenology of how enhancer sequences encode function and robustness.

## INTRODUCTION

Developmental enhancers are among the most evolutionarily dynamic components of a mammalian genome^1–3^. On one hand, they are strikingly robust to mutation—far more tolerant of sequence change than protein-coding genes—allowing them to change rapidly^11^. On the other hand, their ability to drive phenotypic novelty arises from a sensitivity to mutation^12,13^. This duality presumably reflects a highly rugged mapping between DNA sequence and regulatory function, and underscores how little we understand about their underlying grammar. How, at the level of primary sequence, do enhancers encode this distinctive combination of robustness and responsiveness? In the context of development, enhancers orchestrate precise, cell type–specific gene expression programs by integrating the activity of lineage-defining transcription factors within a given trans-acting, cell-type-specific milieu^14,15^. Many enhancers are functionally autonomous: when isolated from their genomic context and tested in the appropriate cell type, they can substantially increase transcription from a minimal promoter^16,17^. This autonomy underlies both conventional and massively parallel reporter assays (MPRAs).

A central goal in regulatory genomics is to achieve a predictive understanding of how DNA sequence encodes patterns of gene expression across developmental contexts. Recently, machine learning models, including convolutional neural networks^18,19^ (CNNs), transformers^8,9,20^, diffusion models^21^, and foundation models^22^, have made impressive strides in predicting biochemical profiles or enhancer activity directly from DNA sequence^4–7^, and in some cases have been used to design synthetic enhancers with targeted activities^23–27^. However, predictive accuracy does not equate to mechanistic understanding. The internal representations of these models remain difficult to interpret in genetic or biochemical terms^28–31^, and their performance has not yet translated into generalizable principles of *cis*-regulatory grammar—such as how activity scales with size, how robustness to mutation is encoded, or how combinations of TFBSs interact. Because genomes are the products of evolution and incompletely sample the vast regulatory design space, training models on endogenous enhancers alone is unlikely to yield such rules^10^. Generating the empirical data needed to fully parameterize such models remains an open challenge.

An alternative to modeling natural sequences is to use MPRAs to interrogate large libraries of designed elements. This strategy has already yielded important insights: random sequence libraries have uncovered unexpected routes to enhancer activity^6^, heuristically designed CREs have probed flexibility and sufficiency^32–37^, and saturation mutagenesis has mapped nucleotide-level contributions^38,39^. Such studies illustrate the power of assaying designed sequences, but most have focused on a single perturbation mode at a time, and not all directions have been explored.

We hypothesized that a multi-scale perturbational dissection of endogenous enhancers could expose organizing principles of *cis*-regulatory grammar that remain inaccessible through natural sequence variation, single-axis perturbational studies, or fully synthetic allelic series. To this end, we performed an intensive analysis of 5 parietal endoderm-specific, autonomous developmental enhancers recently identified via single-cell reporter screens^40^. Using MPRAs, we designed, synthesized, and functionally assayed >35,000 sequence variants that fragmented, compacted, mutated, or reconstituted their original architectures. These variants spanned four perturbation classes (**Fig. 1**): dense multi-size tiling, multi-hit saturation mutagenesis, model-guided compaction, and sequence derivatization anchored on key TFBSs. By interrogating sequence-function relationships along these axes, we sought not only to catalog variant effects, but also to uncover the organizing principles—and limits—of their underlying grammar, thereby advancing a quantitative phenomenology of enhancer function and robustness.

**Figure 1.**
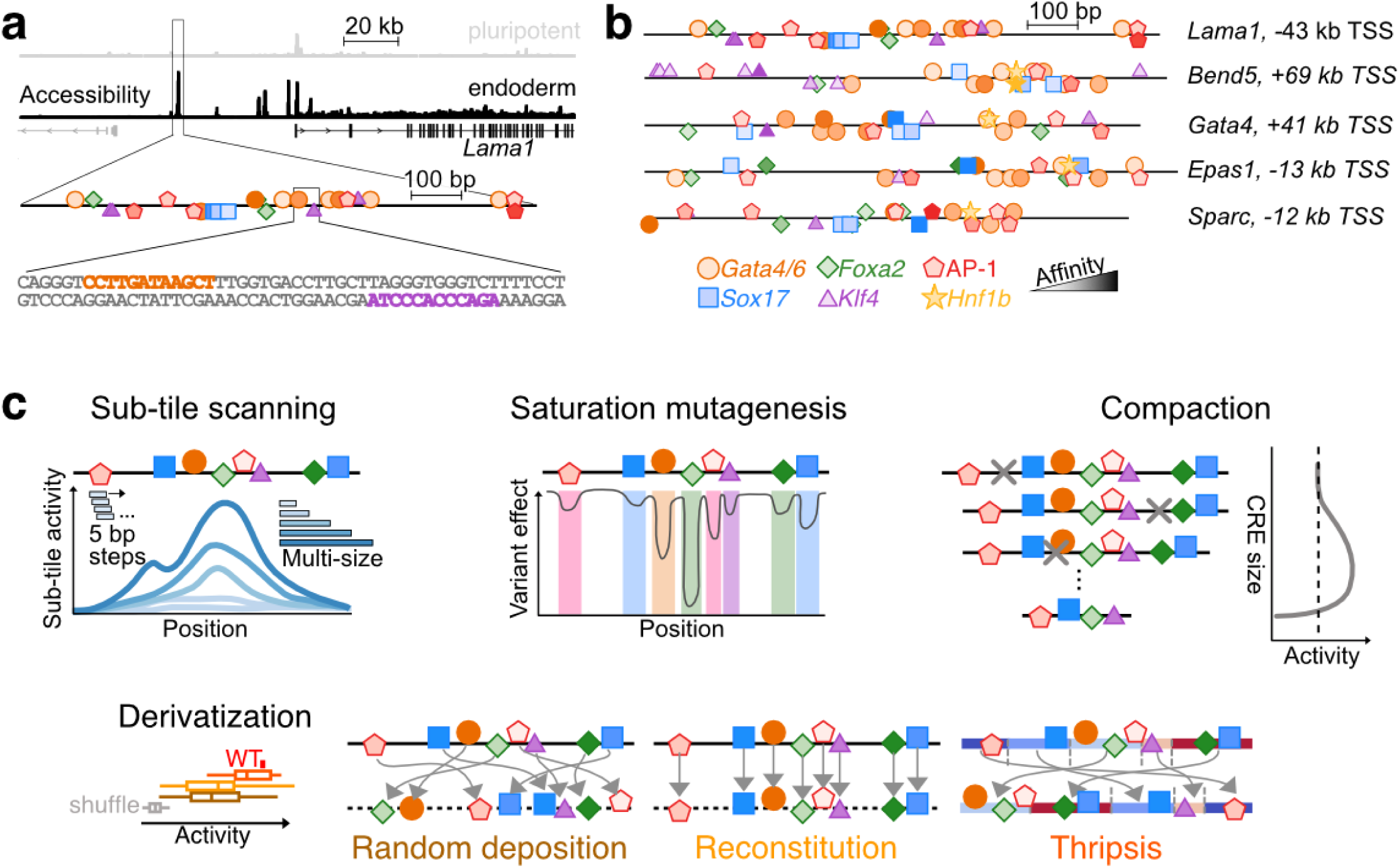
Multi-scale dissection, compaction and derivatization of mammalian developmental enhancers. **(a)** Chromatin accessibility at the *Lama1* locus based on scATAC-seq data from differentiating mouse embryoid bodies (EBs)^40^. A peak exhibiting accessibility specifically within parietal endoderm cells is highlighted, with predicted TFBSs marked. The sequence corresponding to this peak is an autonomous enhancer, in that it drives cell-type-specific activity in a reporter assay. **(b)** The predicted TFBSs of four additional CREs, each an autonomous, parietal endoderm-specific enhancer, are shown. Including the example shown in panel **a**, these five endogenous CREs are our models for the multi-scale phenotyping reported here. **(c)** Schematic of the different classes of CRE variants generated for our multi-scale phenotyping of enhancers: dense multi-size sub-tiling, multi-hit saturation mutagenesis, model-guided compaction, and three modes of CRE derivatization (random deposition, reconstitution, and synthetic thripsis).

## RESULTS

### Endoderm-specific CREs as models for deep dissection

We sought to leverage MPRAs^16,17^ to deeply dissect multiple aspects of enhancer grammar in a limited number of endogenous CREs. The number of cell type-specific developmental enhancers that are well-established to be autonomously active in reporter assays remains limited^41–45^. We therefore turned distal CREs identified in our recent single-cell quantitative reporter (scQer) screen in mouse embryoid bodies (EBs)^40^, and in particular ones that exhibited high activity, cell type-specificity and autonomous activity in the parietal endoderm lineage. Situated near important genes (**Figs. 1**, **S1a**; *e.g.* developmental transcription factor Gata4; structural proteins Laminin-α1 and Osteonectin), these CREs also harbour numerous binding sites for TFs with established roles in *cis*-regulation of the extraembryonic endoderm^46–50^ (**Figs. 1b**, **S1b-c**; *Gata4/6*, *Sox17*, *Foxa2*, *Klf4*, *Hnf1b*, AP-1 factors). As a scalable biotype, we leveraged PYS-2 mouse cells, which model extraembryonic parietal endoderm^51,52^. Bulk ATAC-seq data confirmed excellent correlation with accessibility profiles from parietal endoderm from mouse embryos (*ρ* = 0.64 vs. *ρ* = 0.20-0.26 for other lineages)^53^.

To identify shorter functional tiles for further dissection within the ∼0.9 kb x 10 developmental CREs evaluated by scQer^40^, we synthesized 270 bp tiles scanning their full lengths with a 5 bp stride. These tiles were cloned into an MPRA vector bearing a degenerate barcode (BC) positioned in the 5’ UTR of a reporter gene^54^. Upon sequencing to associate BCs with tiles, we recovered 95% of designed tiles (total library complexity: ∼660,000 BCs, median 492 BC per tile, 1306/1377 tiles associated with ≥10 BCs; **Fig. S2a**). Plasmid libraries were transfected in both PYS-2 (modeling parietal endoderm; 5 replicates) and mESCs (modeling pluripotency; negative control in that activity not expected; 2 replicates) cells, and RNA and DNA were extracted 48 hrs post transfection. BCs were amplified with UMIs and sequenced, separately from RNA and DNA, to quantify their relative abundances. Tile activities were calculated by dividing the summed UMIs of associated BCs observed in RNA by the summed UMIs of associated BCs observed in DNA. Tile activities of transfection replicates were highly correlated (R^2^ [on log-transformed values] > 0.85; **Fig. S2b-c**). This basic strategy for conducting and analyzing bulk 5’ episomal MPRAs is used throughout this manuscript to quantify regulatory activity.

Remarkably, we observed very high activities (15– to 109-fold signal-to-noise) for specific sub-tiles of five of the endoderm CREs, and in four cases, substantially higher than that of the full-length CRE (top row, **Fig. S2d**). Furthermore, these sub-tiles were inactive in mESCs, consistent with the associated region’s parietal endoderm-specific accessibility (**Fig. S1a**). The five developmental CREs with such high-activity sub-tiles (**Table S1**) were taken as the starting point of a comprehensive multi-scale program leveraging bulk MPRA to probe the phenomenology of regulatory sequence-to-function maps (**Fig. 1c**). In a companion manuscript^53^, we densely mapped the functional evolution of these same enhancers across 480 extant and ancestrally reconstructed mammalian genomes, providing complementary insights (Li, Lalanne *et al.* <u>bioRxiv</u> 2026).

### Dense, multi-scale sub-tiling of developmental CREs charts non-linear size-vs-activity relationships

What size of DNA is required for a mammalian enhancer? A single TFBS can be functional (*e.g.* for TP53 ref.^55^), and short synthetic arrangements of a few TFBSs (<50 bp) can act as strong cell-type-specific enhancers^23,26^. However, such short sequences presumably face inherent limits with respect to the magnitude and specificity of activity they can achieve. Previous MPRA studies described high-resolution tiling of endogenous regulatory elements with a fixed tile size (145-200 bp)^24,56,57^, but to our knowledge, none profiled variably sized sub-tiles of a given CRE to systematically map the relationship between size and activity. Other studies assayed genomic DNA fragments and thus *de facto* tested multiple sizes^58,59^, but lacked the coverage and resolution to address this question. Finally, while nonlinear interactions have been observed in synthetic TFBS arrays^32–34,60,61^, it remains unclear whether—and over which length-scales—such interactions shape the activity of endogenous CREs.

To systematically investigate size-vs-activity and nonlinearities in endogenous enhancers, we measured the activity of ∼10,000 sub-tiles derived from our five model CREs at high density (5 bp stride), multiple lengths (seven sizes ranging from 40 to 300 bp), and in both orientations (**Fig. 2a**). Measurements were obtained for nearly all designed sub-tiles (98.5%) via bulk MPRAs performed in PYS-2 cells (**Fig. S3a**). As these measurements were highly reproducible (R^2^ [on log-transformed activities] = 0.71-0.83 between biological replicates), we used the average activity across replicates in the analyses described below (**Fig. S3b**).

**Figure 2.**
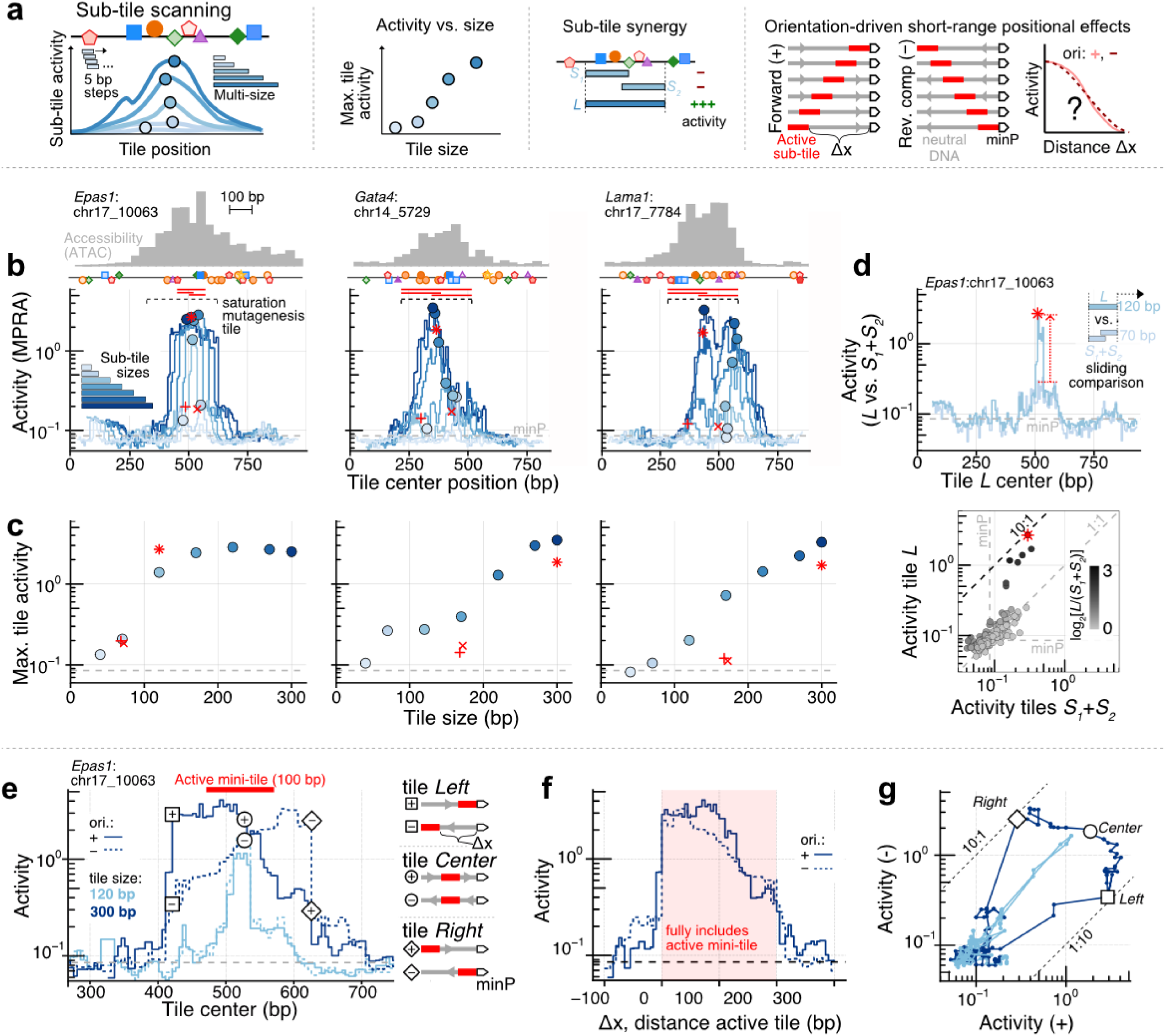
Multi-sized sub-tiling reveals regulatory synergies and orientation-dependent short-range positional effects. **(a)** Schematic of multi-sized sub-tiling MPRA workflow. **(b)** Top: chromatin accessibility across three of the profiled CREs (pseudobulked scATAC-seq from parietal endoderm cells)^40^. Middle: predicted TFBSs for the parietal endoderm *cis*-regulatory module, as in **Fig. 1**; only sites with normalized affinity >0.2 are shown. Bottom: sub-tiling MPRA profiles using a 5-bp stride across each CRE for seven tile sizes (40-300 bp; darker blue indicates larger tiles). Each trace represents the average activity of forward and reverse orientations (arbitrary units) and is smoothed with a 3-bp sliding median filter. The position of maximum activity for each tile size within the corresponding ATAC peak is marked by a circle (see panel **c**). Black dashed brackets above indicate the 300-bp region subjected to saturation mutagenesis (**Fig. 3**). Red vertical lines above denote the positions of short and long tiles exhibiting synergy; in panel **c**, the corresponding tiles are marked with red symbols (short S*_1_*, ‘+’; short S*_2_*, ‘×’; long L, ‘✴’). The grey dashed line indicates the activity of the basal minP-only control. A set of similar plots that includes all five CREs is shown in **Fig. S4c**. **(c)** Maximum activity for each tile size within the accessible ATAC peak, computed from the smoothed data in panel **b**. Red symbols indicate the *S_1_, S_2_* and L tiles highlighted in panel **b**. For the *Epas1*:chr17_10063 CRE, the *L* tile (value from non-smoothened data) exceeds the maximum 120-bp activity because the smoothing procedure used in panel **b** slightly dilutes the signal. A set of similar plots that includes all five CREs is shown in **Fig. S4d**. **(d)** Sliding-window comparison of the summed activity of two adjacent short tiles (*S_1_*+*S_2_* – minP) vs. the corresponding long tile (L) spanning the same region (matched by edge position and overlapping slightly internally), sized 70 bp and 120 bp respectively. Divergence between traces indicates regulatory synergy (activity of L exceeding *S_1_*+*S_2_*) for CRE *Epas1:*chr17_10063. Red symbols mark the triplet tiles highlighted in panels **b** and **c**, which display a 9-fold synergy. Synergy maps for other CREs shown in **Fig. S5**. **(e)** Zoomed-in view of activity profiles for CRE *Epas1:*chr17_10063 for 120-bp (light blue) and 300-bp (dark blue) tiles, now stratified by orientation (forward, solid; reverse, dashed) and smoothed with a 3-bp sliding median filter. The 120-bp tiles show a sharp peak in activity near tile-center position 525 bp that is roughly 20 bp wide, corresponding to an active ‘mini-tile’ of 100 bp (red bar above). Notably, the activities of 120-bp tiles are not dependent on orientation and demonstrate similar maximal activities (light blue lines, solid vs. dashed). In contrast, while the 300-bp tiles show similar maximum activity as the shorter tiles (slight difference in activity arises from smoothing procedure; see panel **c**), they exhibit a marked orientation asymmetry for tiles harbouring the active 100-bp mini-tile (dark blue lines, solid vs. dashed). This asymmetry is drastic for the 300-bp tiles in which the active mini-tile is at either boundary (tile *Left* square-enclosed + vs. –; tile *Right* diamond-enclosed + vs. –; schematic to the right of figure), but absent for the 300-bp tile in which the active mini-tile is centrally located (tile *Center,* circle-enclosed + vs. –). **(f)** Same 300-bp tile data from panel **e**, but replotted such that the x-axis now corresponds to the distance between the mini-active tile and the minimal promoter (Δx). In contrast with panel **e**, the forward and reverse orientation profiles now overlap. **(g)** Direct comparison of forward and reverse orientation activities for matched tiles (*i.e.* differing only by orientation) across the *Epas1*:chr17_10063 CRE scan. The opening of a gap for 300-bp (dark blue) but not 120-bp (light blue) tiles reflects the orientation-dependent positional effect observed in panel **e**. Edge and central tiles are indicated by shapes as panel **e** (*Left*: square, Center: circle, *Right*: diamond).

Initially averaging across both orientations of any given tile, several patterns were immediately clear (**Figs. 2b-c**, **S4**). <u>First</u>, apart from rare isolated tiles (**Fig. S4b**, black arrows), domains of broad functional activity overlapped with accessible regions, with maximal activity and maximal accessibility closely aligned (**Figs. 2b**, **S4c**). Thus, centering on accessibility summits is a valid heuristic for prioritizing sequences for autonomous activity, similar to findings from non-coding CRISPR screens^62^. <u>Second</u>, very short endogenous tiles overwhelmingly exhibited very low activity (**Fig. 2c**; top row of **Fig. S4b**; only 0.5% of 40-bp tiles were >2-fold above minP-only controls). <u>Third</u>, activity increased with sub-tile size near-monotonically for all CREs, but the maximum was reached at different thresholds. Activities of two of the CREs plateaued with relatively short fragments (220 bp for *Bend5*:chr4_8201; 170 bp for *Epas1*:chr17_10063), while for the other CREs, activity continued to increase substantially at longer lengths (**Figs. 2c**, **S4d**). This suggests more context than 200 bp may often be necessary to accurately model the full activity of accessible peaks via MPRAs.

Turning to the shape of size-vs-activity profiles, we observed clear CRE-specific non-linearities (**Figs. 2c**, **S4d**). To localize the sequence determinants underlying these regulatory synergies, we compared the activity of short and long tiles spanning identical regions. Specifically, we calculated the summed activity of two adjacent short tiles (*S_1_+S_2_*) whose outer boundaries matched those of a longer tile (*L*), with a small central overlap larger than one TFBS (top panel of **Fig. 2d**). Scanning across all CREs, we identified focal regions of pronounced non-additivity, in which the long tile’s activity exceeded the summed activity of its constituent short tiles by 7.2-to 15.3-fold (**Figs. 2d**, **S5**). Remarkably, synergistic *S_1_* and *S_2_* tiles were themselves rich in functional TFBSs (see saturation mutagenesis data below: **Figs. 4**, **S9**), suggesting that cooperative interactions among clustered binding sites underlie these nonlinear effects. Taken together, these results show that endogenous enhancer activity depends on element-specific, cooperative TF interactions rather than the additive contributions of individual sites, similar to what has been reported in synthetic TFBS arrays^32–34,60,61^.

### Orientation-dependence reveals short range CRE-to-promoter positional effects in MPRAs

Our comprehensive sub-tiling library included both forward and reverse-complement orientations of all profiled regulatory fragments (**Fig. S4a**). These afford a high-resolution assessment of the consequences of reversing the orientation of CREs in the context of MPRAs. Although we observed broad overall agreement between orientations, in line with the canonical definition of enhancers^63^, closer inspection revealed differences that were exacerbated for larger tiles (**Fig. S4b**). Previously, large-scale libraries also included tiles in both orientations and also noted slight quantitative differences^54,64,65^. However, because those studies assayed discontiguous and/or fixed-length fragments, they could not resolve how orientation effects depend on fragment size or local regulatory context, nor offer mechanistic insight into their origin.

In contrast, our variable-length, contiguous sub-tiling enabled us to localize these orientation-dependent effects within individual CREs. A notable example was the *Epas1*:chr17_10063 CRE (**Fig. 2e**). For this element, the majority of activity is recapitulated by a short 100 bp subregion, evidenced by jumps in activity in 120 bp tiles bearing this subsequence (**Figs. 2b-c**, **S4b**) that coincide with *S_1_+S_2_*synergy (**Fig. 2d**) and multiple functional TFBSs (**Fig. 3a**). Surprisingly, although 120 bp tiles bearing the active subregion do not exhibit orientational dependence, 300 bp tiles clearly do (**Fig. 2e**). At the extreme, 300 bp tiles in which the active subregion is present at one end vs. the other exhibit a 10-fold difference in activity, with higher activity associated with its proximity to the minP promoter (see *Left* and *Right* tiles, **Fig. 2e**). In contrast, 300 bp tiles in which the active subregion is centrally located do not exhibit such orientational dependence (*Center,* **Fig. 2e**). This suggests that at least in this specific case, distance from minP is a major contributor to variance in the activity of the 100 bp subregion when it is embedded within larger tiles.

**Figure 3.**
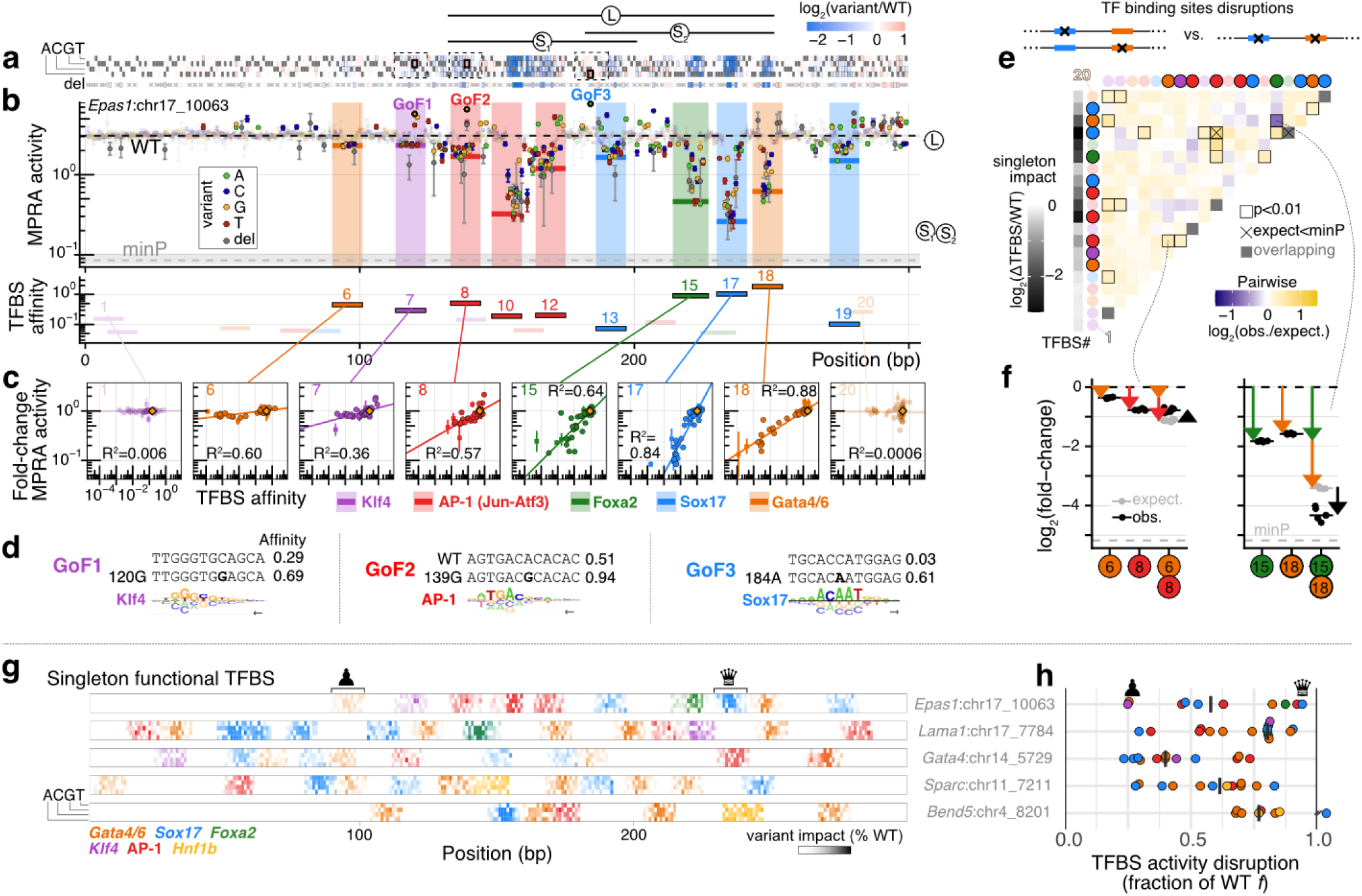
Saturation mutagenesis yields a high-resolution, quantitative map of developmental TFBS. **(a)** Heatmap of variant effects on activity for the *Epas1*:chr17_10063 CRE, shown as log_2_ fold-changes for variants vs. wildtype (WT). Each position is associated with three possible substitutions and one deletion (rows). Variants with insufficient barcode support (<5 BCs) or ambiguous interpretation (*e.g.* deletions within homopolymers) are marked with X’s. Variants with large effect sizes (|log_2_FC| > 0.32 & Wilcoxon test with Benjamini-Hochberg correction: FDR < 0.006) have thinly bolded edges. Three mutations highlighted in panel **d**, GoF1-3, have thickly bolded edges. Positions of synergistic *S_1_*, *S_2_*, *L* tiles from **Fig. 2d** shown at top. **(b)** Top: dot-plot showing variant activity as a function of position. WT activity is marked by a dashed black line. Basal minP-only control activity is marked by a dashed gray line (mean) and shading (range across for replicates). Large effect size variants, as defined in panel **a**, again have their edges bolded. Activities of synergistic *S_1_*, *S_2_*, *L* tiles from **Fig. 2d** shown at right. Positions corresponding to singleton-functional TFBS, as defined in panel **c** legend and text, are shaded by TF identity. Bottom: positions of all TFBSs with predicted WT normalized affinities ≥ 0.05, coloured by TF identity. Non-singleton-functional sites are faded. **(c)** Correlation plots between fold-change in predicted TFBS affinity (x-axes) vs. fold-change in MPRA activity (y-axes) for selected TFBS. Deletions are represented as squares and point mutations as circles. TFBS with an R^2^ > 0.195 for substitutions are designated ‘singleton-functional’. **(d)** Detailed view of 3 GoF mutations highlighted in panel **b**. Quantitative binding model from the ProBound tool^74^ is shown under the sequence to contextualize the mutation. For each GoF mutation, the position and variant is indicated (*e.g.* 120G for GoF1), and the predicted affinity of the indicated TF for the WT and variant sequences are shown. **(e)** Coarse-grained epistatic effect (log_2_(observed/expected), where expected is taken as a multiplicative model) of disrupting pairs of TFBS. All binding sites with predicted normalized affinities >0.05 are considered. Binding site numbering and shading follows panel **b** (bottom). Nominally significant differences in observation vs. prediction (Wilcoxon test: p < 0.01) have their edges marked. Pairs where the expected multiplicative activity falls below our minP-only basal control are marked by an x. **(f)** Example quantification of epistatic effects for two pairs of disruptions from panel **e**. Related to **Fig. S7-S14**. Similar analyses to those shown in this figure for *Epas1*:chr17_10063 CRE are presented for the other four model CREs in **Fig. S9. (g)** Heatmap summarizing variant effect sizes within mapped singleton-functional TFBS for all possible substitutions (sub-rows) of all five CREs (rows). Color hues map to TFBS identity (key at bottom left), and color intensities to the absolute fractional change in activity relative to WT, normalized to the difference between WT activity and the minP-only baseline. **(h)** ‘Necessity scores’ of singleton-functional TFBSs of each CRE (rows), defined as the fraction of WT activity lost upon its disruption. Bar marks the median effect across singleton-functional TFBSs of each CRE. Notably, some CREs such as *Lama1*:chr17_7784 have a median TFBS necessity score >75%, meaning that the majority of TFBSs within the elements are critical for function. In contrast, *Gata4*:chr14_5729 has a median TFBS necessity score <40%. Examples of a weakly necessary site (Gata4/6 site, ♟) and a highly necessary site (Sox17 site, ♕) are marked above the *Epas1*:chr17_10063 CRE row in both panels **g** and **h**.

**Figure 4.**
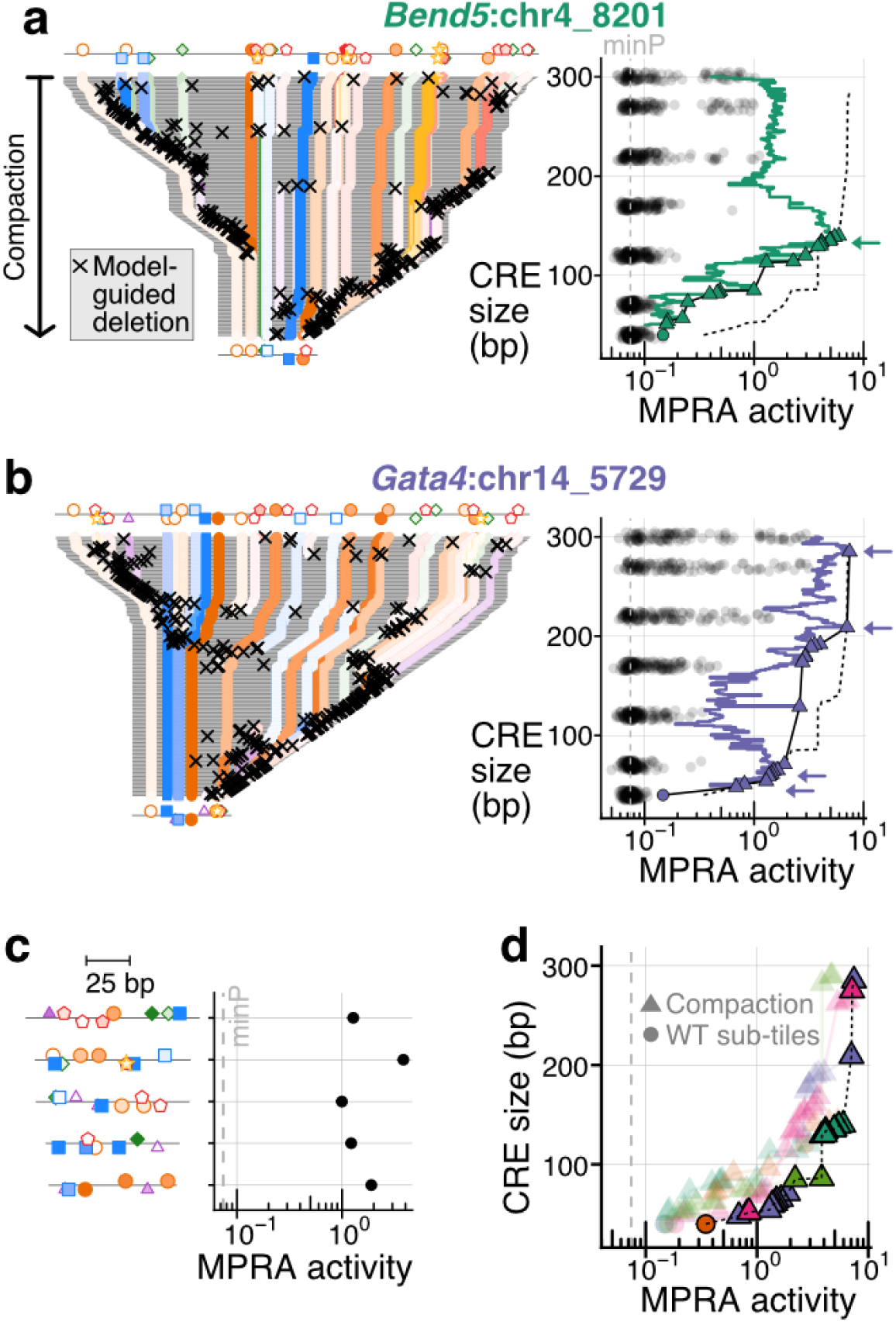
Model-guided compaction of CREs reveals the trade-off between activity vs. length. **(a)** Left: compaction trajectory for *Bend5*:chr4_8201, presented vertically from 300 bp (top row) → 40 bp (bottom row). Colors correspond to positions of mapped TFBS (normalized affinity >0.05). Black ‘x’ marks the model-selected base removed at each step. The starting (300 bp) and final (40 bp) sequences are duplicated above and below for reference. Right: MPRA activity (x-axis) as a function of CRE size (y-axis, aligned with sub-panel to left), plotting compaction trajectory measurements (green), endogenous sub-tiles from the same CRE (semi-transparent grey points; **Figs. 2**, **S4**; forward/reverse averaged; jittered for visualization), basal minP activity (vertical grey dashed line), and global Pareto front (black dashed line). Tiles lying along the CRE-specific Pareto front are marked (triangles: compacted elements; circles: endogenous sub-tiles) and connected by a black line. Tiles contributing to the global Pareto front are highlighted by green arrows. **(b)** Same as panel **a**, but for CRE *Gata4*:chr14_5729. See **Fig. S16** for all compaction trajectories in a similar format. **(c)** The most highly active sub-100-bp compacted tiles for each model CRE: *Epas1*:chr17_10063-Δ204bp, *Sparc*:chr11_7211-Δ214bp, *Bend5*:chr4_8201-Δ215bp, *Lama1*:chr17_7784-Δ226bp, *Gata4*:chr14_5729-Δ229bp). Vertical gray dashed line shows basal minP activity. For each model CRE’s compaction trajectory, we identify at least one sub-100 bp with activity >13-fold above basal minP. **(d)** Individual CRE-specific activity Pareto fronts are plotted (lightly colored, outline-free shapes). Triangles derive from compaction trajectory tiles and circles from wildtype (WT) sub-tiles. Points optimal along the activity-size tradeoff across all CREs are marked with black edges, and collectively delineate the global Pareto front (black dashed line), which is reproduced in panels **a-b**. Tiles from all CREs contribute to this global front (color scheme as in **Fig. S16a-b**). However, considering only >40 bp tiles (as 40 bp tiles are not represented in the final CRE compaction library; **Fig. S15b**), only a single WT tile contributes to any of the CRE-specific fronts.

Consistent with this, plotting activity as a function of distance (Δx) nearly quantitatively collapses the tiling profiles onto the same distance-decay curve (**Figs. 2f**, **S6a**). This short-range positional effect is quantifiable via the opening of a gap in a scatterplot comparing activity for the same tiles the forward (+) vs. reverse (-) orientation (**Fig. 2g**). Although smaller in magnitude, the same phenomenon was also observed at large tile sizes for each of the four other model CREs (**Figs. S6b**). This suggests that as a sub-tile is scanned along the region and gains activity upon acquiring active subsequences at its minP-proximal 3’ end, reverse complementing the sub-tile shifts those active subsequences to be distal from minP, dampening their contribution to regulatory activity.

Our findings suggest that without careful assessment of the placement of regulatory sequences, functional measurements made on larger sequences may be misleading. For example, our full CREs from the original screen in mouse EBs^40^ were selected to be ∼1 kb centered around ATAC peak summits. These full CREs sequences were included in our initial sub-tiling MPRA experiment (**Fig. S2**). While all displayed activities above basal minP-only controls, their activities were sometimes substantially weaker than the maximum activity 270 bp sub-tiles, possibly because of the positional effect reported here. Hence, although larger fragments of regulatory DNA intuitively come with increased potential for function, short-range positional effects might be strong contributors to variable activities as measured within conventional MPRAs.

However, we caution that this effect might be specific to our reporter architecture, which leverages a widely used basal promoter: minP from the pGal4 vector^17,56,66–70^. Whether these observations generalize to other minimal promoters (or endogenous promoters) will need to be assessed, but similar distance-dependent effects have been noted before in the context of synthetic regulatory elements^71,72^ and even used to tune expression explicitly^71,72^.

### Multi-hit saturation mutagenesis of active 300 bp regions

To gain insight into the distribution of effect sizes, as well as the diversity of mechanisms through which point mutations modulate the activity of endogenous developmental enhancers, we performed saturation mutagenesis of our model CREs. For this, we focused on a 300-bp region of each CRE, centered on the highest-activity 270-bp tile from the initial sub-tiling screen (**Fig. S2**). We designed, synthesized, and cloned a barcoded 5′ MPRA library enumerating all possible single-base substitutions of this region (1.6M barcodes; 99.9% of substitutions represented by >100 barcodes; **Fig. S7a-b**). Notably, mapping variants to barcodes revealed an average of 2.3 mutations per insert, with the number of additional mutations following a Poisson distribution (mean λ = 1.3), consistent with low-rate PCR mutagenesis^73^ during library construction (**Fig. S7c-e**). As a consequence, 92% of possible single-base deletions were also represented by ≥5 barcodes, providing coverage of deletion effects even though these were not part of the library design. Variant effect estimates were highly reproducible across biological replicates (R^2^ [on log-transformed activities] >0.65 across biological replicates; **Fig. S8a-b**), as well as when estimated from single-mutation vs. multi-mutation tiles (**Fig. S8c-d**). Together, these observations indicate that the dataset is well powered for quantitative dissection of mutational sensitivities in these developmental enhancers.

Mutagenized tiles were highly active, yet frequently and focally susceptible to point mutations. Wild-type (WT) tiles ranged from 2.0– to 36-fold above basal minP-only controls (bootstrap resampling p < 5⨉10^-4^ for all; **Fig. S8e**). For the four CREs with WT activity >2-fold above background, a high proportion of positions were mutation-sensitive (47-73% of bases with |effect| >1.25-fold & FDR < 0.006; **Figs. 3a-b**, **S9a-b**, **S9e-f**, **S9i-j**, **S9l-m**). Sensitive positions frequently formed contiguous tracts, consistent with coincident TFBSs. To link mutational effects to specific regulators, we mapped motifs for parietal endoderm *cis*-regulatory module (CRM) TFs using quantitative biophysical binding models^74^ (n = 20-37 putative CRM TFBSs per CRE). Of these, a substantial proportion (n = 8-15 per CRE) exhibited correlation between variant affinity and variant effect (some as high as R^2^=0.88; see examples in **Fig. 3c**; all sites across profiled CREs in **Fig. S10**), consistent with previous studies^7,16,17,39,75^. These ‘singleton-functional’ sites are all individually necessary for function by definition (disruption leads to decrease in activity) and cover a surprisingly high proportion of the profiled CREs (**Fig. 3g**). Still, they were individually insufficient to drive substantial activity on their own, in that none of the 40 bp tiles previously assayed bearing these sites drive substantial activity (**Figs. S4b**, **S11a**). Performing an unbiased categorization independent of the definition of these singleton-functional TFBSs, a striking 73% of loss-of-function (LoF) mutations were attributable to disruption of a parietal endoderm CRM TFBS (710/970 with log_2_FC < –0.32 → decrease of >0.075 in normalized affinity units, one-sided Wilcoxon test: p < 2⨉10^-16^; **Fig. S12**), validating our developmental TF nomination process (**Fig. S1c**) and underscoring the central role of dynamic regulators in the function of cell-type-specific enhancers.

Do simple biochemical features such as binding affinity dictate the importance of individual sites? Among all CRM TFs, high-affinity sites were more likely to be functional (Fisher’s exact p < 7⨉10^-4^), yet affinity itself only weakly predicted the magnitude of any binding site’s contribution towards activity (R^2^ = 0.14 between affinity and fractional contribution; **Fig. S11b**). Thus, these data implicate a subset of TFBS as critical contributors to enhancer function, while underscoring that biochemical binding strength is not the sole determinant of a given TFBS’s contribution to activity^76^.

Saturation mutagenesis data also revealed a surprising number of gain-of-function (GoF) substitutions^12,77^ (per CRE, n = 65-123 → >1.25-fold increase, and n = 3-15 → >2-fold increase; **Figs. 3a-b**, **3d**, **S9a-b**, **S9e-f**, **S9i-j**, **S9l-m**, **S13a**), suggesting that—at least with respect to the measured activity—the endogenous sequences of these CREs are far from a local optimum. With a loose threshold, roughly half of GoF mutations were predicted to substantially increase the affinity of at least one CRM TFBS (**Fig. S13a-b**; 205/413 with log_2_FC > 0.32 → increase of >0.075 in normalized affinity units; one-sided Wilcoxon test: p < 2⨉10^-16^). Conversely, ∼30% of mutations with large predicted affinity gains were GoF (**Fig. S13c-d**; 95/313 with increase of >0.25 in normalized affinity units → log_2_FC > 0.32; one-sided Wilcoxon test: p < 2⨉10^-16^). This strong but partial correspondence between TF affinity and enhancer activity confirms the importance of DNA-proximal biochemical events for regulatory function while also underscoring the complexity of the affinity-activity relationship, and that our set of CRM TFs likely misses some regulators of these model enhancers.

With these quantitative activity measurements together with biophysical estimates of TFBS affinity, we sought to assess the ‘distribution of mechanisms’ underlying large-effect GoF substitutions (log_2_FC > 0.5; n = 210). Around 20% appear unrelated to CRM TFs and we speculate are related to missed regulators (n = 45; <0.02 change in normalized affinity). Of the ∼80% associated with a change in CRM TFBS affinity, we delineate three categories (**Fig. S13f-g**): (i) changes strengthening an already-functional TFBS (n=40, singleton-functional with starting normalized affinity ≥ 0.05); (ii) changes strengthening an existing but sub-functional TFBS (n=37, non-singleton-functional sites with starting normalized affinity ≥ 0.05); and (iii) changes that create a new TFBS (n=88, starting normalized affinity < 0.05). For example, in the *Epas1*:chr17_10063 CRE, GoF1 and GoF2 increase the predicted affinity of predicted functional Klf4 and AP-1 TFBSs respectively (class i), while GoF3 creates a strong Sox17 TFBS from a non-functional low-affinity pseudosite (class iii) (**Fig. 3d**). Additional examples illustrate both ‘continuous’ affinity-maturation (classes i and ii) and ‘discontinuous’ TFBS creation (class iii) (**Fig. S9b,f,j**; detailed in **Fig. S13e**). Notably, these categories were associated with similar distributions of effect sizes (**Fig. S13g**). Together, these results show that both affinity maturation and *de novo* TFBS creation represent readily accessible paths for increasing the activity of endogenous enhancers.

The five model CREs were also differentially robust to mutations, and relatedly, the functional necessity of individual TFBS within CREs varied greatly. At one extreme, CRE *Lama1*:chr17_7784 was remarkably fragile, harbouring 8 distinct TFBSs associated with >75% LoF if individually disrupted (**Figs. 3h**, **S11c**). More broadly, 33 mutations modulated its activity by >4-fold (**Figs. S9f**, **S11d**). In sharp contrast, CRE *Gata4*:chr14_5729 was highly robust: none of its CRM TFBS was associated with >75% LoF, and only 1 mutation over the entire CRE modulated activity by >4-fold (**Figs. 3h**, **S9b**, **S11c-d**).

Does this differential robustness have its roots in differential buffering via epistatic interactions? The multi-mutation aspect of our saturation mutagenesis data, due to the superimposition of programmed substitutions and PCR errors (**Fig. S7c**), positions us to ask this long-standing question^17,39,75,78^. While library complexity was insufficient to assess pairwise epistasis at base-pair resolution, coarse-graining provided enough coverage at the level of TFBSs (**Fig. S14**). Examining all pairs of TFBS and comparing to a null multiplicative expectation^79^, we mapped CRE-dependent epistasis landscapes and identified rare but clear examples of both synergistic and buffering interactions (respectively, instances where paired disruption was significantly worse or better than the product of individual disruptions; **Figs. 3e-f**, **S9c-d**, **S9g-h**, **S9k**). Binding-site-level epistasis was uncommon, except for the *Lama1*:chr17_778 CRE. However, the pervasive >0 epistatic ratio (**Fig. S9g**) in that element was a reflection of its fragility: most constituent binding sites were fully necessary for function (**Fig. 3k**) such that pairwise disruptions cannot bring activity lower than baseline. Intriguingly, the robust *Gata4*:chr14_5729 did not display buffering, *i.e.* pairwise perturbations of it were generally no worse than the multiplicative expectation except in two isolated examples (**Fig. S9c-d**), suggesting that the mutational tolerance of this enhancer goes beyond second-order interactions.

In summary, our multi-hit mutational scanning of these endogenous developmental CREs provided a high-resolution, quantitative view of regulatory function, with the following insights. <u>First</u>, profiled elements, selected for their cell-type specificity, harboured a remarkably high density (approximately one per 20 bp) of functional TFBSs for the orthogonally nominated parietal endoderm CRM TFs (**Fig. 3g**). <u>Second</u>, distinct TFBSs had drastically different functional importance across CREs in a pattern that was only weakly encoded by their individual affinity (**Fig. S11b**). <u>Third</u>, despite having similar activity, distinct CREs were differentially tolerant to mutations, and our small set of model CREs included clear examples of both fragile and robust enhancers (**Fig. 3h**). <u>Fourth</u>, GoF mutations were common, and detailed analysis revealed creation of a new TFBS vs. affinity maturation of an existing TFBS as roughly equiprobable paths to activity gains (**Fig. S13g-g**). <u>Fifth</u>, systematic assessment of pairwise TFBS disruptions demonstrated that second-order epistasis was rare, at least at the level of resolution allowed by our library (**Figs. 3e**, **S9c**, **S9g**, **S9k**). <u>Finally</u> and perhaps most importantly, mutagenesis underscored the ‘individuality’ of each regulatory element.

### Delineating the activity-size regulatory Pareto front with model-guided enhancer compaction

Although a high proportion of base-pairs within our model CREs were functional, 1-bp deletions commonly exhibited modest deleterious effects (102-141 1-bp deletions [supported by ≥5 BCs] per 300-bp CRE with log_2_FC > –0.32). This raised the possibility that tolerated deletions can be combined without compromising activity. Given that TFBS density is a key biochemical parameter^60^, such compaction might even increase activity. Pushing this further, we might ask, how far are endogenous enhancers from *maximal compaction*?

To investigate this, we framed a two-objective optimization problem in which Pareto front defines the optimal trade-off between activity and sequence length^80^. Our prior high-density sub-tiling experiments sampled part of this space but was limited to contiguous fragments of the native sequence (**Fig. 2b**), and therefore could not reveal how far wild-type CREs lie from the true front. We developed a model-guided strategy for *in silico* compaction by training a chromBPNet^19^ model to predict parietal endoderm-specific chromatin accessibility^40,53^. Then, starting from the 300-bp endogenous tiles, we predicted accessibility for all possible 1-bp deletions, selected the one with the highest predicted accessibility, and iterated until the tile was compacted to 40 bp. Model-selected deletions were enriched near sequence edges, and extended regions lacking singleton-functional TFBSs frequently removed in contiguous blocks (**Figs. 4a**, **S16a**).

This *in silico* procedure yielded trajectories compacting each CRE from 300 bp → 40 bp (**Figs. 4a-b**, **S16a**). We then synthesized these sequences and measured their activities via MPRA in PYS-2 cells. Of note, cloning biased the library towards shorter elements (**Fig. S15a-b**; median BC: all elements, 94; 46-80 bp elements, 287; 260-300 bp elements, 30; 40-45 bp sequences were lost due to overly stringent size selection). Nonetheless, replicate concordance was very strong (R^2^ [on log-transformed values] > 0.98; **Fig. S15c-d**).

Strikingly, model-guided compaction yielded many sequences that maintained or exceeded the activities of their 300-bp parent sequence. Across the five model CREs, the shortest sequences with >WT activity ranged from 82-191 bp (**Figs. 4a-b**, **S16b**). Moreover, when compared to endogenous tiles of matched length, compacted sequences overwhelmingly exhibited higher activities (**Figs. 4a-b**, **S16b**). Aggregating trajectories across CREs allowed us to approximate the global activity-size Pareto front (dashed lines, **Fig. 4e**). Although this front exhibits a sharp drop in activity below ∼75 bp, all trajectories harbored sub-100-bp compacted CREs with >13-fold activity over the basal minP control (**Fig. 4f**).

ChromBPNet^19^ predictions were correlated with MPRA activity for 3 of 5 compaction curves (**Fig. S16c-d**). However, as we only tested one compaction curve per CRE and all five trended upwards, we cannot draw strong conclusions about the efficacy of model guidance vs. a scenario in which deletions were introduced in a random order. Empirical compaction curves were much more rugged than their *in silico* counterparts, with many sharp changes caused by individual deletions (**Fig. 4b,d**; not attributable to technical noise: **Fig. S15d**). Model-compacted sequences harboured a higher density of CRM TF binding sites compared to random *in silico* deletion trajectories or endogenous sub-tiles (**Fig. S16e**), suggesting the model implicitly prioritizes this feature.

Together, these results show that endogenous enhancers are typically far from optimally compact. All CREs contributed sequences to the aggregated front, suggesting compaction from different functional starting points efficiently explores the sequence space. Thus, repeating the process starting from more CREs might converge on the true regulatory trade-off, especially if using function-focused predictive models^81,82^. Compaction also represents a complementary strategy to *de novo* design of compact synthetic enhancers^4,24–26,83^.

### Enhancer derivatization empirically tests the global stringency of cis-regulatory grammar

Mutagenesis identifies ‘stiff’ and ‘soft’ directions in local regulatory sequence space: ablating a TFBS can abrogate function (**Fig. 3**), but removing multiple ‘neutral’ bases between TFBSs is often tolerated (**Fig. 4**). What about changes that explore larger leaps in the sequence space? Despite a vast literature on *cis*-regulatory mechanisms and predictive modeling (reviewed in refs^76,84,85^), empirical assessments of the stringency of regulatory function are restricted to a handful of strategies corresponding to either very short radii (*e.g.* saturation mutagenesis^38^ or TFBS ablation, both effectively local variation around wildtype) or extremely large radii (*e.g. de novo*-designed synthetic CREs^4,24–26,83^ or random sequences^6,10^). Inspired by a vision set forth by DePace and colleagues^86^ and by random regulatory assemblies^87,88^, we instantiated an alternative approach: ‘enhancer derivatization’, which aims to explore the vast sequence space that lies between these two extremes in a structured manner.

Enhancer derivatization involves applying simple TFBS-anchored sequence transformations to endogenous CREs. Although many transformations are conceivable, we explore three here: ‘reconstitution’, ‘random deposition’ and ‘synthetic thripsis’ (**Fig. 1c**). We first mapped TFBS for four parietal endoderm CRM TFs (Gata4/6, Sox17, Foxa2, Klf4; 9-18 TFBS per model CRE) and merged overlaps, resulting in 8-10 indivisible ‘TFBS anchors’ per CRE, which summed to 103-159 bp per 300 bp element (**Fig. S17**). We then designed 18,405 anchor-retaining derivatives, corresponding to reconstitution (n = 10,000), random deposition (n = 3,000), and synthetic thripsis (n = 3,000), together with negative controls (n = 2,400) and WT sequences (n = 5) (**Fig. S18a-b**). Following synthesis and cloning (**Fig. S18c**; total 1.89M BCs; median 91 BCs per CRE), we subjected these to bulk MPRA in PYS-2 cells. Activity measurements were reproducible and spanned three orders of magnitude (**Fig. S18d**; R^2^ [on log-transformed activities] ≥ 0.94 between biological replicates). Negative controls overwhelmingly exhibited low activity, while many derivatives displayed substantial activity (**Fig. S18f**; 32%-84% per model CRE with activity >3-fold above minP).

<u>Reconstitution</u> involves introducing the TFBS anchors to a limited number of non-overlapping positions within a larger number of background DNA sequences (**Fig. 1c**). In brief, we designed and characterized 2,000 reconstitution derivatives per model CRE: 5 position sets for the TFBS anchors (3 endogenous with varying TFBS flank retention, 2 randomized with WT orientation) × 400 backgrounds (**Fig. 5a**). The 400 background sequences included 200 inaccessible endogenous sequences and 200 dinucleotide shuffled sequences, and were reused across the 5 model CREs × 5 position sets. Reconstituted derivatives broadly activated expression, but in an anchor set and background-specific manner (**Figs. S19a, S20**). TFBS anchors from CREs *Lama1*:chr17_7784 and *Gata4*:chr14_5729 were particularly potent (median 19.5 and 17.1 fold above minP, respectively). Strikingly, despite background sequences having little autonomous activity, they contributed substantially to variance, with the range of most position sets across 400 backgrounds exceeding the >100-fold range separating minP and the WT sequence (**Figs. 5b-c**, **S19b-c,** average span: 150-fold, mean 47% of log-activity variance explained by background identity for fixed TFBS sets). Across reconstitutions, certain backgrounds were broadly permissive, or on the contrary, broadly recalcitrant (**Fig. S19d**). Beyond such examples, the responses of a given background to different reconstitutions were weakly correlated (median R^2^ = 0.05 on log-transformed activities; **Fig. S19e-f**). Of note, we could not identify simple properties of background sequences that predicted their responsiveness, *e.g.* CRM TFBS present in background sequences were not explanatory (**Fig. S19g**), while TFBS flanking bases^5,89^ were only modestly relevant (**Fig. S21**). Surprisingly, for two model CREs (*Lama1*:chr17_7784, *Sparc*:chr11_7211), random TFBS anchor location sets exhibited significantly higher mean activity than the endogenous location set (**Fig. S19a**), foreshadowing the next transformation operation. Altogether, we conclude that background DNA plays a large and complex role in shaping endogenous CRE activity, as adding 9-18 CRM TFBS at fixed positions –-endogenous or otherwise, and collectively covering one-third to one-half of the sequence –– is insufficient to yield consistent levels of activity across a range of backgrounds.

**Figure 5.**
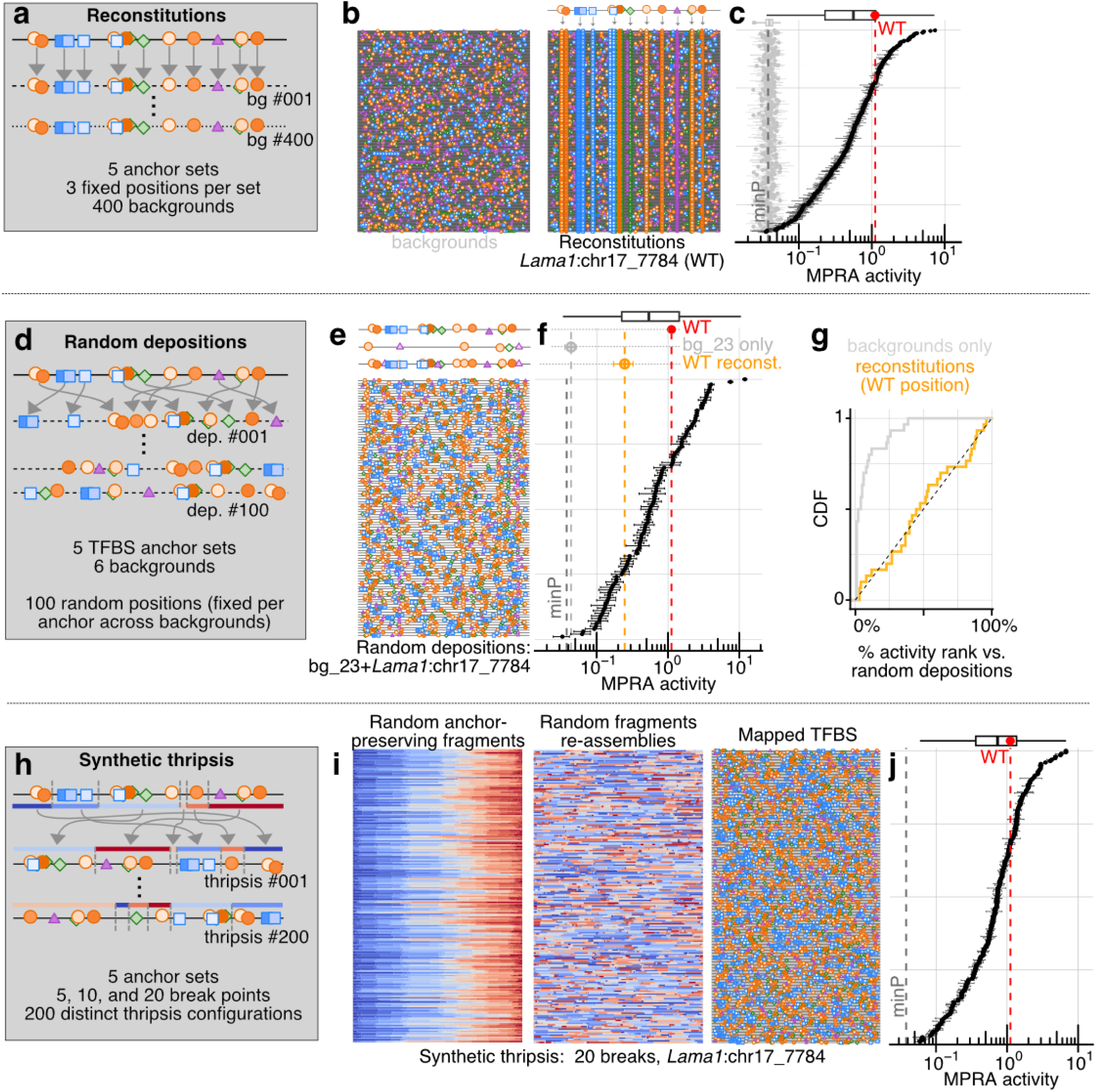
Enhancer derivatization empirically tests the global stringency of *cis*-regulatory grammar. TFBS anchors were selected from each model CRE and re-arranged according to three types of transformations: reconstitutions, random depositions, and synthetic thripsis. For reconstitutions (panels **a-c**), random depositions (panels **d-g**) and thripsis (panels **h-j**), respectively 1/15, 1/30, and 1/15 of the data is shown. **(a)** Schematic of reconstitution (one anchor set shown): TFBS anchors from 5 model CREs are deposited to 3 fixed position sets across 400 background sequences. **(b)** TFBS (norm. affinity > 0.1) for the four anchor CRM TFs (Gata4/6, Sox17, Foxa2, Klf4) mapped to background sequences alone (left) and following reconstitution with *Lama1*:chr17_7784 anchors at WT positions (right). Only non-autonomously active backgrounds are shown (n = 347/400, activity < 1.5× minP), ordered by activity. **(c)** MPRA activity of background sequences (grey) and Lama1:chr17_7784 WT-position reconstitutions (black), aligned with the TFBS schematics in panel **b**. Each point is a single sequence (error bars: s.d. across replicates); box plots summarize each distribution (line: median; box: IQR; whiskers: 1.5× IQR). Dashed lines indicate minP (grey) and the full WT CRE (red). **(d)** Schematic of random deposition (one CRE shown): TFBS anchors from 5 model CRE are placed at 100 randomized, anchor-collision-free position sets (fixed across backgrounds) on each of 6 background sequences. **(e)** TFBS (norm. affinity > 0.1) anchor CRM TFs mapped onto (from top to bottom): full WT *Lama1*:chr17_7784 CRE, background bg_23 alone, WT-position deposition, and 100 random depositions of *Lama1*:chr17_7784 TFBS anchors onto bg_23. Depositions are by activity. **(f)** MPRA activity of the 3 reference configurations and 100 random depositions shown in panel **e** aligned with the corresponding TFBS schematics. Full WT CRE (red), bg_23 background (light gray), WT-position deposition (orange), and minP (dark gray) activities are highlighted with vertical dashed lines as well for reference. **(g)** Cumulative distribution of the activity rank of WT-position reconstitutions (orange) and background-only controls (grey) relative to the 100 randomized depositions on the same background across the 5 CRE x 6 background = 30 deposition sets. The dashed diagonal represents the expectation if WT positions conferred no advantage over random positions (observed median rank = 46%). **(h)** Schematic of synthetic thripsis (one model CRE and break-point density shown): WT sequence of 5 model CREs are fragmented at randomly selected, anchor-avoiding break points (5, 10, or 20) and the resulting fragments are reassembled in random, orientation-preserving orders (200 per anchor set). **(i)** Illustration of thripsis procedure (n = 20 breakpoints) applied to CRE Lama1:chr17_7784. Fragments are colored from blue (5′) to red (3′) based on their position in the starting CRE. Shown are the anchor-preserving fragment selections (left), their randomly reassembled configurations (middle), and the resulting CREs with mapped TFBS (right). **(j)** MPRA activity of the synthetic thripsis elements shown in panel **i**. Dashed lines and box plot as in panel **c.**

<u>Random deposition</u> involves introducing the TFBS anchors to a large number of position sets of a limited number of background sequences (**Fig. 1c**). In brief, we designed and characterized 600 random deposition derivatives per model CRE: 100 position sets per TFBS anchors (randomized but fixed, WT orientation) × 6 backgrounds (**Fig. 5d-g**). Similar to the reconstituted derivatives, random deposition derivatives exhibited widely ranging activities (*e.g.* **Figs. 5e-f**, **S22a, S23**), TFBS anchor sets from *Lama1*:chr17_7784 and *Gata4*:chr14_5729 again were particularly potent, and TFBS anchor identities explained 52% of the log-activity variance. Still, even on a fixed background with a fixed set of TFBS anchors being deposited, the activities spanned, on average, a 64-fold range across position sets (**Fig. 22a-b**). For a given model CRE, background DNA and position set explained, on average, only 19% and 18% of the log-activity variance, respectively, pointing to complex determinants of activity. Indeed, the correlation of random deposition activities across different backgrounds was essentially null (median R^2^ = 0.007, **Fig. S22c-e**), suggesting strong interactions between deposited TFBS and background sequence features. As in the reconstitution data, the sequence determinants of these interactions appear non-trivial, *e.g.* the slight variations in final TFBS numbers on the resulting CREs arising from anchor/background chimeric junctions explain <20% of the log-activity variance (**Fig. S22f-g**). As the six selected background DNA sequences used in random depositions here were included in the reconstitution set, we could conduct a controlled comparison of the activity of CREs with randomized vs. WT anchor site positions deposited on the same DNA sequence. Across backgrounds and anchor sets, WT position sets (orange cross, **Fig. 5f, S22b**) spanned the full activity range of randomized position sets (median quantile = 0.46, **Fig. 5g, S22b**). Hence, the endogenous positions of CRM TFBS do not maximize activity, and are not even favoured statistically compared to randomized position sets. Overall, systematic measurements of random deposition CREs across backgrounds and different anchor sets confirm the dramatic functional consequences of higher-order TFBS arrangements.

Finally, inspired by cancer genome chromothripsis^90^, <u>synthetic thripsis</u> involves the *in silico* shattering and reassembly of an endogenous CRE, thereby minimizing the alteration of overall sequence composition while reorganizing the element globally (**Fig. 1c**). In brief, we designed and characterized 600 synthetic thripsis derivatives per model CRE: 3 breakage densities (5, 10 or 20 breakpoints) × 200 configurations that preserve TFBS anchor sequences and their relative orientations (**Fig. 5g-j**). Quantitative analysis of gapped k-mer compositions^91,92^ for thriptic CREs compared to WT indeed confirmed the very high similarity in sequence space for this transformation compared to others (**Fig. S18e**). Overall, thripsized derivatives exhibited a wide range of activities (average max/min span 83-fold; **Figs. 5j**, **S24a**, **S25**) that are generally higher than reconstitution or random deposition derivatives, even at a high breakage density (**Figs. S18f**, **S24b**). Is this because thripsis preserves more of the higher-order structure among TFBS anchors, creates new TFBS at thripsis junctions, or preserves sequence features beyond the TFBS anchors? Stratifying thripsis derivatives by preservation of linkage between consecutive pairs of TFBS anchors found only 1/39 consecutive pairs to be significantly predictive of activity (**Fig. S24c-d**). Hence, although local pairwise interactions <u>can</u> be highly constraining (*e.g.* the Foxa2-Sox17-Gata4 triplet of the *Epas1* CRE; **Fig. 3a**), this is uncommon (with tne caveat that some anchors were composed of multiple TFBS). The creation of new TFBS at thripsis junctions was also uncommon and poorly predictive (<13% of log-activity variance explained; **Fig. S24f-g**). In contrast, disruption of functional sites not included in our TFBS anchors (*e.g.* the AP-1 sites highlighted in **Fig. S17b**) by breakpoints was broadly correlated with significant reductions in activity (**Fig. S24e**). This suggests that the generally higher activity of synthetic thripsis derivatives, relative to reconstitution and random deposition derivatives, is best explained by this transformation’s greater likelihood of preserving additional non-anchor TFBS present in the endogenous enhancer sequence. Overall, our results with synthetic thripsis show that: (i) the collective organization of TFBS, including and beyond key CRM TFBS, is critical for activity, and (ii) strong pairwise functional linkage is the exception rather than the rule.

## DISCUSSION

Multiple qualitative views have been proposed to describe the cell type-specific *cis*-regulatory grammar of metazoans, *i.e.* how the relative positions, orientations, number, affinities, and interactions among TFBSs influence activity. Famously, at the two extremes, models include complete flexibility (‘billboard’^93^, ‘bag-of-motif’^37^) vs. absolute rigidity (‘enhanceosome’^94^), with other views falling in between (‘TF collectives’^15,95^). Adding quantitative nuance to these conceptual frameworks, recent work leveraging deep learning models^4,5,96^ suggest the existence of non-strict but nevertheless consequential preferences in TFBS arrangements (‘soft syntax’). Still, the implications of these preferences for the functional robustness of CREs to sequence perturbations remains unresolved. Previous studies leveraging deep learning predominantly focused on pairwise effects for *in silico* explorations of TF-to-TF synergies^4,96^. However, our data on short 40 bp tiles suggests that a cluster of two TFBS is typically insufficient to generate function, at least as measured within our reporter system. Hence, TFBS pairs probably need to be embedded within a richer set of TFBS to productively lead to mRNA production. As such, higher-order contextual information of other TFBS cannot be abstracted out, and querying pairs of TFBS is unlikely to be a sufficient level of complexity to truly deconstruct regulatory function. Indeed, our tiling indeed suggests activity is the result of extensive nonlinear interactions between a much larger number of sequence features (*i.e.* with high activity typically arising from ∼8-15 TFBS within the enhancers we profiled).

Within this context, we sought to obtain a richer view of the stringency of the regulatory code, beyond saturation mutagenesis and pairwise interactions, but still centered on key TFBSs. This led us to the various derivatization operations, in which the endogenous architectures of a small number of cell type-specific enhancers were subjected to a large number of fragmentation, compaction, mutation, or reassembly operations, substantially altering CREs but in an ‘anchor preserving’ manner. The approach taken here provides complementary insights to those described in a companion study, in which evolutionary and synthetic mutational trajectories of these same five developmental enhancers were deeply explored^53^.

The activities of enhancer derivatives do not align with predictions of full flexibility or rigidity (respectively all derivative activities at WT or zero). Rather, derivatives display ∼100-fold activity variation within samplings of same-class sequence transformations, and the underlying sequence determinants are complex. As a specific case of variability (consistent with previous large datasets of synthetic enhancers employing TFBS depositions^27^), we find that ‘background DNA’ is a surprisingly strong contributor to activity, even when it does not harbor apparent binding sites to the key lineage-defining TFBSs for the cell type in question. This is probably reflective of the vast repertoire of active TFs within any given cell type, many of which are not cell-type-specific but nonetheless may serve as key cofactors (*e.g.* AP-1). However, there may be other ways in which background sequence modulates activity, *e.g.* DNA shape characteristics^97^ or bendability^98^.

Beyond derivatives, the different classes of sequences we profiled here uncover a rich set of quantitative regulatory phenomena which warrant further mechanistic dissection in the future. <u>First</u>, what is the molecular basis of the strong non-additive effects between sub-fragments seen across all model enhancers profiled? Functionally, this effect reminiscent of an ‘AND’ gate^99^ on recruited TFs, and many (non-exclusive) explanations are conceivable: enzymatic action from different TFs at different limiting steps of the transcription cycle (‘kinetic control’)^100^, nucleosome mediated cooperativity^101^, direct TF-to-TF interactions^99^, transcriptional condensates^60^, etc. Compiling a comprehensive set of such high-synergy tile pairs genome-wide could provide discriminating information to disentangle mechanisms of combinatorial regulatory control. <u>Second</u>, what drives the large short-range positional effects that we see within minP-based reporter assays? Possible explanations include differential stability of the resulting reporter mRNA if substantial transcription initiation happens within the enhancer, or nucleosome invasion^102^ of the promoter region upon moving the active sequence further out. <u>Third</u>, we find a remarkably emergent simplicity in the variant effects of many TFBS within our mutagenesis data, namely a strong linear relationship in a log-log plot between predicted affinity and observed activity, with the slope of the line differing across TFBSs (**Figs. 3c, S10**). How do these simple relationships, which also seem to validate affinity models, coexist with the higher order dependencies among TFBSs? What are the determinants of the observed susceptibility slopes?

Given the recent improvement in predictive performance of sequence-to-function models^4–7^ and successes in difficult tasks such as *de novo* enhancer design^23–27^, the main open modeling challenges for the field, in our view, consist of learning, predicting and designing higher-level properties for regulatory DNA. One such avenue could be exploration of the nature of fundamental trade-offs between possibly opposed features of enhancers. A <u>first</u> example of such opposed features explored here is CRE length vs. activity level. Using both multi-sized sub-tiling and model-compacted enhancers, we find a preliminary delimitation in the quantitative shape of the activity-by-size Pareto front. But what is the fundamental limit? Furthermore, how is the shape of the Pareto front modulated by the requirement of specificity? Indeed, a <u>second</u> natural trade-off is between activity and specificity. A large number of gain-of-function mutations are found within our model CREs, but it remains to be tested which of these variant elements, if any, also lose specificity (each model CRE dissected here having been selected as highly specific to parietal endoderm vs. pluripotent progenitors). Previous reports have found competing objectives between activity and specificity, especially for gain-of-affinity variants^103–106^, but whether this trade-off is unavoidable in general remains to be determined. A <u>third</u> intriguing trade-off is between activity and genetic robustness. Or are these two properties entirely orthogonal? Like activity, the robustness of enhancers to genetic mutations is a complex result of sequence-level features, which should presumably be tunable. Empirically, we observe different tolerance to mutations within endogenous developmental elements of similar activity, but can such properties be synthetically programmed in an activity-preserving way? If so, what are the limits of such robustness? Can one design an enhancer that is perfectly insensitive to point mutations? In our opinion, focusing on such complex and challenging design tasks holds epistemological value, and the potential to yield generalizable, actionable information about the regulatory code.

Our study has several limitations. <u>First</u>, like most high-throughput functional enhancer assays, we leverage episomal assays, such that differences in chromatinization could mean that our findings might not translate when contextualized within the genome, particularly for short-range positional effects. <u>Second</u>, we focus on activity predominantly within a single static cell line (PYS-2, a surrogate for parietal endoderm), whereas true biological activity of developmental enhancers is within multicellular contexts, and regulatory function is thus inherently multi-functional. <u>Third</u>, we used discrete molecular categorizations for our gain-of-function mutations (*e.g.* definition of affinity maturation vs. creation of new TFBS) even though affinity is a nearly continuous variable. However, the underlying biophysical binding curve in effect implements a nonlinear function, leading to sites with drastically different TF occupancies based on their predicted affinity. We used 5% of our normalized affinity as a reasonable heuristic cutoff for our definition, with the understanding that optimal threshold could have been set on a TF-by-TF basis based on their determined expression levels. <u>Fourth</u>, our pairwise epistatic mapping was limited to coarse-grained view at the level of full TFBSs, thereby obscuring a possibly rich landscape of interactions between individual mutations. Comprehensive, higher resolution epistatic mapping might reveal more pervasive TF-to-TF interactions. <u>Fifth</u>, our set of selected TFBS anchors for the derivatization procedure did not include the critical AP-1 factors, and possibly other important TFs, whose roles within our enhancers are mapped contemporaneously to the derivatization designs. An updated scheme including these important TFBSs might have revealed less variability in derivatives’ activities (although our anchors already covered 34-53% of the CRE sequence). <u>Sixth</u>, our definition of singleton-functional TFBS (looking for correlation between affinity and activity variation within a site) fails in cases where there is overlapping TFBS (breaking the correlation). This would show up as not reliably attributable to a given TF, hindering our sequence-level interpretation (*e.g.* the AP-1 cluster within CRE *Epas1*:chr17_10063). <u>Finally</u>, we see the most serious limitation of our work as the lack of clear sequence-level interpretation or predictive power in attributing the activity of our CRE derivatives and other libraries, such that integration of state-of-the-art predictive modeling architectures is a key next step. Of note, previous efforts in developmental systems, as well as our results here, underscore the difficulty of this task (*e.g.* impact of background DNA^27^). More generally, while we have assessed robustness of regulatory activity with derivatization, it remains a formidable challenge to extrapolate design rules from these results. Even though tested CREs are constrained in sequence space, explicit featurizations remain too high-dimensional.

In the end, this study exemplifies empirically what makes the *cis*-regulatory code challenging to resolve: stiff and soft directions in a rugged landscape, massive deviations from simple views of grammar, and multiple high-order interactions between TFBS collectively leading to biological activity. A clear lesson is that to truly understand the regulatory code, training predictive models must gather more sequences than endogenous ones^10^. We propose that enhancer derivatives, obtained from simple sequence transformations, explore useful regions of sequence space. Given their trivial stochastic generative nature, the resulting sequences could provide a scalable path towards data augmentation^107^ for the next generation of machine learning models in regulatory genomics. In the present moment, a key opportunity for the field may be to shift focus away from diminishing returns in predictive power in specific cell lines, and towards the inference of generalizable principles for endogenous gene regulation in developmental contexts.

## AUTHOR INFORMATION

## Acknowledgments

We thank the Shendure lab ‘gene regulation’ subgroup members, particularly F. Abadie, D. Calderon, W. Chen, and T. McDiarmid for extensive discussions that have shaped the direction of the project. We also thank all members of the Shendure lab for critical inputs. We acknowledge N. Ahituv, M. Kircher and their team for critical comments. We thank Jin Woo Oh for sharing and adapting a script to generate gapped k-mer composition vectors^91^. This work was supported by the National Human Genome Research Institute (UM1HG011966 to J.S.), the Brotman Baty Institute for Precision Medicine and the Seattle Hub for Synthetic Biology, a collaboration between the Allen Institute, Chan Zuckerberg Initiative and University of Washington (award number CZIF2023-008738 to J.S.). J.-B.L. was supported by a Damon Runyon Cancer Research Foundation fellowship (DRG-2435-21) and a Next-Generation Scientist award from the Cancer Research Society (grant no. 1155581). J.S. is an Investigator of the Howard Hughes Medical Institute.

## Author Contributions

J.-B.L. and T.L. conceived of the project, designed/cloned complex libraries, and performed most MPRA experiments. E.K. and T.L. performed compaction MPRA. C.H. and J.-B.L. performed sub-tile MPRA in mESC. T.D. performed some sub-tiling MPRA experiments with T.L. B.K.M. provided guidance with MPRA reporter cloning and construction and assisted with data curation/deposition. S.R. provided expert guidance and data on early mouse developmental models. J.-B.L. and T.L. analyzed all data and generated figures. J.-B.L. and J.S. wrote the manuscript with input from T.L. J.S. supervised the study.

## Data and Code Availability

Raw and processed sequencing data generated in this study have been deposited and are freely available, including MPRA data (IGVF portal https://data.igvf.org/, accession numbers IGVFDS8273CGGA, IGVFDS3736EOJH, IGVFDS0841KEKZ, IGVFDS7673WNAA, IGVFDS3857FNVD, and IGVFDS2523LIBG), custom sequencing amplicon data for CRE-barcode associations (GEO, accession number GSE328309). Other used datasets included scRNA-seq (accession GSE217686) and scATAC-seq (accession GSE217683) data from mouse embryoid bodies^40^. Parameters from quantitative binding models for various TFs were used from MotifCentral^74^ (FOXA2: 17084, GATA4: 16735, GATA6: 17095, HNF1B: 12948, KLF4: 17150, SOX17: 10629, JUN:ATF3 [AP-1]: 15946). Amplicon & plasmid DNA maps, and activity tables have been deposited on Zenodo (10.5281/zenodo.19658455). Core scripts for data analysis can be found on github: https://github.com/shendurelab/mammalian-cre-evolution-and-design.

## AI Disclosure Statement

We disclose that language editing, proofreading and coding were supported by AI-based tools; these were not used for conceptual development or primary manuscript writing. The authors take full responsibility for the contents of this manuscript.

## METHODS

### Nomination of TF for the parietal endoderm cis-regulatory module

To nominate transcription factors putatively implicated in gene regulation in the parietal endoderm lineage, we considered a criterion of differential expression coupled with differential accessibility of their associated motifs^108^. Our expectation was that TF with both induced expression and whose motifs were more highly accessible were good candidates as drivers of the lineage specific regulatory program. We used existing data from our previous scRNA-seq and scATAC-seq in mouse embryoid bodies^40^ (accession GSE217686 and GSE217683 respectively). Briefly, we computed pseudobulk RNA expression from the gene by cell count matrix, taking the mean UMI count per cell as measure of expression. For motif accessibility score, we used function peakAnnoEnrichment from ArchR^109^ (version 1.0.1) which performs a hypergeometic enrichment test of representation within marker peaks from different clusters, using cutoff of FDR≤0.05 and log_2_FC≥1. We used a custom set of position weight matrix combining Hocomoco^110^ v11 (both mouse and human), JASPAR^111^ 2018 and those reported in ref.^112^. Starting from a set of TFBS used in prior analyses^113^ (see: https://www.vierstra.org/resources/motif_clustering), we extracted mouse gene names (from one-to-one orthologs when dealing with human names) associated with each motif. RNA expression was then obtained from the associated gene, and motif accessibility from the unique motif position weight matrix identifier. Enrichment scores were averaged over all motifs associated to the same gene to enable a simple match to expression values. In cases for which TFs were associated with motifs from different similarity clusters, the maximum enrichment motif was taken. Fold-change in RNA expression in parietal endoderm (parietal endoderm divided by median over all clusters) was then plotted vs. motif enrichment score (**Fig. S1c**), revealing a small set of TFs which we take to be putative cis-regulatory module TFs in that cell-type (**Fig. S1d**).

### TF binding site mapping using ProBound

Reliable mapping of TFBS together with the change in affinity resulting from mutation is a key segment of CRE interpretation. Here, we used the ProBound^74^ tool. Briefly, we created a wrapper script that breaks down any given input sequence into stride-1 consecutive k-mers (size matched to the binding model of the TF of interest), computes the affinity of both orientations for all k-mers using: `loadMotifCentralModel(TF_ID).buildConsensusModel().addNScoring().inputTXT(seq_IN).bindingModeScores(/dev/stdout,profile)`

The wrapper then puts the information back in the original CRE position reference. This results in a 1-bp resolution affinity maps for a given TF binding model. To avoid binding model length drastically leading to different scales, we normalized the output affinity by the 99.99th quantile from a 1M random k-mer sampling (of note, with this definition, in some rare cases, the affinity of a select sequence could exceed 1). We applied a heuristic cutoff of 0.05 normalized affinity as putative TFBS to explore further (e.g., analysis of **Fig. 3c**). Orientation for TFBS was taken as the maximum-affinity orientation. Given the redundancy/similarity between Gata4 and Gata6 TFBSs, affinity was scored for both models. We grouped the binding affinities of these two TFs under the label Gata4/6, using the maximal normalized binding affinity observed for each (and reversing the TFBS Gata6 orientation if called since the base model had, arbitrarily, different orientation than that of Gata4).

### Selection of model developmental enhancers for deep dissection

As a starting point for our investigation, we used elements that were identified as cell-type specific in a recent screen of accessible chromatin regions in proximity of developmental loci using single-cell resolved MPRAs (scQer)^40^. As measured in a multi-cellular model of early development (mouse embryoid bodies), these included 10 developmental regions with activity restricted in the parietal endoderm lineage (n=7), in pluripotent states (n=2), together with one bi-functional element active in both lineages. The genomic coordinates and sequences of these ‘full length’ elements are listed in **Table S1**.

The initial screen considered larger regions spanning about 1 kb, in line with the typically large size of regulatory DNA tested in classic developmental reporter assays REF. To restrict the size of the sequence space to explore and facilitate downstream dissection, we sought to assess whether the regulatory activity of smaller subsets of the full 1 kb sized regions would suffice for regulatory activity. As such, we synthesized 270 bp sub-tiles with a stride of 5 bp. Of note, we extended the region of the Sox2 control region beyond the ‘full length’ element tested in the original single-cell reporter screen to include regions covered in a previous high-content dissection of the locus REF. All in all, this library consisted in n=1377 270 bp tiles, which were profiled in a single orientation (same as the original construct) in PYS-2 and mESC lines (**Fig. S2**). Data for the 270 bp tiles revealed a strong cell-type specific activity towards endoderm cells for 5 of the CREs.

The maximum-activity tiles for these 270 bp tiles traces were (one per full length CRE) selected for a comprehensive dissection of the sequence-to-function relationship (saturation mutagenesis, compaction, derivatization, and evolution^53^. To work with 300 bp tiles, 15 bp symmetric extensions were applied on both sides. In two out of five cases, there was a substantial drop in activity as a result of the symmetric 15 bp extension from the maximum activity 270 bp sub-tile (*Bend5*:chr4_8201 and *Sparc*:chr11_7211, see **Fig. S4**). With regards to this sub-optimal choice, libraries for the other class of sequence variants were designed before having generated the second round of sub-tiling (going all the way to 300 bp). Subsequent saturation mutagenesis indeed suggests repressing features near the 3’ segments of these tiles near minP (**Fig. S9**). The selected tiles together with the sequences of all synthesized CREs can be found in **Table S1**.

### MPRA libraries design

Metadata, details, CRE sequences and synthesized sequences (Twist Biosciences) are all listed in **Table S1**.

#### Multi-size sub-tiling

For each of the full CRE genomic intervals, a custom script was used to generate a bed file of all tiles included within the original interval of desired set of length (40, 70, 120, 170, 220, 300 bp). The same orientation as the full CRE was noted. From the coordinates/strand of the intervals, we used getfasta from Bedtools (version 2.29.2 with option –s to preserve the original orientation of the full CRE tile) to obtain the sequences from the mouse genome (mm10, coordinates now converted to mm39 in metadata tab in **Table S1**). For synthesis, constant handles for PCR (ctagtccaagcaagg– … –gtgcagcgcgatgta), specific to primers oJBL684 and oJBL685 (see below) were added, together with ‘dial out’ handles on the outside for extracting specific sizes from the synthesized pool separately (distinct for different sizes, see **Table S1**). To maximize usefulness of different synthesized oligonucleotides, some tiles of short sizes were combined on the same oligo (each with their own set of dial out primers).

#### Saturation mutagenesis

From the starting ‘maximum activity’ 300 bp tiles as noted above, a custom script was used to generate all point mutations (no deletion) at all positions within our model 300 bp element. Constant handles for cloning in the reporter backbone were added on both sides: (ctagtccaagcaagg– … –gtgcagcgcgatgta). ‘Dial out’ handles were finally added on the outside (one different per CRE) to enable separate cloning if needed.

#### Enhancer compaction

For each 300 bp ‘maximum activity’ mouse CRE tested, we employed *in silico* compaction to identify which single base pair deletion was predicted to enhance accessibility. Using a deep learning model (ChromBPNet^19^ trained on pseudobulked parietal endoderm accessibility from scATAC-seq in mouse embryoid bodies^40^) from our companion paper^53^, we predicted the effects of all possible deletions on accessibility (see below) and selected the most optimal mutation to increase accessibility. This iterative process was repeated until there were no more nucleotides to delete. All designs were generated through a custom Python script. Following this, we synthesized the compactified sequences from Twist Biosciences down to 40 bp. Dial out primer sequences used for amplification were oJBL864+oJBL881. Given the limitation in size recovery in the cloning process, we were only able to recover CRE sizes from 47-300 bp.

#### Enhancer derivatization – general notes

To dissect the sequence features governing CRE activity, we designed a combinatorial library of derivative CRE sequences based on our five 300 bp ‘maximum activity’ CRE tiles. For each CRE tile, we first identified endoderm-specific TFBS motifs (*Gata4/6, Sox17, Foxa2, and Klf4*, normalized affinity > 0.1). Overlapping TFBS intervals were merged, and the sequence was partitioned into a non-overlapping set of TFBS regions and intervening gap regions. All following classes of derivatized sequences below were generated through a custom Python script using these indivisible sets of TFBS as ‘anchors’ for sequence transformations.

#### Enhancer derivatization – Selection of background sequences

Background sequences were selected using the *prep nonpeaks* function in ChromBPNet^19^, applied to pseudobulk parietal endoderm scATAC-seq peaks. n=200 background sequences representing GC-matched, inaccessible genomic regions in mouse parietal endoderm, were selected to serve as negative controls. Each selected genomic background sequence was then dinucleotide shuffled (once per sequence), to generate another 200 sequences to augment our negative control set. These n=400 ‘background DNA’ sequences served as starting points for reconstitution and random deposition derivatization operations.

#### Enhancer derivatization – Reconstitution

We performed TFBS reconstitution by implanting the TFBS subsequences from each CRE into diverse background sequences (see above). The TFBS anchors were written over each background sequence at the exact start and end positions they occupy in the native CRE, preserving inter-motif spacing while replacing all non-TFBS anchor sequences with unrelated background. These reconstitutions were performed on both the original 200 backgrounds and the dinucleotide-shuffled version of each background (400 reconstituted sequences per CRE). To further assess how much TFBS-flanking sequence context contributes to activity, we repeated the reconstitution procedure using TFBS definitions trimmed by 1 bp and 2 bp on each flank (the “minus-1” and “minus-2” flank variants), generating an additional set of reconstituted sequences with progressively less native flanking context around each motif.

#### Enhancer derivatization – Random deposition

In addition to reconstitution, we performed random deposition of TFBS motifs into background sequences. For each CRE, the same set of TFBS anchors was placed into each background sequence at randomly selected, non-overlapping positions rather than at their native coordinates. Candidate insertion sites were drawn uniformly from positions 1 through (L – k), where L is the CRE length and k is the motif length; a position was accepted only if it did not overlap any previously placed motif. This was repeated with two fixed random seeds per background across the full background pool (both original and dinucleotide-shuffled variants) effectively generating more instances of reconstitutions (but with non endogenous TFBS positions), and additionally with 100 random seeds (100 different depositions) per CRE for a panel of six backgrounds.

#### Enhancer derivatization – Synthetic thripsis

Performing a synthetic analog of chromothripsis, for each CRE, N random cut sites were selected from positions within the gap (non-TFBS) regions of the sequence, partitioning it into N + 1 segments. These segments were then randomly shuffled, concatenating them in a new order (within the fixed orientation). Cut sites were constrained to fall outside merged TFBS anchor intervals, so individual motif sequences remained intact. This procedure was performed at three fragmentation levels (N = 5, 10, and 20 cuts), each with 200 independent random seeds per CRE, yielding a total of 600 thripsis variants per CRE tile. For displaying the thriptic fragments, there is some ambiguity regarding the generated fragments vs. the actual sequence content (e.g., if a fragment is short and extends the actual thriptic fragment to be longer than its origin would suggest). To avoid these ambiguities in our representation (**Fig. 5i, S25**), we bioinformatically reconstructed the thriptic fragments iteratively. Briefly, comparing the original sequence with the thripsized version, we searched for the longest common substring (function LCSn from package PTXQC). We then mapped and stored the respective positions within the two starting sequences. These positions were then masked, and the process repeated until the full CRE had been covered.

### Cloning barcoded reporters for 5’ MPRA

All libraries were cloned following a previously described protocol^40^, with slight modifications in some instances described below (e.g., libraries with variable size elements such as the sub-tiling). Generally, we performed three rounds of PCR using Kapa Robust (standard conditions) separated by dilutions to: (1) extract relevant subsequences from the synthesized oligonucleotide pool as necessary using library-specific ‘dial out’ primers (see **Table S1**) typically 10-12 cycles, (2) add homology arms to the MPRA backbone and minP (compaction: oJBL681+oJBL682; sub-tile round 2 in the reverse orientation: oJBL889+oJBL890, all other libraries: oJBL684+oJBL685) typically 5-7 cycles, (3) add the MPRA barcode (compaction: oJBL681+oJBL056, sub-tile round 2 in reverse orientation: oJBL890+oJBL056, all other libraries: oJBL684+oJBL056), typically 5-7 cycles. All steps of PCR were tracked by qPCR by spiking in SYBr green and stopped before the inflection point to avoid over-amplification. Between steps until the final amplifications, products were cleaned up with Ampure beads. Amplicon libraries after PCR3 were size selected on agarose on polyacrilamide gel depending on the final size. The resulting product was introduced in the 5’ MPRA backbone (p001-PB-MPRA digested with SbfI + AgeI and size selected on agarose) via Gibson assembly (nominally: 4 μL reaction at 0.02 pmole of backbone, 0.04 pmole barcoded library, NEBuilder HiFi). Following purification (Zymo Clean and Concentrator, 3:1 binding buffer) and elution in 6 μL of water, 2 μL were mixed with 20 μL of electrocompetent cells (C3020, NEB), electroporated (gene pulser, Bio-Rad), and outgrown in 500 μL of recovery medium for 1h before plating to assess library complexity and confirm good representation of BC per CRE. Transformants were then amplified overnight in LB with ampicillin before collection for plasmid purifications. Library integrity (proportion of plasmids with inserts) was confirmed by screening multiple colonies with colony PCR.

The following modifications to the above protocol can be noted: first, for the round 2 sub-tiling, libraries from each size were both amplified with primers to clone in the ‘normal orientation’ and primers that swap the identity of the homology from the backbone to ultimately generate reverse complement relative to minP (see **Fig. S4a**). Throughout, the different sizes were kept separate for PCR and only pooled equimolarly before the Gibson assembly to attempt to minimize size biases. Notably, the sub-tile round 1 (270 bp tiles) were also re-cloned in parallel and added to the final pool, such that the total number of sizes in the library (**Fig. S3**) was 7 (40, 70, 120, 170, 220, 270, and 300 bp). Second, in the case of the 20 control promoters (IGVF set: Engreitz.housekeeping.promoters.controls^114^), the library was bottlenecked to a lower complexity (63.1k BCs) given the small number of represented elements. Third, other control libraries (negative: pJBL002 minP only, positive: pJBL003 EEF1A1p including first intron) with 5’ BC MPRA architecture were cloned as follows: for pJBL002 the barcoded insert was a PCR product including minP and a 5’ BC created with oJBL047+oJBL056 using as template p001-PB-MPRA. For the pJBL003, the promoter insert was generated by PCR from plasmid Ef1_PURO_GFP (from S. Regalado) with primers oJBL049+oJBL050 (also adding the barcode). Their complexity was also bottlenecked explicitly to 9.0k for pJBL002 and 4.3k BCs to pJBL003.

### Barcode to CRE associations

We generated the dictionary connecting BC to their associated CRE for MPRA using short-read sequencing. In brief, the final purified 5’ MPRA plasmid libraries were taken as template for PCR appending Illumina adapters using primers flanking the CRE and BC (indexed-P5 primers oJBL057-oJBL059 binding upstream of the CRE and inside GFP oJBL060 appending P7). PCR were conducted by tracking using qPCR and stopping the reaction before the inflection point using standard conditions with Kapa Robust. Amplicons were purified with Ampure and paired-end sequenced on the NextSeq platform using the following primers (in all cases except compaction and sub-tile round 2): read1 CRE oJBL684, index1: BC oJBL061, index2: sample index oJBL064, and read2: CRE oJBL686. For sub-tile round 2, since primers oJBL684/686 bind the to the handle just flanking the variable and these handles are swapped for half of the library (reverse orientation), sequencing primers outside that region had to be used: read1: oJBL907, read2: oJBL908. Similarly, for the compaction libraries, the handles just outside of the CRE were different, requiring other primers to sequence the CREs, namely read1: oJBL681, read2: oJBL683.

Bioinformatically, the reads mapping to CREs were aligned to the expected set using Bowtie2^115^. Alignments combined to the BC using custom scripts, and number of reads summed to specific CRE-BC pairs. CRE-BC were thresholded on coverage on a library specific threshold (identified from the bimodality of coverage, where the low count pairs correspond to sequencing errors, PCR chimeras, or other molecular biology artifacts). Finally, total reads per BC were determined, and the fraction of reads attributable to a given BC from a CRE-BC pair was determined. BC with a fraction lower than 0.95 were not retained as they might be non uniquely paired.

The following libraries require special comments: For saturation mutagenesis, since mapping was for point variants instead of enhancer mapping, the identified mutations were also extracted from the alignment file, and compiled upon piling up the reads. There in addition to the total read cutoff and the requirement for unique pairing, a variant was retained only if it was supported by >90% of the reads associated with the BC (the reads covered the full CRE). For compaction CREs, sizes spanned a wide range of lengths, a single pooled alignment would allow a short compact CRE to map spuriously to an internal prefix of a longer compact CRE. To eliminate this ambiguity, we implemented a length-stratified alignment pipeline.

Briefly, paired end CRE sequences (159 nt on read 1 + 158 nt on read 2, giving enough overlap for a 300 bp CRE) were first assembled into a single contiguous read per fragment using PEAR^116^ (v0.9.11), then trimmed the constant regions outside the CRE (if it was shorter than read length) with Trim Galore (v0.6.x, https://github.com/FelixKrueger/TrimGalore), producing one PEAR-assembled CRE read per BC read pair. Next, for each CRE length L from 40 to 300 bp, we performed the following steps: (1) we extracted the subset of compact reference CREs with length equal to L using seqtk (https://github.com/lh3/seqtk) *subseq* on a list of entries identified with seqtk comp, and built a length-specific Bowtie2 index (bowtie2-build –-seed 42) containing only those references; (2) using the same seqtk *comp/subseq* procedure on the PEAR-assembled read FASTQ, we identified all reads with length equal to L and extracted both those reads and the matching mBC reads by shared read ID; and (3) we aligned the length-L CRE reads against the length-L Bowtie2 index using end-to-end alignment (bowtie2 –-end-to-end), which requires the read to match the reference over its full length and thus eliminates any partial-alignment artifacts that could otherwise associate short CREs with prefixes of longer ones. BC reads were then merged and tallied for each mapped CRE, and filtered for unique pairings as before.

### Cell culture and transfection of libraries

PYS-2 cells (ATCC, #CRL-2745) were grown in DMEM with sodium pyruvate (Thermo Fisher, cat. no. 11995065) supplemented with 10% FBS (Fisher Scientific, Cytiva HyClone fetal bovine serum, cat. no. SH3039603) and 1× penicillin/ streptomycin (Thermo Fisher, cat. no. 15140122). Cells were passaged every 48h. Cells were transfected using the standard ‘forward’ protocol (∼50-70% confluent cells in 10 cm plates per replicate amounting to about 5M cells, 10 µg of plasmid DNA per transfection). Medium was changed after 12h, the cells harvested at 48h post transfection (2× PBS wash, lifted with 0.05% trypsin) and fixed with 80% methanol before placing in –80C until RNA/DNA extraction. The transfection mixes for the different experiments were as follow: compaction (87% compaction library, 5% IGVF Engreitz housekeeping promoters, 5% full length CRE, 2.5% pJBL002 minP, 0.5% pJBL003 EEF1aP), derivatization (87% CRE derivative library, 5% IGVF Engreitz housekeeping promoters, 5% full length CRE, 2.5% pJBL002 minP, 0.5% pJBL003 EEF1aP), sub-tile round 2 & saturation mutagenesis (35% saturation mutagenesis library, 60% sub-titles round 2, 2% IGVF Engreitz housekeeping promoters, 2.5% pJBL002 minP, 0.5% pJBL003 EEF1aP), and sub-tile round 1 (2.5% pJBL002 minP, 2.5% pJBL003 EEF1aP, 30% sub-tile round 1, 5% full length CRE, 60% other libraries). We note that sub-tile round 2 and saturation mutagenesis were profiled at the same time (co-transfection), and that sub-tile round 1 was with other libraries not analyzed here. We caution against using even 2.5% of libraries to the EEF1A1 promoter given its strength as it can end up dominating the RNA barcode MPRA libraries.

mESC cells were cultured as previously described^40^. Briefly, a monoclonal male BL6 (male WD44, ES-C57BL/6 gift from C. Disteche/C. Ware,University of Washington) mES cell line stably expressing dCas9-BFP-KRAB was used (generated by S. Regalado and S. Domcke). Cells were grown on 0.2% gelatin-coated (Sigma, cat. no. G1890) plates and cultured in DMEM (Thermo Fisher, cat. no. 11965118) supplemented with 15% FBS (Biowest, Premium bovine serum, cat. no. S1620), 1× MEM nonessential amino acids (Thermo Fisher, cat. no. 11140050), 1 GlutaMAX (Thermo Fisher, cat. no. 35050061), 10^−5^ β-mercaptoethanol and 10^−4^ leukemia inhibitory factor (Sigma-Aldrich, ESGRO Recombinant Mouse LIF Protein ESG1107), with daily medium changes and passage every 48h. Transfection of libraries proceeded according to the ‘reverse’ method using Lipofectamine 2000 (Thermo Fisher, cat. no. 11668030). Briefly, cells washed with 1× PBS, lifted with trypsin 0.05% (Gibco), triturated with medium, and spun down at 300 g for 5 min. Following straining (40 µm), cells were counted and diluted to 0.5 M ml^−1^ with medium. Concurrently, the lipofectamine + opti-MEM and the opti-MEM + DNA solutions were separately prepared and mixed by pipetting. The 2.5 mL Lipofectamine + DNA + opti-MEM mix was then added to a gelatin-coated plate, 5M cells (10 mL) from the single-cell suspension was added to the plate, and gently mixed. For the sub-tile round 1 libraries, the 5 M cells per replicate were transfected with 20 µg of MPRA plasmid DNA (composition: 99.33% sub-tile round 1 library, 0.33% barcoded minP only negative control library, 0.33% barcoded EEF1A1 only positive control library). Medium was changed after 12h, the cells harvested at 48h post transfection (2× PBS wash, lifted with 0.05% trypsin) and fixed with 80% methanol before placing in –80C until RNA/DNA extraction.

### Bulk MPRA assay

Bulk MPRA was as follows: RNA and DNA were extracted from transfected cells with AllPrep kit (Qiagen). All PCRs listed below were with Kapa Robust (Roche) and standard conditions (95C denaturation, 60C annealing, 72C elongation, extension time 30 s). For the libraries of RNA barcodes: 5 μg of RNA was treated with Turbo DNase (stringent protocol), and directly transferred for a reverse transcription. 11 μL of the RNA was mixed with 2 μL of 1 μM primer oJBL789 (contains 10Ns serving as UMIs), denatured at 65C for 5 min and then placed on ice. 7 μL of master mix (4 μL 5x buffer, 1 μL 0.1 M DTT, 1 μL 10 mM dNTP, 1 μL SuperScript IV reverse transcriptase; Thermo Fisher) was then mixed in. The reaction was placed at 55C for 1h and then 80C for 10 min. A minus enzyme control is always added to confirm complete removal of DNA in the RNA prior to reverse transcription (should rise in the final PCR >10 cycles later than the sample with enzyme). PCR is then performed (25 μL reaction, using 10 μL of the RT mix directly) with primers oJBL790+nextera_i7 for 4 cycles with standard conditions. Following a 1.5x Ampure bead cleanup and elution in 12 μL 10 mM Tris 8, a final PCR tracked by qPCR was performed with oJBL076+oJBL077 (Illumina P5/P7 alone) and the libraries taken out before the inflection point (usually about 10 cycles). The final product is cleaned up with 1.5x Ampure beads and eluted in 10 μL for final quality control before sequencing. For the DNA libraries, 1-2 μg of DNA is amplified with oJBL789+oJBL790 for 4 cycles. Importantly, this low-cycle PCR’s purpose is to append a pseudo-UMI (contained in oJBL789) to collapse possible jackpotting from the second PCR. After 1.5x Ampure bead cleanup and elution in 12 μL, 4 μL from PCR1 are taken in 10 μL total reaction PCR2 with primers oJBL076+ nextera_i7, tracked by qPCR and stopped before the inflection point (typically 10 cycles for episomal MPRA).

The resulting libraries were sequenced on the NextSeq platform with the following architecture and primers: reads 1 MPRA BC for 15 cycles with oJBL072, index 1 sample index for 10 cycles with oJBL760 (Nextera idx1), index 2 MPRA BC reverse complement for 15 cycles with oJBL074, and read 2 UMI for 10 cycles with oJBL761.

### Quantification of regulatory activity from MPRA

Quantification of MPRA activity from RNA and DNA barcode sequencing was performed as before^40^. Raw fastq files were processed by trimming unnecessary cycles from the 3′ end as applicable for the specific sequencing run using seqtk. Forward and reverse BC reads were joined and error corrected with PEAR (options –v 15 –m 15 –n 15 –t 15). Using custom Python and R scripts, successfully assembled barcode reads were combined with UMI reads, BC–UMI pairs were counted, and the read and UMI counts per BC were determined. The read and UMI counts for the BC present in the reporter library (determined before the experiment when generating CRE-BC dictionaries) were collected for downstream analysis. The MPRA activity for a given CRE was computed by first determining the total UMI per sample mapping to the BC in our list was for both RNA and DNA libraries. Then, BC UMI counts were then normalized by the summed of UMIs in its respective sample type (DNA and RNA). The summed normalized RNA UMI count was then divided by the summed normalized DNA UMI count for all BC associated with the CRE. A 1% Winsorization was applied on normalized UMI values prior to summation to clip possible extreme BC outliers. Saturation mutagenesis data demanded a slightly different pipeline, as detailed below.

#### Saturation mutagenesis

Each variant in the library typically harboured multiple mutations (**Fig. S7c**), such that the standard approach described above is inapplicable. Similarly to previous strategies for this type of data^38^, to compute the variant effect of a given mutation, we first identified all barcodes associated with that mutation. Importantly, these barcodes could also be associated with other mutations, but had to at least be associated with the mutation of interest. Variant effect was then taken as the same RNA/DNA normalized and Winsorized summed UMI ratio as above. The underlying assumption for this approach is that other mutations present with the barcodes of a given mutation are averaged over and possibly only globally shift (down if there is prevalence of loss-of-function mutations) activity. This assumption breaks down in cases where the number of BC associated per variant is low, which typically was not the case for our current libraries with >100 BC per variant (except for deletions). To systematically assess the impact of this procedure, we isolated BC associated with a single mutation (no PCR-induced error in the cloning) in our dictionary, and BC associated with 2 or more mutations. We then computed the variant effects from the two barcode sets separately, and found overall remarkable agreement (**Fig. S8d**), except with a slight shift down for the multi-mutation variant effects, as expected. To calculate fold-change to WT in the most conservative way and to circumvent this systematic shift, we self-consistently took the activity value that led to the smallest number of significant non-zero changes.

#### Assessing pairwise TFBS epistatic effects

Our multi-hit mutagenesis library opened the possibility to measure epistatic interactions between mutations. However, the space of pairwise mutations even for a modest 300 bp element exceeds 400k. As such, our library complexity was insufficient to measure interactions at base-pair resolution. However, at the coarse-grained level of TFBS, where many mutations can similarly disrupt activity, we had sufficient representation. First, for each TFBS, we identified the top 10 activity-disrupting mutations within the top 15 affinity-disrupting variants for the associated TF (to have confidence of both functional impact and TF binding site disruption). Second, for TF pairs, the set of BC with at least one mutation from both constituents of the pair. These BCs were then used to estimate the activity of disrupting both TFs (collectively assessed by tallying RNA and DNA UMIs from BC associated with all variants with the TFs disrupted). The same procedure (selection of mutations) was applied for single TFBS and the functional effect of disrupting individual TFBS was calculated in an entirely analogous manner. Epistatic score was taken as the ratio of the observed pairwise disruption over the null multiplicative model expectation.

### ChromBPNet score (compaction)

To assess the functional relevance of each oligo sequence synthesized, we embedded each sequence at the center of 1,000 mouse genomic background sequences, which were deemed inaccessible for the specific cell type of interest (described in “*Enhancer derivatization – Selection of background sequences*”), using a custom Python script. We then averaged the predictions across these background sequences to compute the marginal footprint for each sequence. The final prediction scores were normalized by the null value, which represents the activity of the background sequence alone.

## SUPPLEMENTARY TABLES

**Table S1**: List of starting developmental CREs, final model 300 bp tiles for deep dissection, and synthesized sequences (sub-tiling version 1, sub-tiling version 2, saturation mutagenesis, compaction, derivatization), together with control sequences (minP, EEF1A1p, and housekeeping controls).

**Table S2**: List of used oligonucleotides, plasmids, and sequencing amplicons.

**Table S3**: List of mapped main parietal CRM TFBS in profiled CREs (threshold normalized affinity of 0.05).

**Table S4**: Selected TFBS anchors for CRE derivatization (**Fig. S17**).

## SUPPLEMENTARY FIGURES 1–25

**Supplementary Figure 1.**
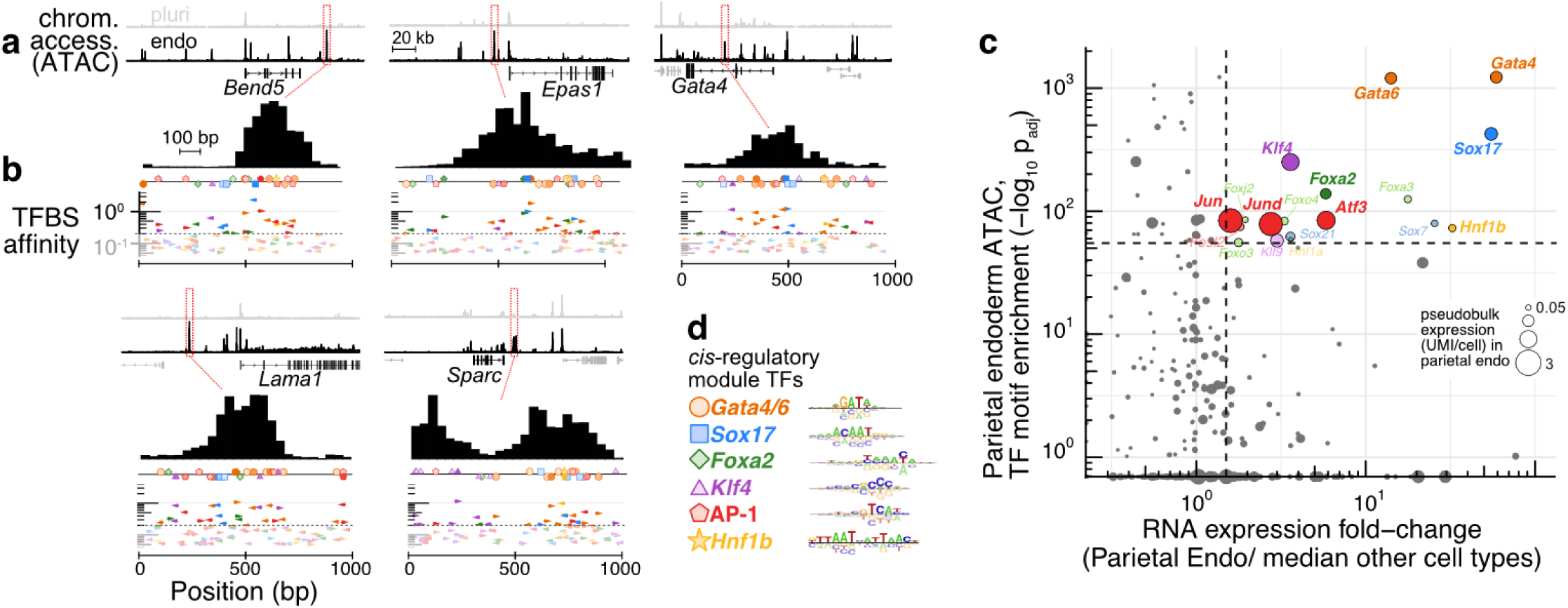
Profiled parietal endoderm CREs and *cis*-regulatory module TFs. **(a)** Upper: pseudobulk scATAC-seq profiles^40^ of pluripotent (grey) and parietal endoderm (black) cell types in mouse EBs in the vicinity of selected differentially expressed genes (200 kb windows). Profiled CREs are boxed in red. Lower: zoomed in view of the accessibility in parietal endoderm region over the full-length CRE included in the original screen^40^ (≈1 kb, variable from CRE to CRE as these were originally PCR-cloned). **(b)** Arrowheads: Putative TFBSs for *cis*-regulatory module TFs, affinity mapped using quantitative binding models^74^. The x-axis (position) is aligned to the x-axis of the lower part of panel **a**. Affinities are normalized to the 99.99% score among 1M random test k-mers to uniformize scores for binding sites of different lengths. A threshold of 0.2 (dashed line) was used to select binding sites shown at the top and also in **Fig. 1b**. **(c)** Identification of the parietal endoderm *cis*-regulatory module in mouse EBs. y-axis: TFBS enrichment in parietal endoderm (-log_10_ adjusted p-value from hypergeometric enrichment in marker peaks as computed using ArchR^109^ peakAnnoEnrichment function). x-axis: fold-change in pseudo-bulked mRNA expression levels of associated TF. scATAC and scRNA-seq data from mouse eEBs^40^. Highlighted TFs meet cutoffs of –log_10_p_adj_ >55 and expression fold-change >1.5. Top members of TF paralogous families with similar binding preferences are shown with the dark hue. Pseudobulk expression level in parietal endoderm maps to point size. Most TFs have no motif enrichment (-log_10_p_adj_ <0.6) and thus projected on or near the x-axis. **(d)** Shown are the model parameters (akin to a position weight matrix) from the MotifCentral database^74^ for the CRM TFs highlighted in panel **c** (MotifCentral IDs: FOXA2: 17084, GATA4: 16735, GATA6: 17095, HNF1B: 12948, KLF4: 17150, SOX17: 10629, JUN:ATF3 [AP-1]: 15946).

**Supplementary Figure. 2.**
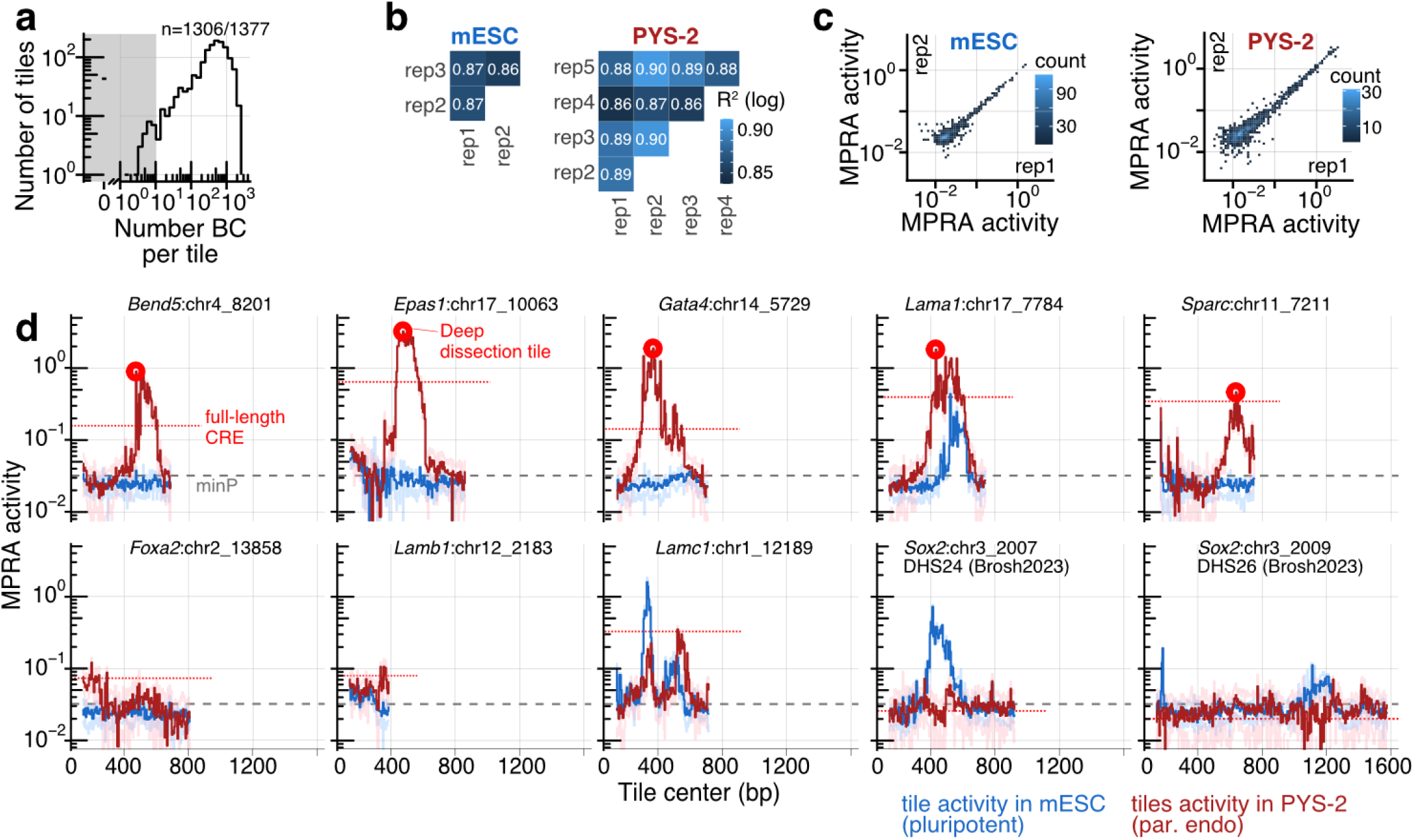
Single-size (270 bp), single-orientation sub-tiling of 10 developmental CREs in two cell lines. **(a)** Number of BCs per tile in the MPRA library dictionary of 270 bp tiles scanning across 10 developmental CREs identified by a scQer performed in mouse EBs^40^. Total library complexity: 658.6k BCs for 1377 tiles, with a median of 492 BCs/tile and n=1306/1377 with ≥10 associated BCs. **(b-c)** Reproducibility for episomal MPRA data in mESCs and PYS-2 cells across biological replicates, including Pearson’s R^2^ on log-transformed activities (panel **b**) and representative scatter plots (panel **c**). **(d)** MPRA data from the sub-tile scan (270 bp tiles) with a stride of 5 bp for the 10 developmental CREs, which are of different sizes. These include, from left-to-right then top-to-bottom, 7 specific to parietal endoderm, 1 bi-functional, and 2 specific to pluripotent cells, from the mouse EB scQer study^40^. Median activity across replicates per sub-tile position is shown (red: PYS-2, blue: mESC). Pale lines delineate standard deviation and min/max across replicates in PYS-2 and mESC respectively. The full-length versions of the 5 enhancers on the top row exhibited strong endoderm-specific activity in our scQer screen in mouse EBs^40^, while the full-length versions of the *Sox2*-control-region enhancers (*Sox2*:chr3_2007 and *Sox2*:chr_2009) were inactive in PYS-2 cells. Tiles used for fine dissection (saturation mutagenesis, compaction, derivatization) were taken as the 300 bp tiles centered on the maximum PYS-2 activity 270 bp tiles from each of the 5 enhancers on the top row (light red points). Notably, these all had activity >15-fold above the basal minP-only control in PYS-2, and were below the minP control in mESC cells.

**Supplementary Figure 3.**
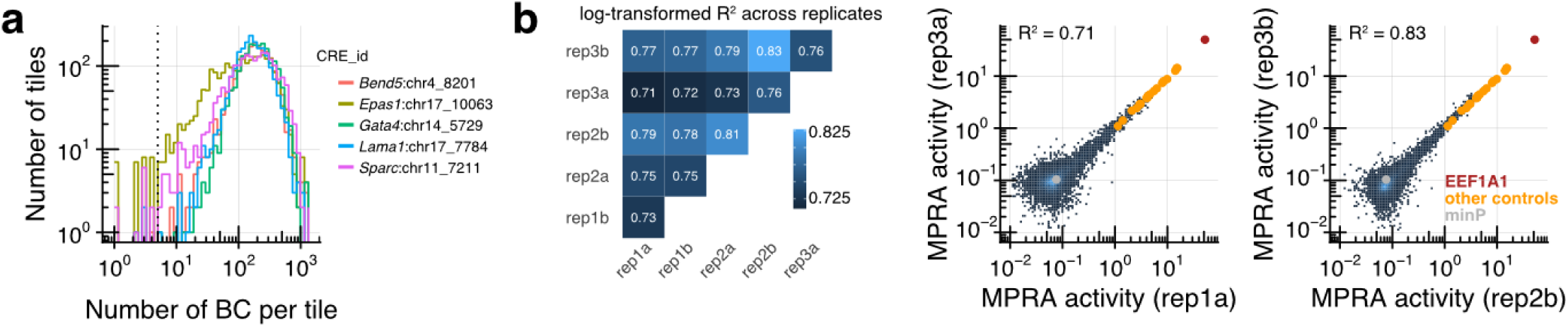
Quality metrics for the multi-sized sub-tiling MPRA library. **(a)** Distribution of BC counts per sub-tile (defined as a regulatory sub-fragment specified by tile size, genomic position, and orientation), stratified by CRE. The final reporter library recovered the vast majority of all possible sub-tiles (9,959 of 10,108; 98.4%) with ≥5 BCs per sub-tile, comprising 1.91 million total BCs. The interquartile range of BC counts per sub-tile was 94-253. **(b)** Reproducibility of MPRA activity per sub-tile across biological replicates. The heatmap reports pairwise R² values computed on log-transformed activity for all replicate pairs. The replicate pairs with the lowest and highest correlations are shown at right. Control promoters are highlighted for reference (minP-only in grey, the IGVF-20-promoter control set in orange, and the *EEF1A1* promoter in red).

**Supplementary Figure 4.**
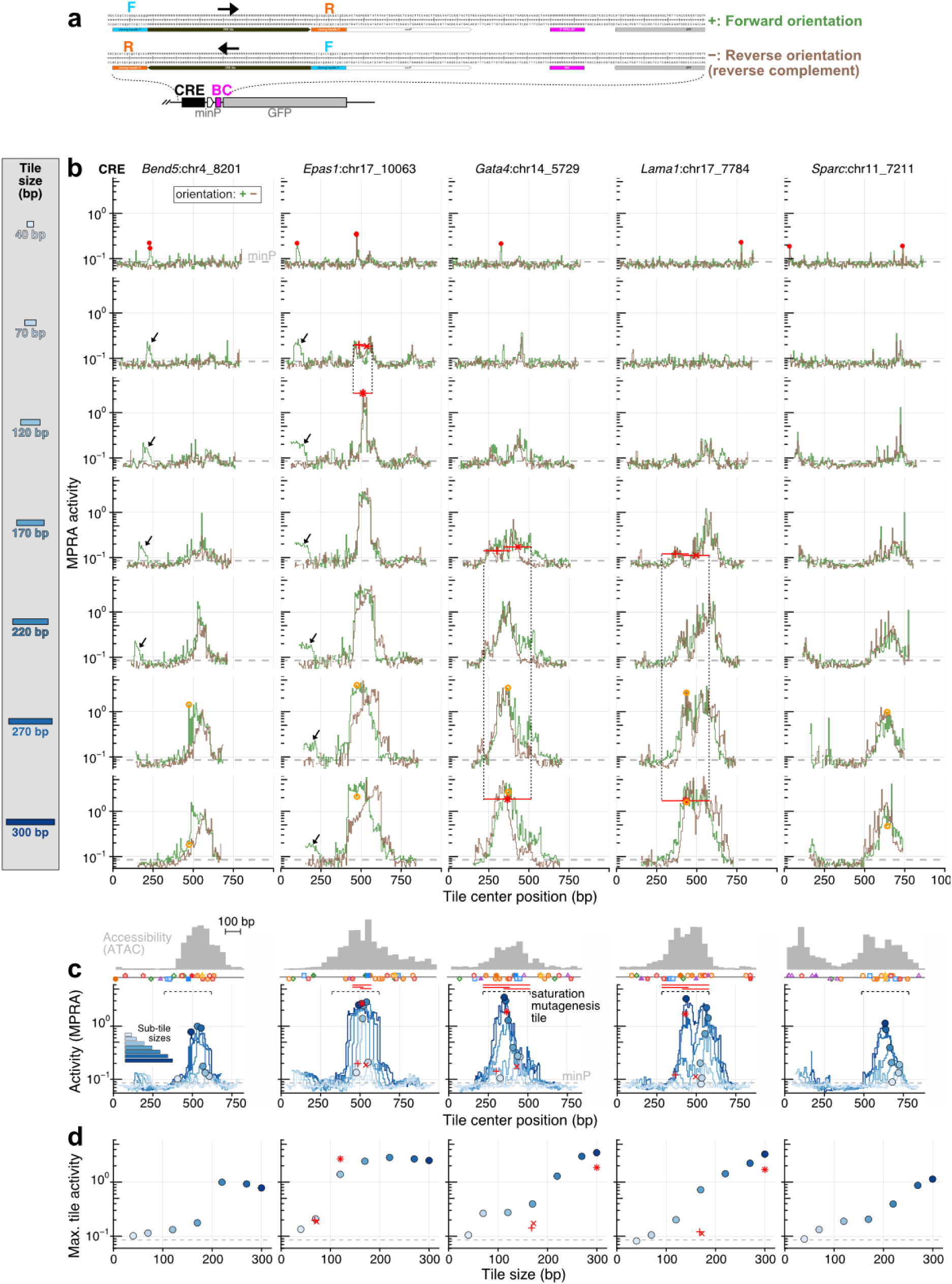
Dense, multi-sized, both-orientation sub-tiling MPRAs of high-activity developmental CREs. **(a)** Schematic of 5′ MPRA construct that illustrates how CRE sub-tiles were cloned in both forward and reverse orientations. Each CRE sub-tile contained distinct left and right 15-bp cloning handles (F and R) used for amplification from the oligo pools. Reverse-orientation constructs were generated using primers with swapped homology arms. Thus, tiles in each orientation are flanked by different handles relative to the minimal promoter (minP). Consequently, we cannot fully exclude the possibility that chimeric junctions between CRE sequence and cloning handles create additional TF binding sites that contribute to measured activity, although we consider this unlikely. **(b)** Unfiltered data from dense, multi-sized sub-tiling MPRAs of all five CREs, shown separately for forward (green) and reverse (brown) orientations. Each column corresponds to a CRE and each row to a tile size. The grey dashed line indicates the mean activity of the minP-only control. Red points in the 40-bp row denote the 9 tiles with activity >2-fold above minP background. Black arrows indicate extended regions of activity outside major chromatin accessibility peaks; of note, these were overwhelmingly orientation-specific. Red symbols (+, ×, ✴) mark the short and long synergizing tiles highlighted in **Fig. 2**. Orange points indicate the maximum-activity 270-bp tiles from our initial MPRA screen (**Fig. S2**) that were selected for saturation mutagenesis after extension by 15 nt on each side to generate a 300-bp tile; the activity of the resulting 300-bp tiles is shown in the bottom row. In one instance (*Bend5*:chr4_8201), the extended tile had drastically lower activity (see also **Fig. S9**). **(c-d)** Similar to **Fig. 2b-c**, but for all five CREs, and ordered horizontally in line with panel **b** above.

**Supplementary Figure 5.**
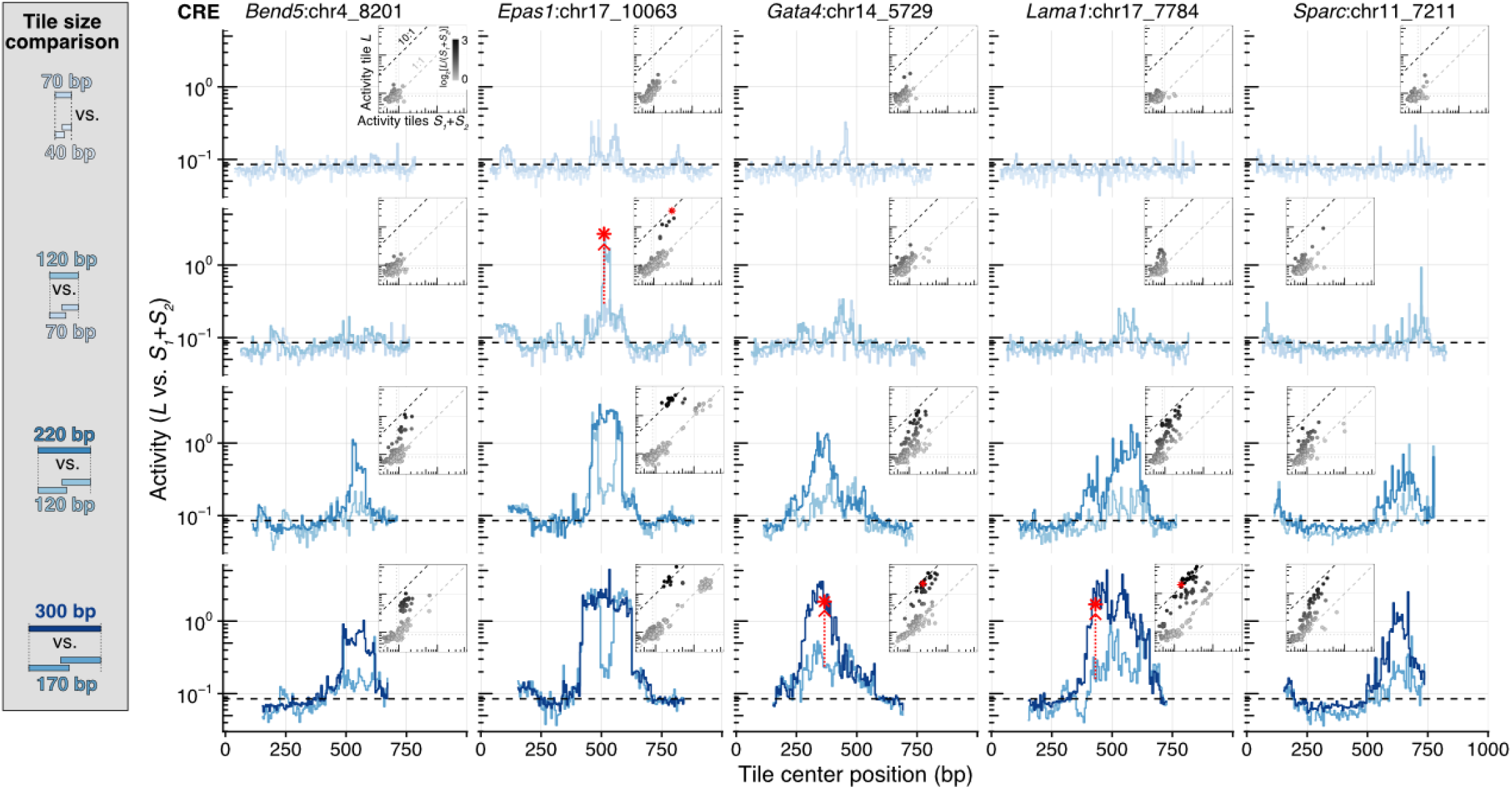
Sliding-window comparisons of matched short and long tiles reveal synergistic CRE sub-tiles. Similar to **Fig. 2d**, but shown for all CREs (columns) and for multiple short/long tile size pairings (rows: 40+40 vs. 70 bp, 70+70 vs. 120 bp, 120+120 vs. 220 bp, 170+170 vs. 300 bp), as illustrated to the left. For each comparison, two adjacent short tiles (*S_1_*and *S_2_*) are matched to the corresponding long tile (L) spanning the same genomic interval. Red points mark the tiles highlighted with red symbols in **Fig. 2b-c**, and red arrows below these points that indicate the fold synergy of the long tile (*L*) relative to the summed activity of the two short tiles (*S_1_*+*S_2_*). Each inset shows a direct comparison of *S_1_*+*S_2_* vs. *L* activity, where points deviating from the diagonal indicate synergistic sub-tiles. Color scale used for points in insets is shown only in the top left panel. Grey and black dashed lines in insets correspond to 1:1 (no synergy) and 10:1 (strong synergy).

**Supplementary Figure 6.**
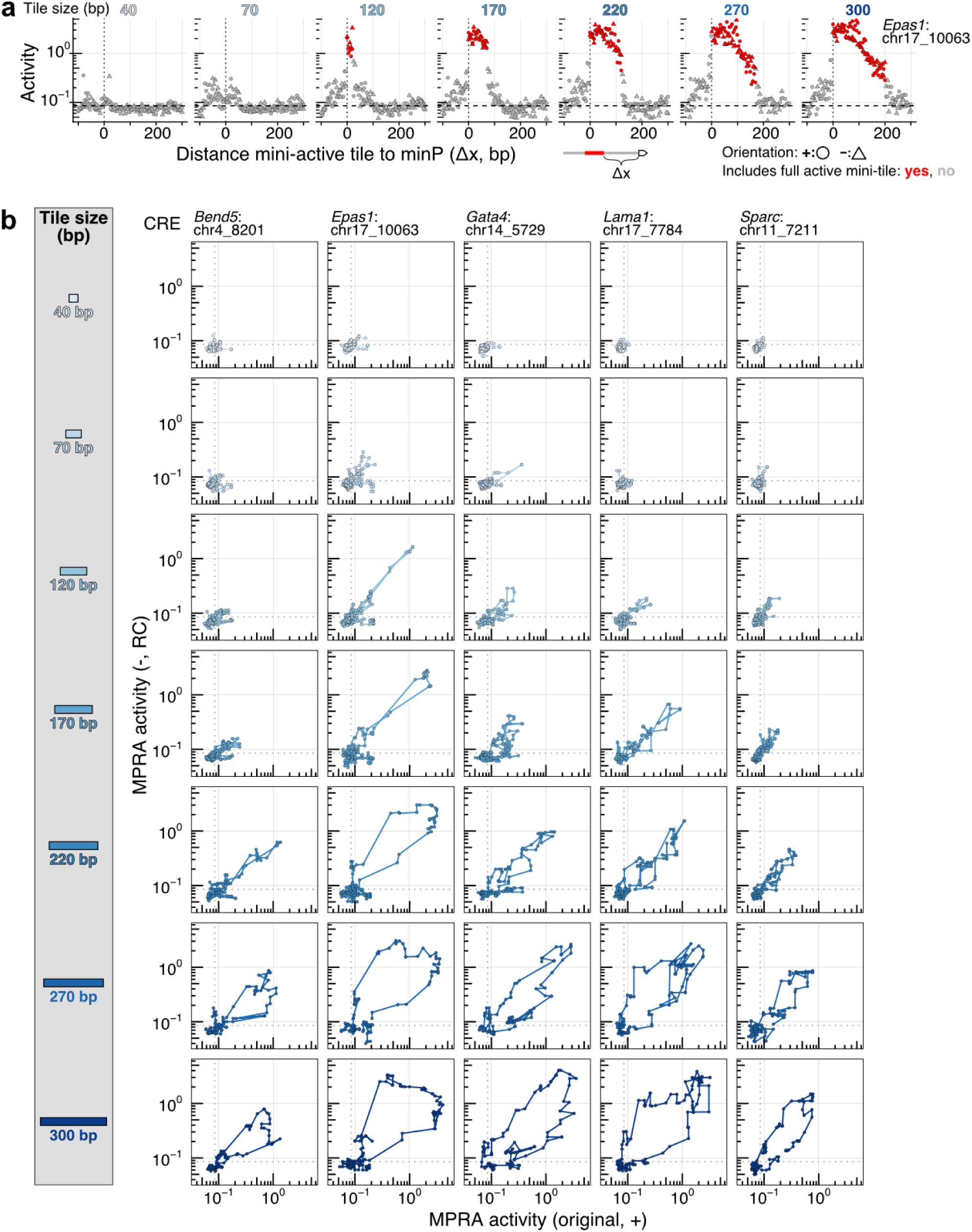
Multi-sized sub-tiling reveals orientation-dependent short-range positional effects in MPRAs. **(a)** Re-plot of the sub-tiling MPRA data for CRE *Epas1*:chr17_10063, with activity shown as a function of the distance between minP and the active mini-tile, rather than absolute tile position. Forward (circles) and reverse (triangles) orientation tiles collapse onto a common distance–activity curve, indicating that distance to the promoter explains a substantial fraction of the variability between orientations. Red points denote tiles that fully contain the active mini-tile, whereas grey points correspond to tiles that do not. **(b)** Direct comparison of MPRA activity for matched forward (+) and reverse (–) orientation tiles. Each panel corresponds to a different CRE (columns) and tile size (rows). The progressive opening of the forward/reverse trajectories at larger tile sizes indicates increasing orientation-dependent differences for longer fragments, consistent with a length-dependent positional effect on enhancer-minP activity in this reporter context.

**Supplementary Figure 7.**
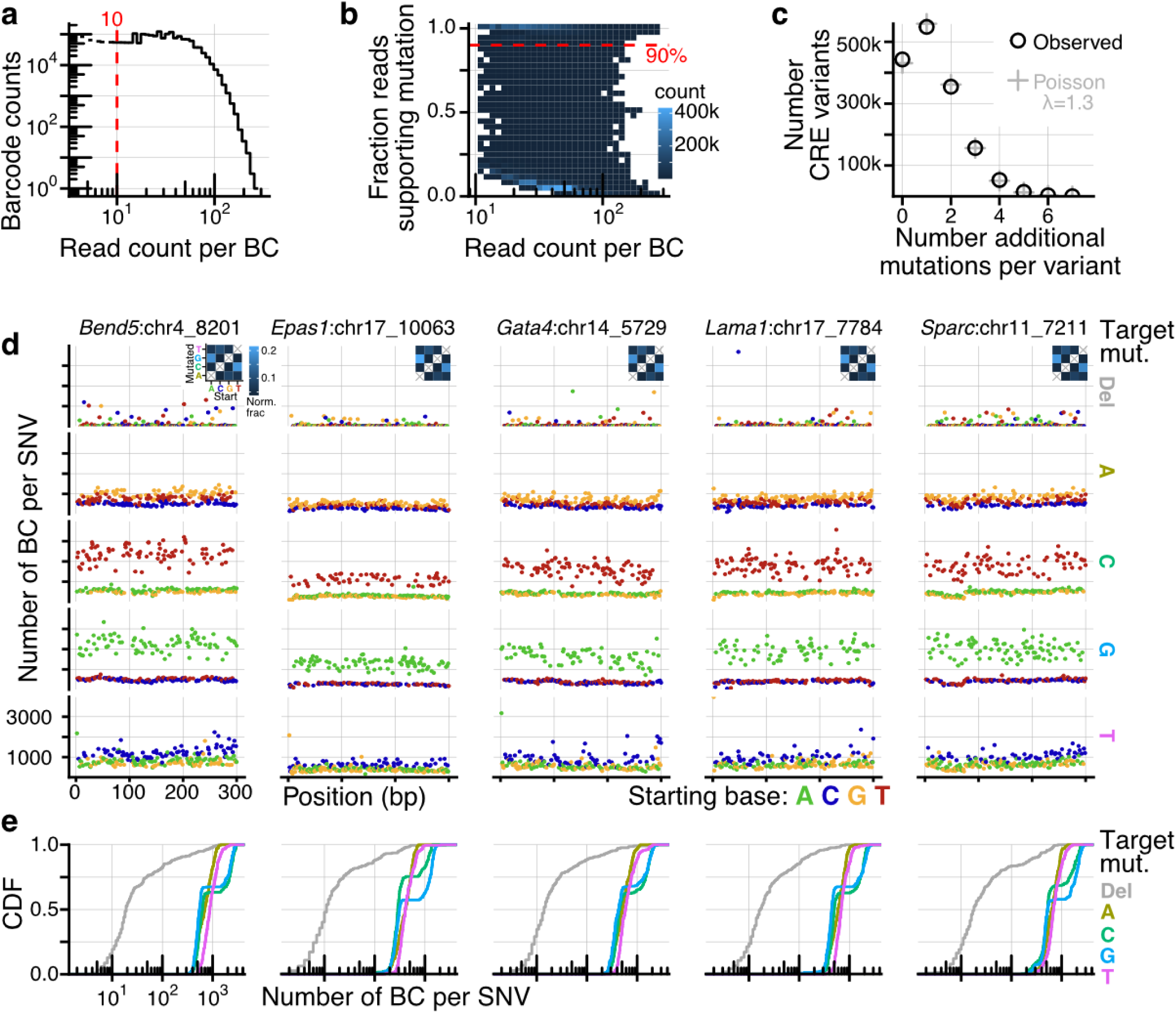
Quality control and characterization of multi-hit saturation mutagenesis library. **(a)** Distribution of read count per BC for the saturation mutagenesis MPRA library. The final library consisted of 1.6M barcodes (with >10 reads in the variant-to-BC association phase). **(b)** Two-dimensional histogram (color indicating density) showing read count associated with a given BC vs. fraction of reads supporting a specific mutation. Mutation-to-BC associations were called if >90% of reads supported the mutation (red dashed line). **(c)** Distribution of additional mutations relative to what was programmed (*i.e.* # mutations minus 1) mapped in the variant-to-BC dictionary. The observed distribution (black circles ‘o’) closely matches the Poisson distribution (grey crosses ‘+’), as expected for a random fixed rate mutagenesis. Mean of distribution is λ=1.3, corresponding average additional mutation rate of 0.4% per base from the PCR amplification and cloning process. **(d)** Characterization of the mutational spectrum: number of mapped barcodes with a given mutation (rows: different target mutation, columns: different CREs, colors: starting base). Taq-based polymerase preferences are immediately manifest (A:T→G:C) and add mutations over an otherwise constant baseline as expected from commercially synthesized libraries with a single programmed point mutation per sequence. Inset shows quantification of the mutational spectrum using a normalized fraction accounting for base composition. **(e)** Cumulative distribution of barcode per target mutation ID. Cs and Gs BC counts are bimodal as a result of the putative PCR-derived mutations manifesting itself in only ⅓ possible contexts.

**Supplementary Figure 8.**
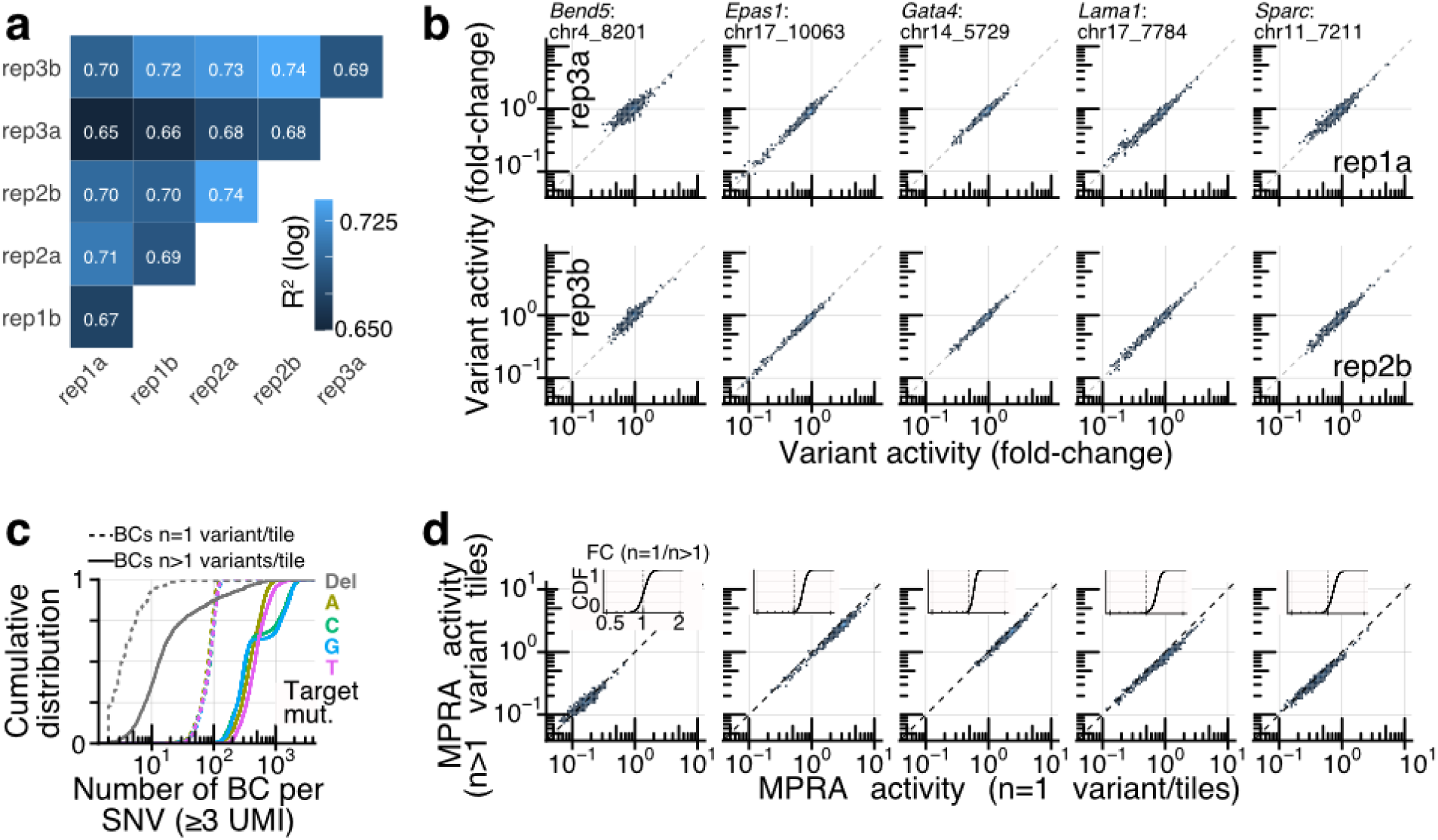
Saturation mutagenesis MPRA data quality metrics. **(a)** Reproducibility of MPRA activity for saturation mutagenesis across replicates. The heatmap reports pairwise R² values computed on log-transformed variant effects for all replicate pairs, *i.e.* 2 technical replicates for each of three biological replicates. **(b)** Variant effects comparisons for replicate pairs with the lowest and highest correlations. Each column corresponds to a different CRE and each row is a different comparison (top: rep1a vs. rep3a, bottom: rep2b vs. rep3b). **(c)** Distribution of number of BC per target mutation (with at least 3 UMIs mapped in the dictionary) in the library, stratified by whether the tile has a single mutation (‘n=1’ tiles) vs. multiple mutations (‘n>1’ tiles). As a result of the inferred PCR mutagenesis, most CRE variants bear multiple mutations. **(d)** Comparison of variant effect estimated from ‘n=1’ (x-axes) vs. ‘n>1’ (y-axes) tiles. Each column corresponds to a different CRE as in panel **b**. Inset shows cumulative distribution of fold-change between ‘n=1’ vs. ‘n>1’ tiles, showing a slight systematic shift to lower values in the ‘n>1’ tiles. In panels **b** and **d**, deletions are not included to avoid overt counting variability given their low representation in the library (see panel **c** and **Fig. S7e**).

**Supplementary Figure 9.**
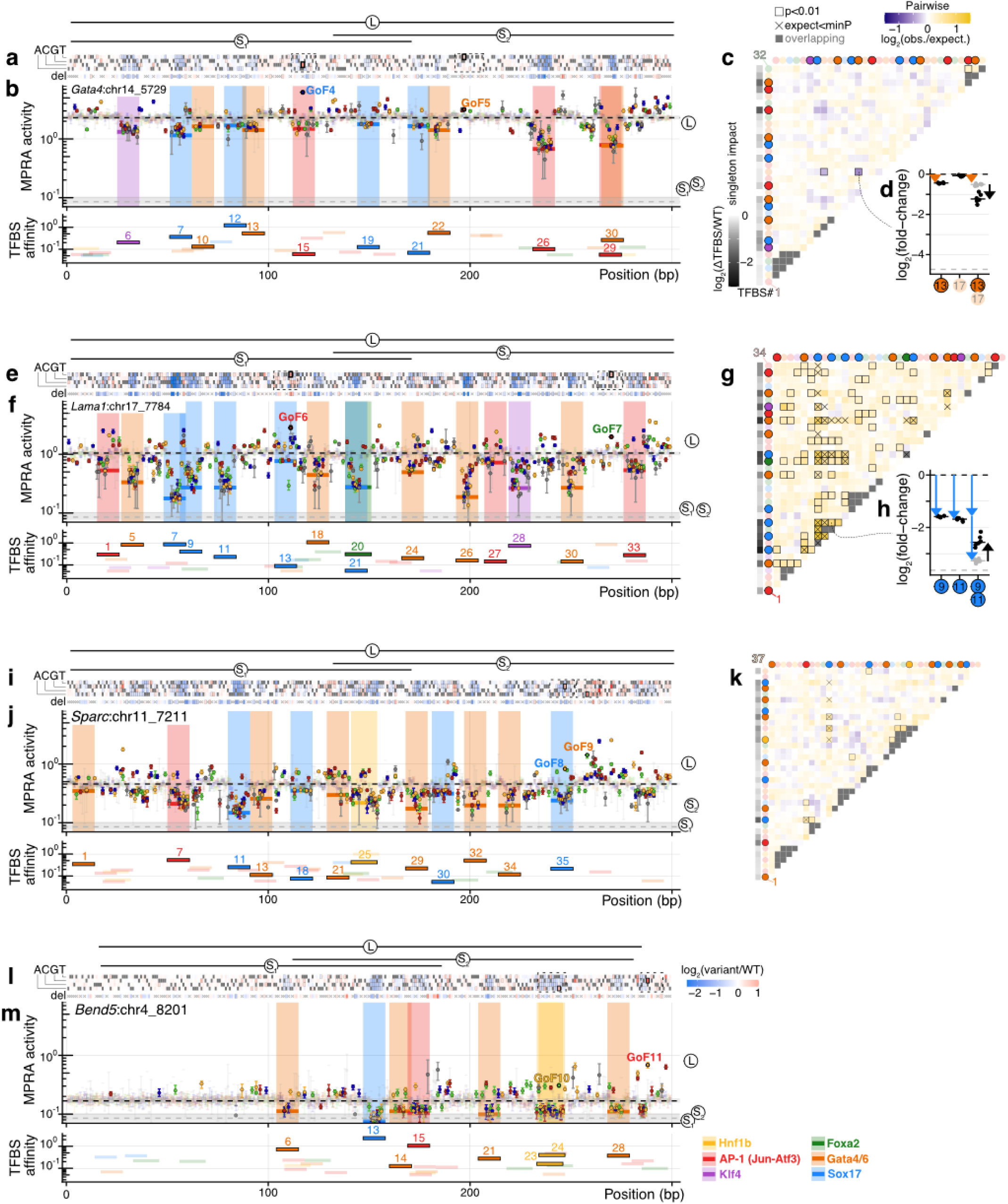
High-resolution saturation mutagenesis TFBS maps for four of the model CREs. Similar to **Fig. 3**, but here for the *Gata4*:chr14_5729 (panels **a-d**), *Lama1:*chr17_7784 (panels **e-h**), *Sparc*:chr11_7211 (panels **i-k**), and *Bend5*:chr4_8201 (panels **l, m**) CREs. The activity of the mutagenized tile for the *Bend5:chr4_8201* CRE was too close to the minP baseline to allow mapping of epistatic interactions.

**Supplementary Figure 10.**
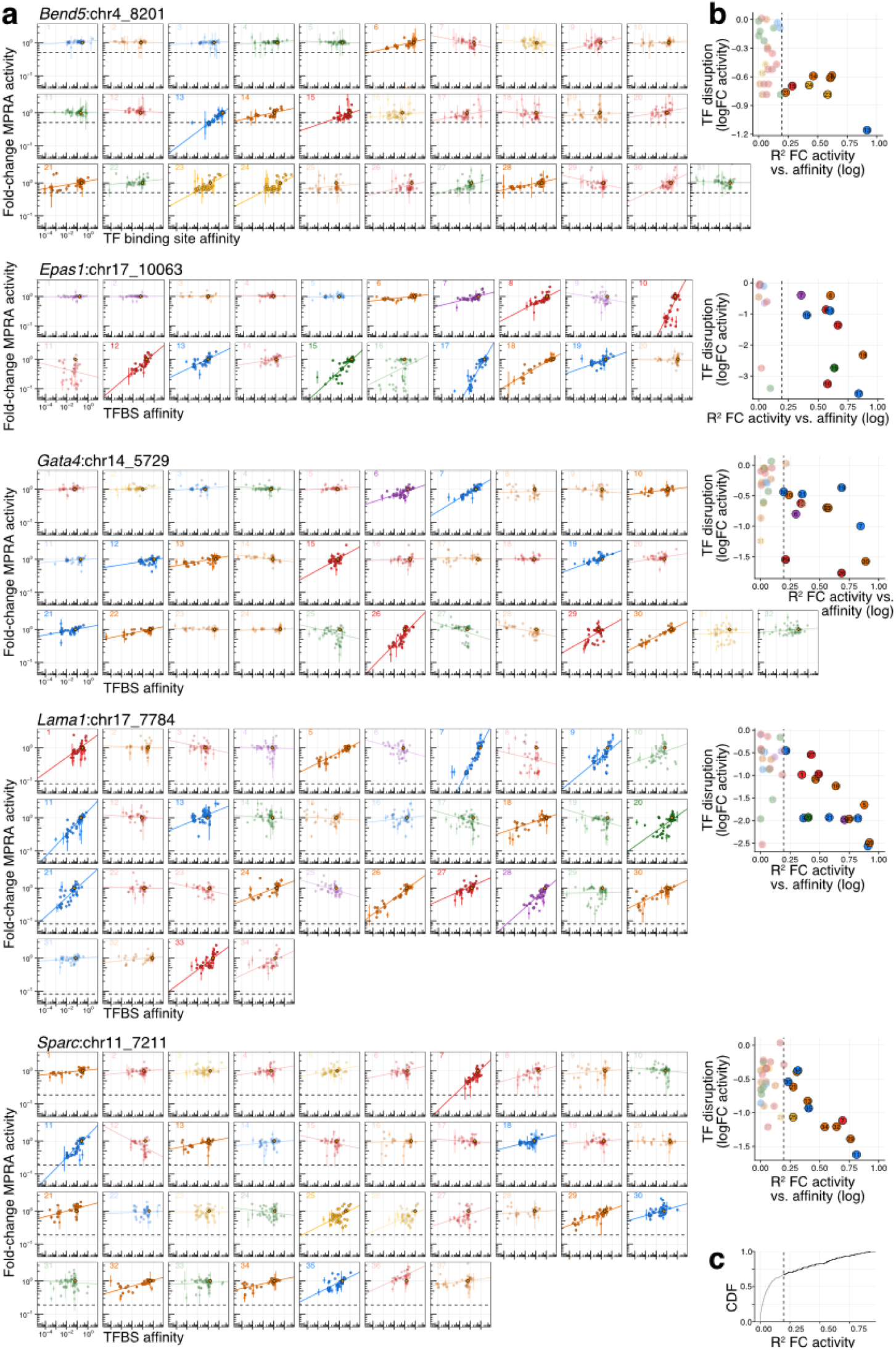
Affinity vs. activity correlation for putative binding sites of parietal endoderm TFs. **(a)** Multi-panel view of correlation between variant effect predicted affinity (x-axes) vs. measured activity (y-axes) for mutations within mapped TFBS, organized by CREs. Each plot corresponds to mapped TFBS (numbering follows positional order from left to right along the sequence). Each point in each panel corresponds to a mutation of the TFBS. WT sequences are marked with yellow diamonds. Errorbar spans the standard deviation across replicates. Line corresponds to linear regression on the log-transformed variables. Non-singleton-functional TFBS are shaded in a paler hue. Colors for different TFs match those of **Fig. 1b**. The expected fold-change to minP only is shown as a horizontal dashed line. **(b)** Plot per CRE showing the relationship between the fold-change in activity from disrupting the TFBS (y-axes) vs. the R^2^ correlation in the log-transformed plots shown in panel **a** (x-axes). Binding site color and numbering follow those of panel **a**. Singleton-functional TFBS are defined as those with R^2^>0.195 (dashed line). **(c)** Cumulative distribution of R^2^ across all TF binding sites illustrating the threshold used to designate singleton-functional TFBS.

**Supplementary Figure 11.**
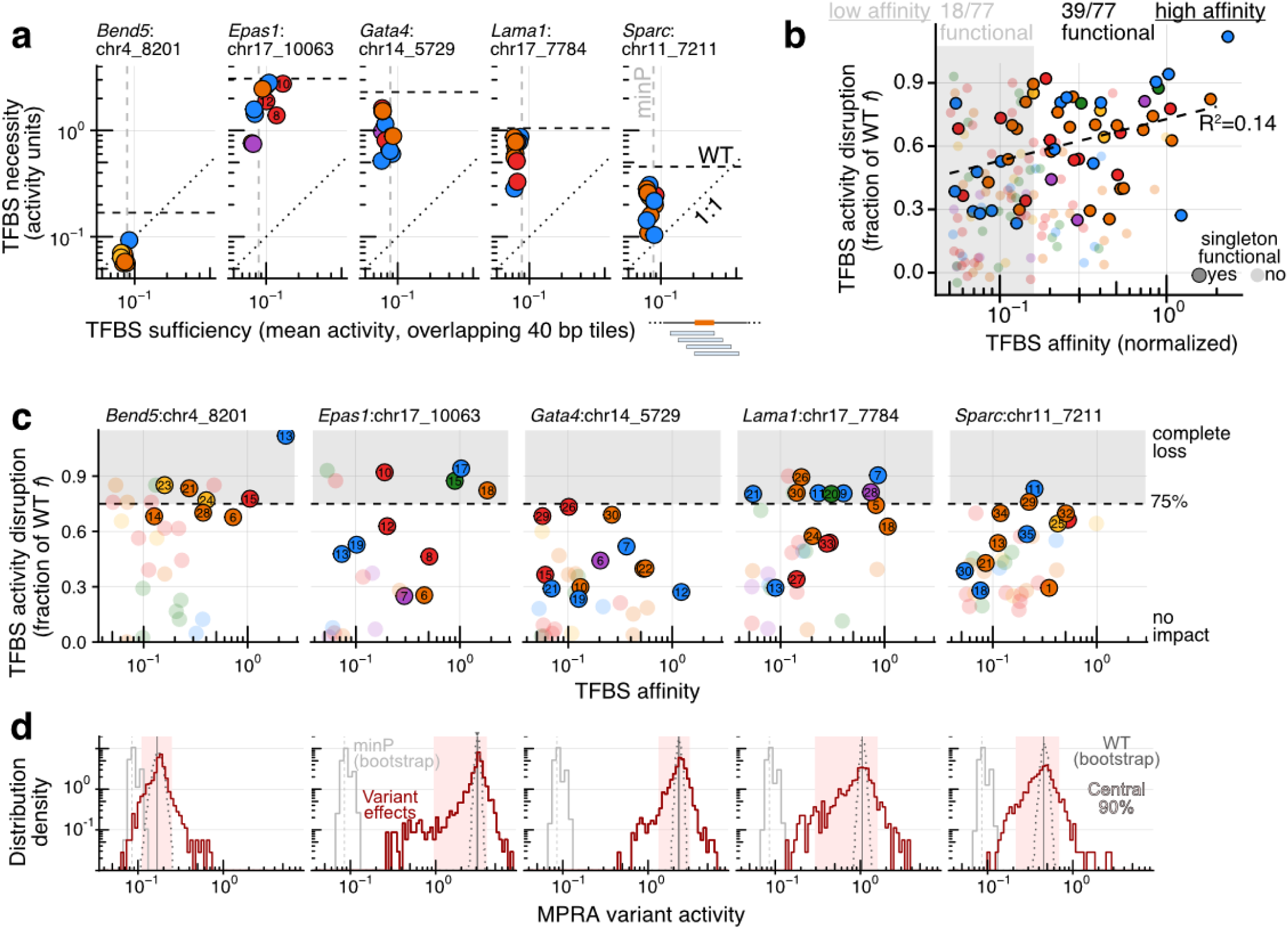
Sufficiency & necessity of TFBSs for activity and variant effect distribution. **(a)** Quantification of the sufficiency (x-axes: log-scaled average activity of all 40 bp tiles bearing the TFBS from sub-tiling experiment shown in **Fig. 2**) vs. necessity (y-axes: log-scaled activity disruption from panel **b**) of individual TFBS within each of the model CREs. **(b)** Necessity scores of TFBSs (*y-*axis) are only weakly predicted by affinity (*x-*axis). Necessity scores are quantified as in **Fig. 3h**, but plotted here for all TFBSs mapped to parietal endoderm CRM TFs. Singleton-functional sites are marked with a dark edge. Although affinity correlates only weakly with functional importance, higher-affinity sites are ∼2-fold enriched for singleton-functional TFBSs (Fisher’s exact test: p < 0.0007). **(c)** Quantification of necessity of individual TFBS (y-axes: log-scaled activity disruption: fraction of WT *f* lost upon disruption) as a function of affinity (x-axis). Grey zone delineates binding sites that abrogate >75% of WT CRE activity upon disruption. Binding site numbering and shading follows **Figs. 3b**, **S9b,f**,**j**,**m**. **(d)** Distribution of MPRA activity variant effects (dark red) for each of the model CREs (ordered as in panel **b**). WT activity is marked with a vertical black line. Bootstrap resampling distribution of the WT sequences matching the coverage of mutations within the respective CRE is shown in dotted grey line. Central 90% of variant effects is shown as a light pink shading. Bootstrap resampling for minP is shown as a solid gray line (same subsampling for all sub-panels).

**Supplementary Figure 12.**
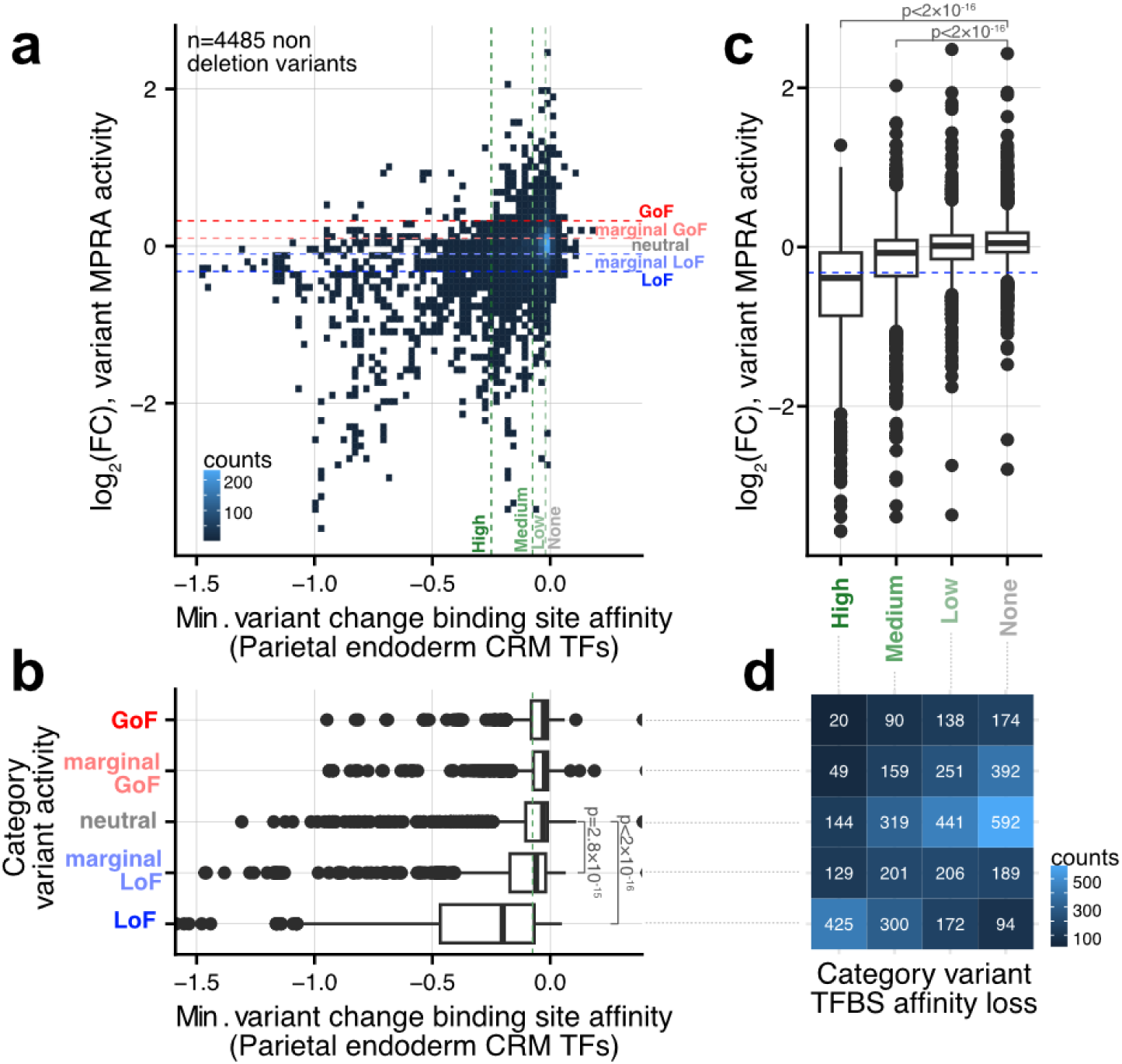
Change in TFBS affinity vs. change in TFBS activity for loss-of-function mutations. **(a)** Two-dimensional histogram of correlation between the maximal decrease in normalized affinity δ_min_ to parietal endoderm CRM TFs (x-axis; Gata4/6, Sox17, Klf4, Foxa2, AP-1, Hnf1b) vs. variant effect (y-axis; log_2_ fold-change activity) for every possible substitution in all five model CREs. Categorizations used in panels **b-d** are indicated by dashed lines. To calculate δ_min_, systematic mapping of TFBS is done and the CRM TF associated with the largest decrease in normalized affinity at that position is assigned. **(b)** Boxplot showing δ_min_ values stratified into categorized functional impact of the mutation (neutral: |log_2_FC|<0.1, marginal LoF: –0.32<log_2_FC<-0.1, LoF: log_2_FC < –0.32, marginal GoF: 0.1<log_2_FC<0.32, GoF: log_2_FC> 0.32). Marginal LoF and LoF categories are associated to significantly more TFBS disruptions as quantified by the maximal decrease in normalized affinity δ_min_ (one-sided Wilcoxon test p<10^-14^ compared to neutral mutations). Concretely, ∼75% of LoF variants (>1.25-fold decrease) are associated with δ_min_< –0.075 (vertical dashed line). **(c)** Converse plot of functional impact stratified by nature of TFBS disruption (none: δ_min_>-0.02, low: –0.075< δ_min_<-0.02, medium: –0.25< δ_min_<-0.075, high: δ_min_< –0.25). Medium/high disruptions to affinity lead, on average, to significantly lower activity than non-disruptive mutations (one-sided Wilcoxon test: p < 10^-15^). **(d)** Heatmap showing the number of mutations in the different bi-dimensional categorizations, *i.e.* the relationship between the extent of loss-of-affinity (columns; four categories aligned to panel **c** above) and the change-of-function (rows; four categories aligned to panel **b** to left).

**Supplementary Figure 13.**
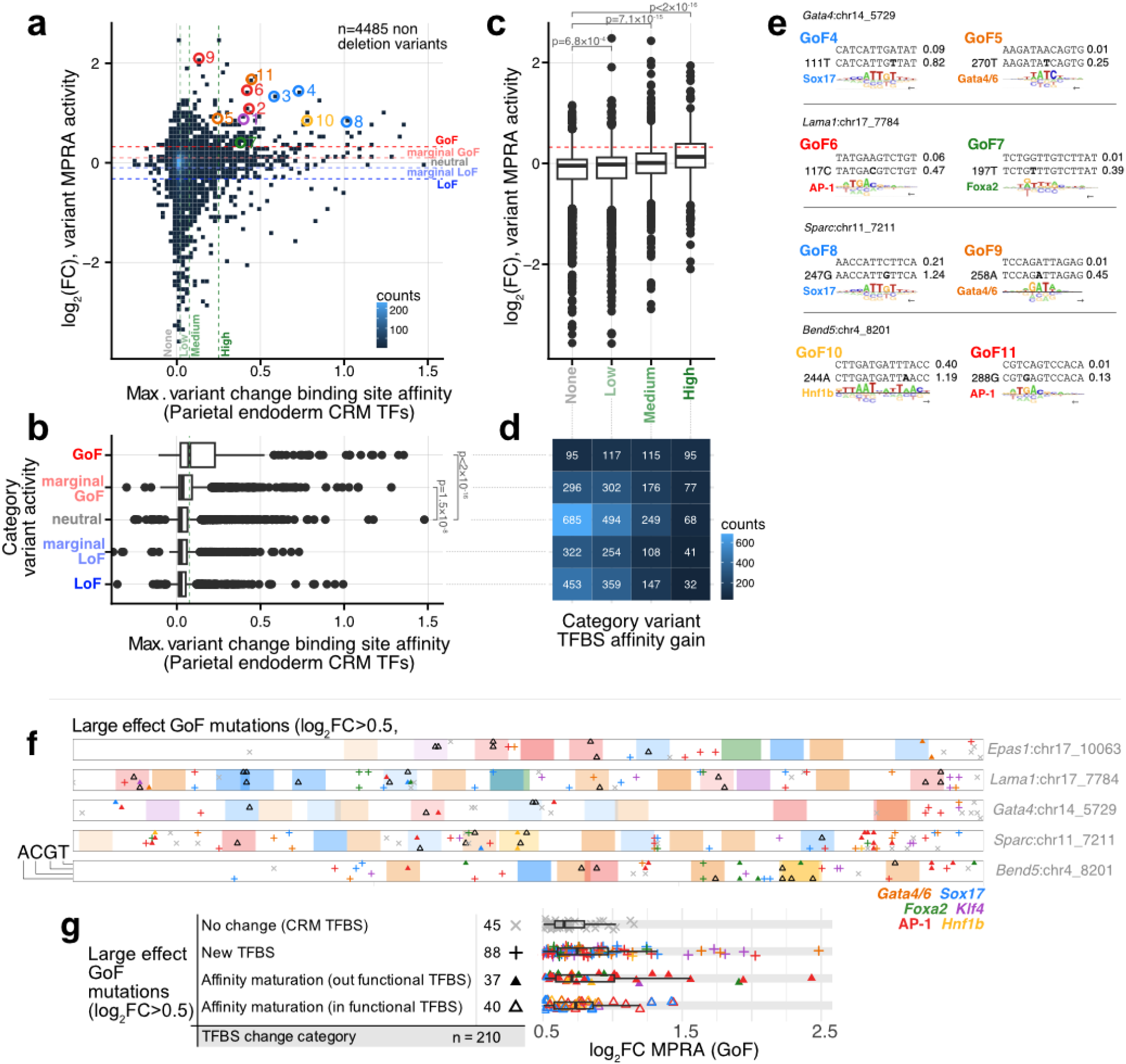
Change in TFBS affinity vs. change in TFBS activity for gain-of-function mutations. (**a-d**) Similar to **Fig. S12**, but here focused on the maximal increase in normalized affinity δ_max_ to parietal endoderm CRM TFs. In panel **a**, specific GoF mutations highlighted in **Fig. 3** and **S9** are marked. **(e)** Same as **Fig. 3d**, but here for the 8 GoF mutations highlighted in **Fig. S9**. One example per CRE was selected to be related to affinity-maturation (left column) and another to *de novo* TFBS creation (right column, corresponding to mutation in the core segment of the motif). **(f)** Heatmap summarizing the distribution of large GoF mutations (log_2_FC > 0.5; n = 210) for all five CREs (rows). The locations of singleton-functional TFBSs are delineated by shading, with hues map to TFBS identity (key lower right) and intensities to necessity scores. The locations of GoF mutations are marked by different symbols based on the nature of the TFBS changes: grey x: change not mappable to a CRM TFBS; open coloured triangles: affinity-maturation of a singleton-functional TFBS (class i in text); full coloured triangle: affinity-maturation of a non-singleton-functional TFBS (class ii in text); colored +: creation of new TFBS (class iii in text). Colors of symbols correspond to the associated TF, and their vertical position to the identity of the substitution. **(g)** Summary of the number of GoF mutations in each mechanistic category shown in panel **f**. Effect-size distributions for corresponding rows of the GoF categories shown at right. No pairwise comparison between categories was significant (two-sided Wilcoxon rank sum test with Bonferroni correction: p > 0.42).

**Supplementary Figure 14.**
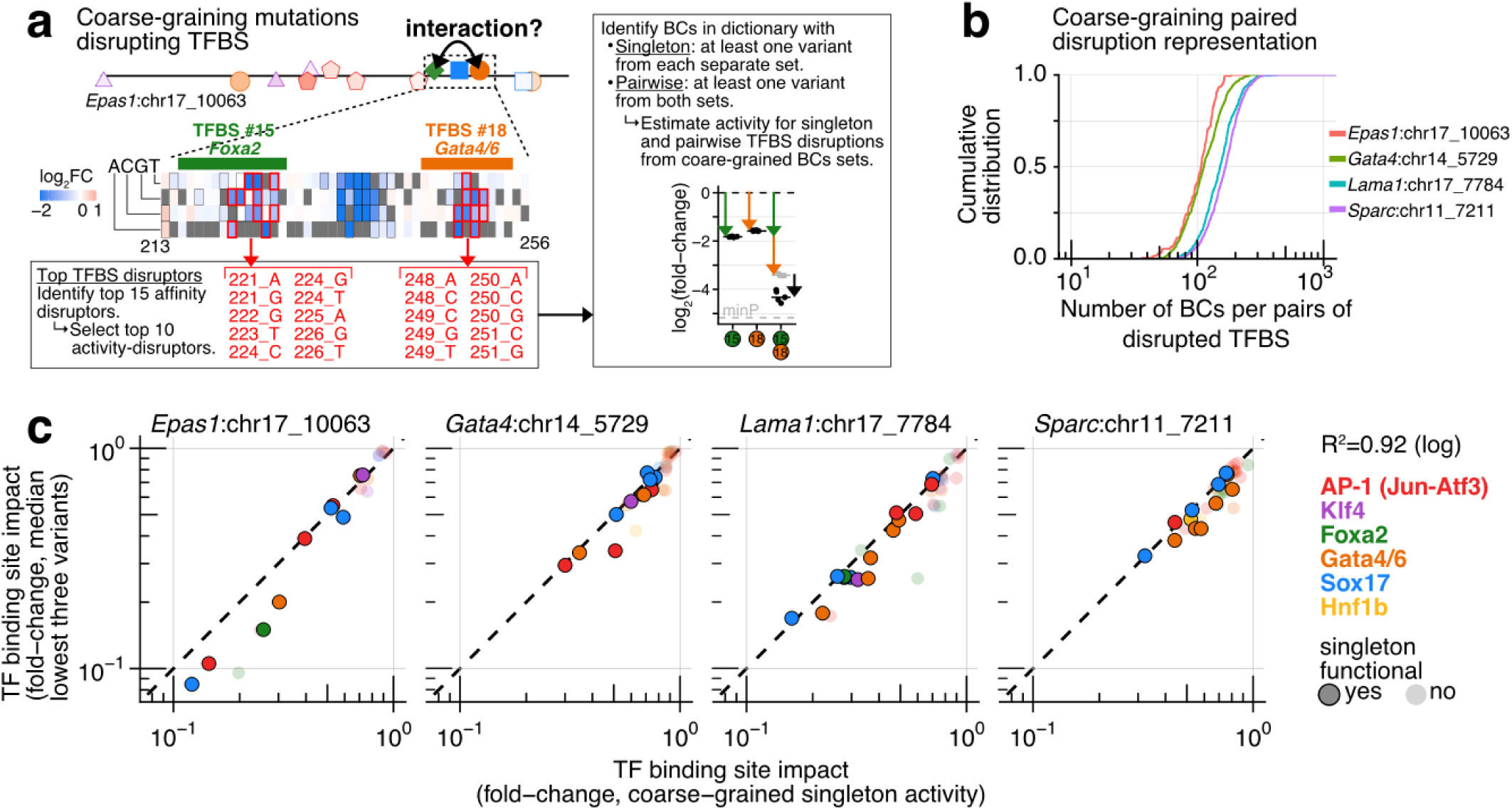
Details of coarse-graining procedure for pairwise TFBS epistatic analysis. **(a)** Illustration of the procedure used to coarse-grain mutations at the level of full TFBSs to gather enough statistical power for epistatic mapping. In brief, for each TFBS, the top fifteen affinity-disrupting mutations were identified from the biophysical binding model, and the top ten activity-disrupting mutations among those were identified. All barcodes harbouring any of these TFBS disrupting variants were retained for activity estimates. For coarse-grained epistatic mapping, barcodes associated with at least one TFBS-disrupting variant from both binding sites in a pair were used for activity estimates. **(b)** Cumulative distribution of the number of BCs in the variant library associated with all pairs of disrupted TFBSs. **(c)** As a control for the coarse-graining procedure, the functional importances of single TFBSs were computed using the set of BCs identified above. Comparison of these coarse-grained ‘singleton’ binding site disruption estimates (x-axis) to the binding site impact as estimated by the median of the top three disrupting variants per position (from the full saturation mutagenesis, not coarse-grained) showed overall excellent agreement, albeit with the expected slight underestimation from coarse-graining.

**Supplementary Figure 15.**
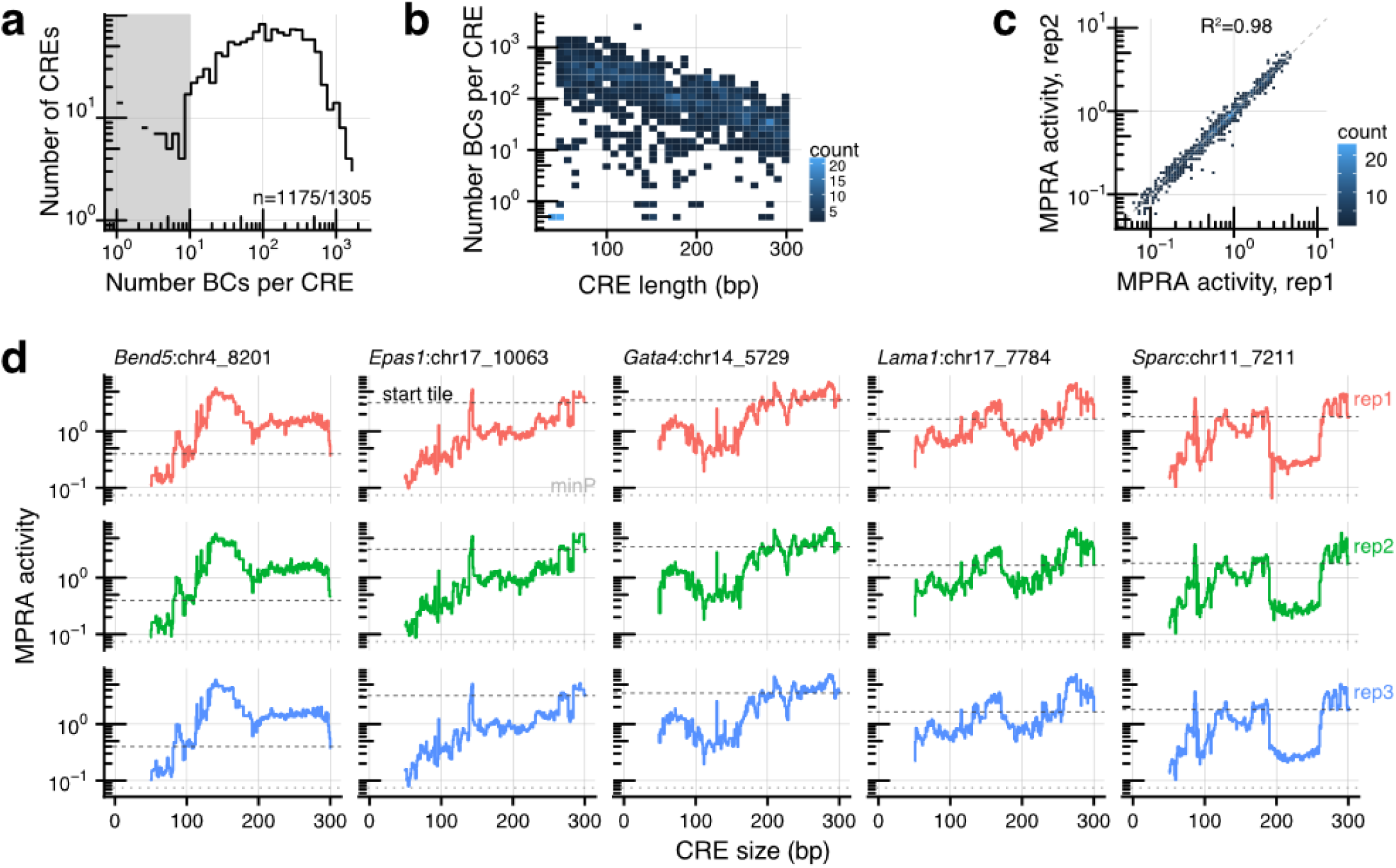
Dictionary quality metrics and reproducibility of CRE-compactification MPRA data. **(a)** Distribution of number of BCs per compacted CRE (median 94 BCs per CRE; total = ∼226,000; 1175 of 1305 CREs were represented by >10 BCs. **(b)** 2D heatmap highlighting a negative correlation between element size (CRE length in bp; x-axis) and representation in the library (# of BCs; y-axis), which led to a ∼10-fold difference in the number of BCs for elements below 100 bp vs. above 250 bp. However, although shorter elements were more highly represented, very short tiles were strongly depleted due to gel-based size selection during cloning (*e.g.* all 35 tiles of length 46 bp or shorter had 0 BCs). **(c)** Example correlation in MPRA activities across biological replicates. We note that correlation is higher for this experiment presumably because of >20-fold more sequencing coverage compared to other experiments. **(d)** Direct visualization of the different compaction trajectories across replicates to highlight the biological as opposed to technical origin of the ruggedness in the data (y-axis: MPRA activity, x-axis: CRE size; a different model CRE is represented in each column).

**Supplementary Figure 16.**
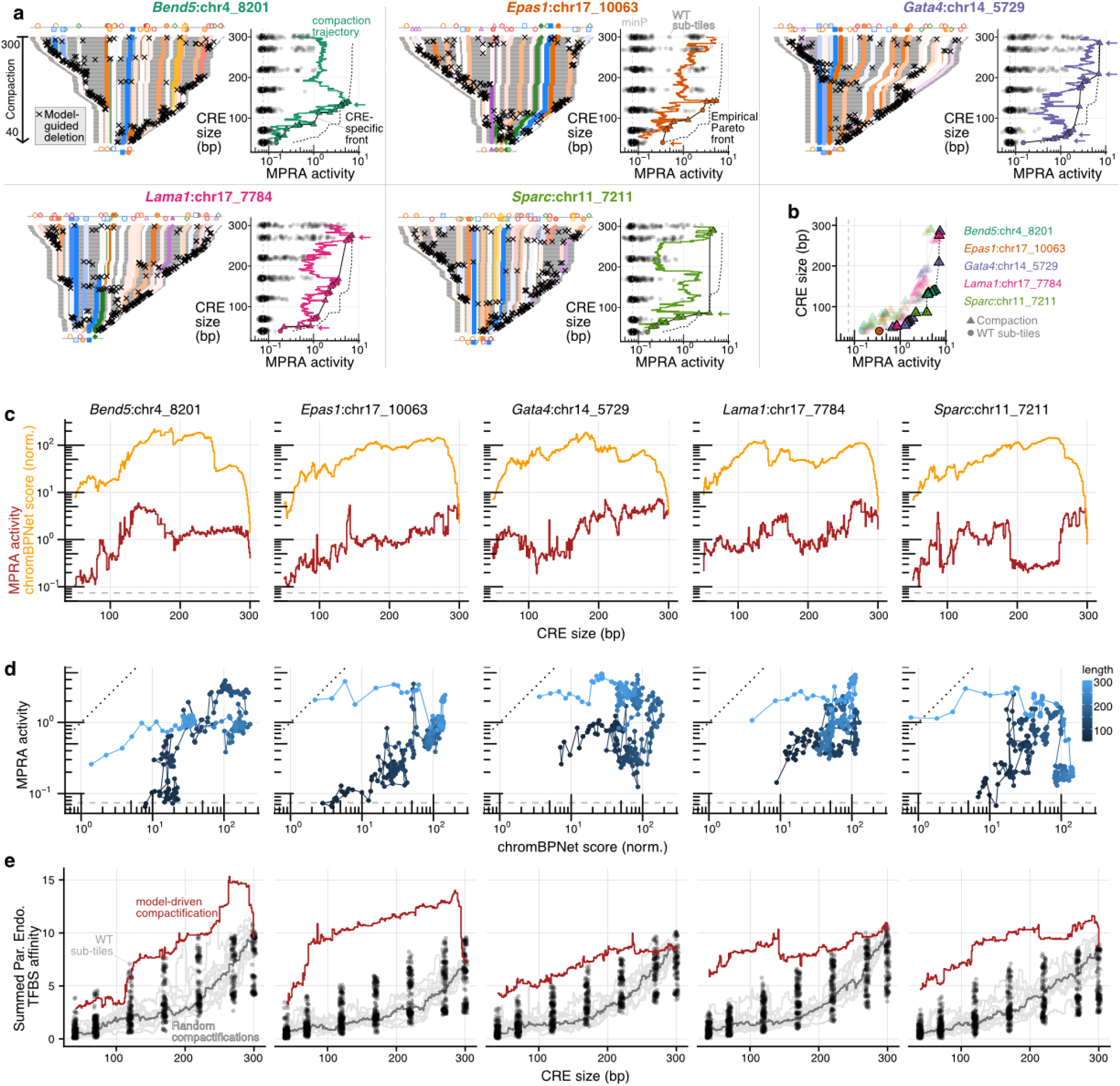
Model-guided compaction of CREs reveals the trade-off between activity vs. length. **(a)** Similar to **Fig. 4a-b** but here for all 5 model CREs. **(b)** Reproduction of the global empirical Pareto front shown in **Fig. 4c** for reference. **(c)** Comparison of model accessibility scores (chromBPNet, orange) vs. activities (MPRA, dark red) along compaction trajectories. Model scores are normalized so that the average score of background sequences is the same as experimentally measured minP value. **(d)** Replotting of the data shown in panel **c**, but here comparing the model accessibility scores (x-axis: chromBPNet) vs. activities (y-axis: MPRA) directly, and encoding CRE length with color (light to dark blue: long to short). Dashed line represents the 1:1 line. Substantial correlation (R^2^) is observed for the 3 of 5 elements:: *Bend5*: 0.40; *Epas1*: 0.37; *Gata4*: 0.0026; *Lama1*: 0.16; *Sparc*: 0.27. **(e)** As a measure for the number of TFBS within the sequences, y-axes shows quantification of summed normalized affinity to parietal endoderm CRM TFBSs for model-guided compaction trajectory sequences (dark red), wild-type endogenous tile sequences (points with x-jitter), and sequences from ten random deletion trajectories (*in silico*, not tested experimentally; light gray are individual trajectories, dark gray is median across the 10 trajectories). Model-guided compaction trajectory sequences show elevated TFBS densities compared to both WT endogenous subtile sequences and random deletion trajectory sequences.

**Supplementary Figure 17.**
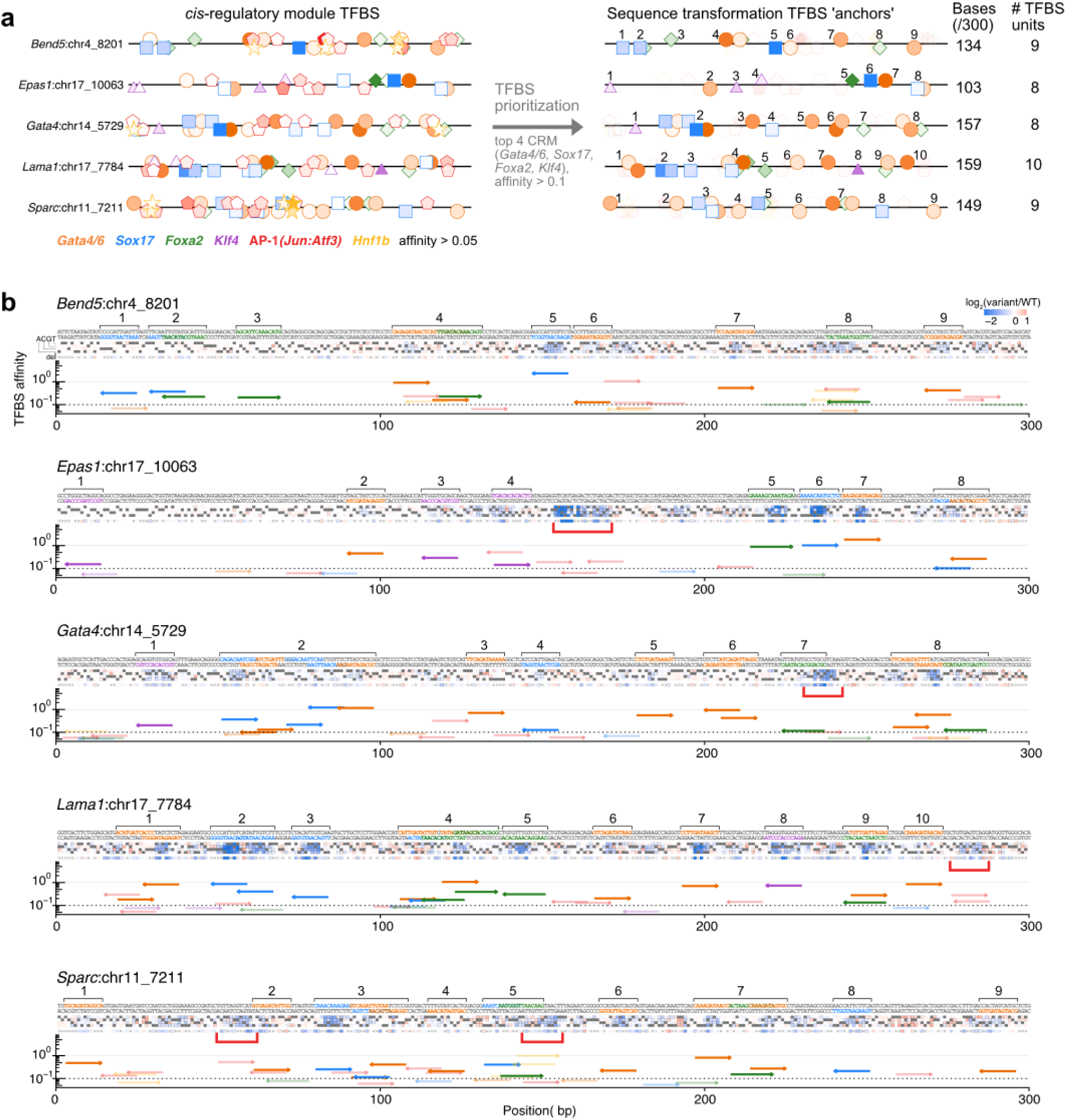
Details on the selection of anchor TFBS units for enhancer derivatization. **(a)** Illustration of the selection process for the definition of TFBS anchors for the derivatization process. To limit the total covered sequence by anchors out of all putative identified CRM TFBS (left, norm. affinity>0.05), the following criteria was applied (right, non-selected TFBS shaded): (1) the top four TFs associated with the highest differential accessibility (Gata4/6, Sox17, Foxa2, and Klf4, **Fig. S1b**) were selected and (2) a threshold of normalized affinity of 0.1 was chosen. These defined TFBSs which were then joined if they overlapped into indivisible units for the derivatization process. Of note, we did not rely on singleton-functional definitions or our saturation mutagenesis data to make our selection. Derivatization anchors cover between 103 and 159 out of 300 bp, and include between 8 and 10 units (some composed of multiple overlapping TFBSs). **(b)** Illustration of the identified anchors (numbered as in panel a) contextualized with the saturation mutagenesis data (heatmap, same as **Fig. 3**, **S9**) and TFBS affinities (arrows indicate directionality of the sites). Red brackets identify AP-1 TFBSs with strong functional impact that were not selected for our procedure (see **Fig. S24** on analysis of synthetic thripsis for additional context).

**Supplementary Figure 18.**
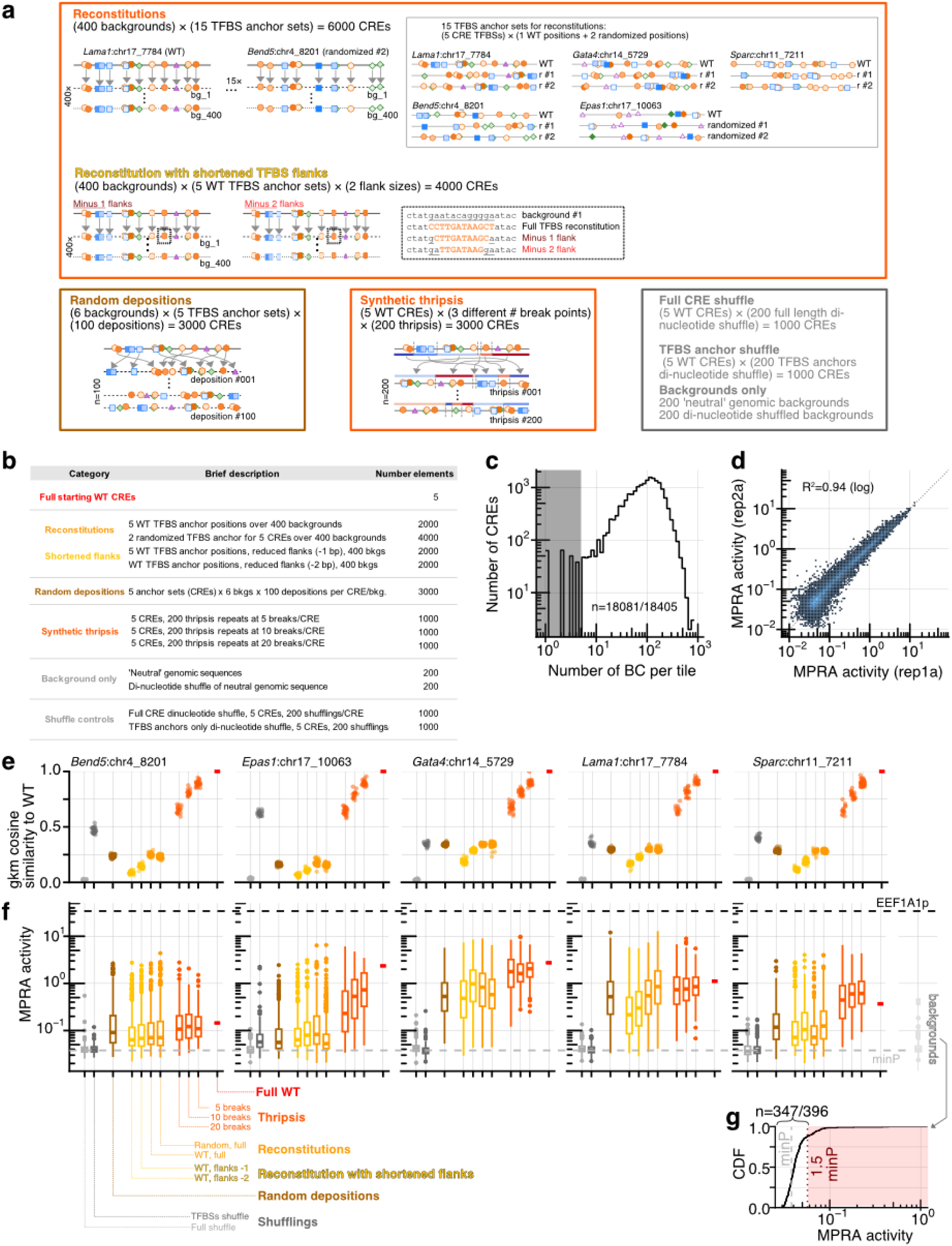
Composition of tested enhancer derivatives and quality metrics of library/MPRA. **(a)** Schematics of the different classes of derivatization and controls included in the derivatization library. Each category of sequence is boxed. Reconstitutions consisted in 400 background DNAs ⨉ 5 sets of anchor sites (one per model CRE) ⨉ 3 anchor positions (original n=2000; two randomized and fixed n=4000). Schematic of reconstituted TFBS positions (yellow) are shown at right. To test for the importance of flanking bases, the WT TFBS positions reconstitutions were also implemented with –1 and –2 bases from the flanks of the anchor sites (taken to be the full extent of the quantitative model used, ranging in size from 12 to 14 bp), n=4000. Inset shows an example Gata4/6 binding site with –1 (10 bp TFBS deposited) and –2 bp on both sides (8 bp TFBS deposited). Random depositions (n=3000, brown) were performed on 6 fixed background DNAs ⨉ 5 set of anchor sites (one per model CRE) ⨉ 100 random anchor depositions. Importantly, the same deposition positions were fixed across backgrounds for a given CRE. Thrispis (n=3000) was performed 200 times on 5 sets of anchor sites (one per CRE) with three number of break points (5, 10, and 20 breaks). The following negative controls were also included: (1) backgrounds alone used for reconstitution and random deposition were also tested (n=200 genomic neutral, n=200 ni-nucleotide shuffled), (2) n=200 full dinucleotide shuffle per starting CREs (n=1000), and (3) n=200 dinucleotide shuffle of just the anchor sites per CRE (n=1000). The starting WT sequences were also profiled, leading to a total of 18405 CREs of 300 bp. **(b)** Summary table of the number of CRE per class. **(c)** Representation of the CRE derivatives in the MPRA library: distribution of number of BCs per element (98.2%, or 18081/18405, of the synthesized sequences were represented with ≥5 BCs, 1.89M BCs total, median coverage 91 BC/CRE). **(d)** Representative correlation between replicates for the MPRA data (R^2^ on log-transformed data = 0.94). **(e)** Comparison of gapped k-mer composition^91^ (size l=11 with k=7 non-gapped position) between enhancer derivatives and their respective wild-type quantified as the cosine similarity between WT and derivative enhancer gkm vectors. Similarities for a sampling of 400 derivatives per starting CREs were computed. **(f)** Box plot summarizing the activity of all classes of derivatization and controls (color aligned with panel **a** categories), faceted by model CRE (left to right panels). Spiked-in minP (basal) only and EEF1A1 (positive) controls are shown as grey and black dashed lines respectively. Background only sequences (light grey) are shown to the right. Full di-nucleotide shuffling of CREs displayed overall low activity (n=887/1000 from 87% to 93% of full di-shufflings per CRE with <1.5-fold minP). Furthermore, di-nucleotide shufflings of only the anchor sites also broadly abrogated activity in all (<1.5-fold minP, from 88% to 96% of anchor shuffling per CRE) but CRE *Epas1*:chr17_10063 (<1.5-fold minP for 50% of anchor shuffling), presumably because of the functionally important cluster of AP-1 binding sites left unperturbed in this process (**Fig. S17**). **(g)** Cumulative distribution of MPRA activity of the background sequences, which had had overwhelmingly low activity over our basal control. With a threshold of 1.5 above minP, n=347/396 (4 background sequences were not represented in our final library) of background sequences displayed no substantial autonomous activity.

**Supplementary Figure 19.**
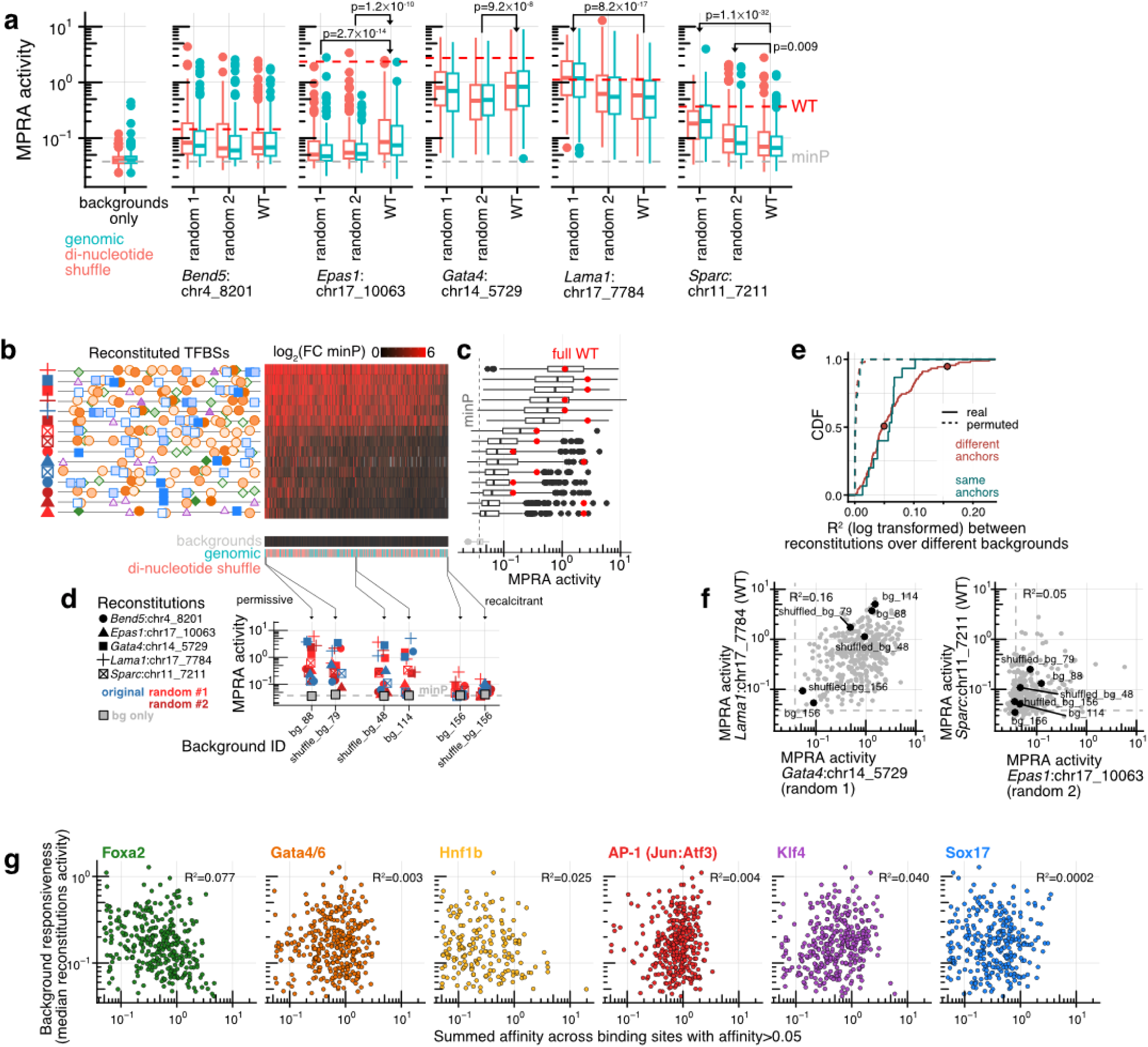
Highly variable background ‘responsiveness’ to TFBS reconstitutions. **(a)** Box plot showing the MPRA activity of different reconstitutions and negative controls. Background-only activities (leftmost) are stratified by sequences sampled from putatively neutral regions of the mouse genome (‘genomic’, cyan, n=200) and di-nucleotide shufflings of these sequences (‘di-nucleotide shuffle’, light red, n=200). There is no difference in overall activity between the two categories (p=0.2 two-sided rank-sum test). Each panel corresponds to reconstitutions from different model CREs, and each model CRE has three depositions (WT positions, and two randomized positions). There was no significant difference in overall responsiveness from genomic or shuffled versions of the backgrounds. Instances with significant differences (one-sided rank-sum test with Bonferroni correction p-value shown) are indicated by the black arrow (e.g., *Lama1*:chr17_7784 randomized 1 more potently inducing activity than its WT counterpart). minP basal activity is shown as dashed grey line. **(b)** Headmap of log_2_ fold-change over minP basal activity (black: low activity, red: high activity) summarizing the full dataset on reconstitutions for the 347 non-autonomously active background sequences (see **Fig. S18g**). Rows are organized according to depositions, with the schematic of the anchor TFBS shown on the left, with the origin following the legend of panel **d** (see also **Fig. S18a**). Columns correspond to different background sequences, which are ordered from most (left) to least (right) responsive. Data for background sequences alone are shown at bottom in a separate line. **(c)** Boxplot for the activity for all backgrounds corresponding to the reconstitution of the aligned rows. Full WT activity for the underlying model CRE is shown as a red dot. **(d)** Examples of permissive, intermediate, and recalcitrant background sequences (arrows indicating identities in heatmap of panel **b**). Even though all are equally autonomous inactive, their response to adding TFBS varies drastically. **(e)** Cumulative distribution of correlations (R^2^ on log-transformed activities) for all possible pairs (15 choose 2 = 105) of reconstitutions across the backgrounds. Pairs from anchors from different model CREs are shown in red, and those from the same model CRE in cyan. Dashed lines show permuted samples comparisons. Points mark the two examples shown in panel **f**. (f) Example pairs showing underlying data to the correlations in panel **e**. Backgrounds shown in panel **d** are marked by black dots. **(g)** Correlations between background sequences pre-existing parietal endoderm CRM TFBS (calculated as the summed affinity of all mapped TFBS with normalized affinity > 0.05) and median responsiveness (across all 15 reconstitutions).

**Supplementary Figure 20.**
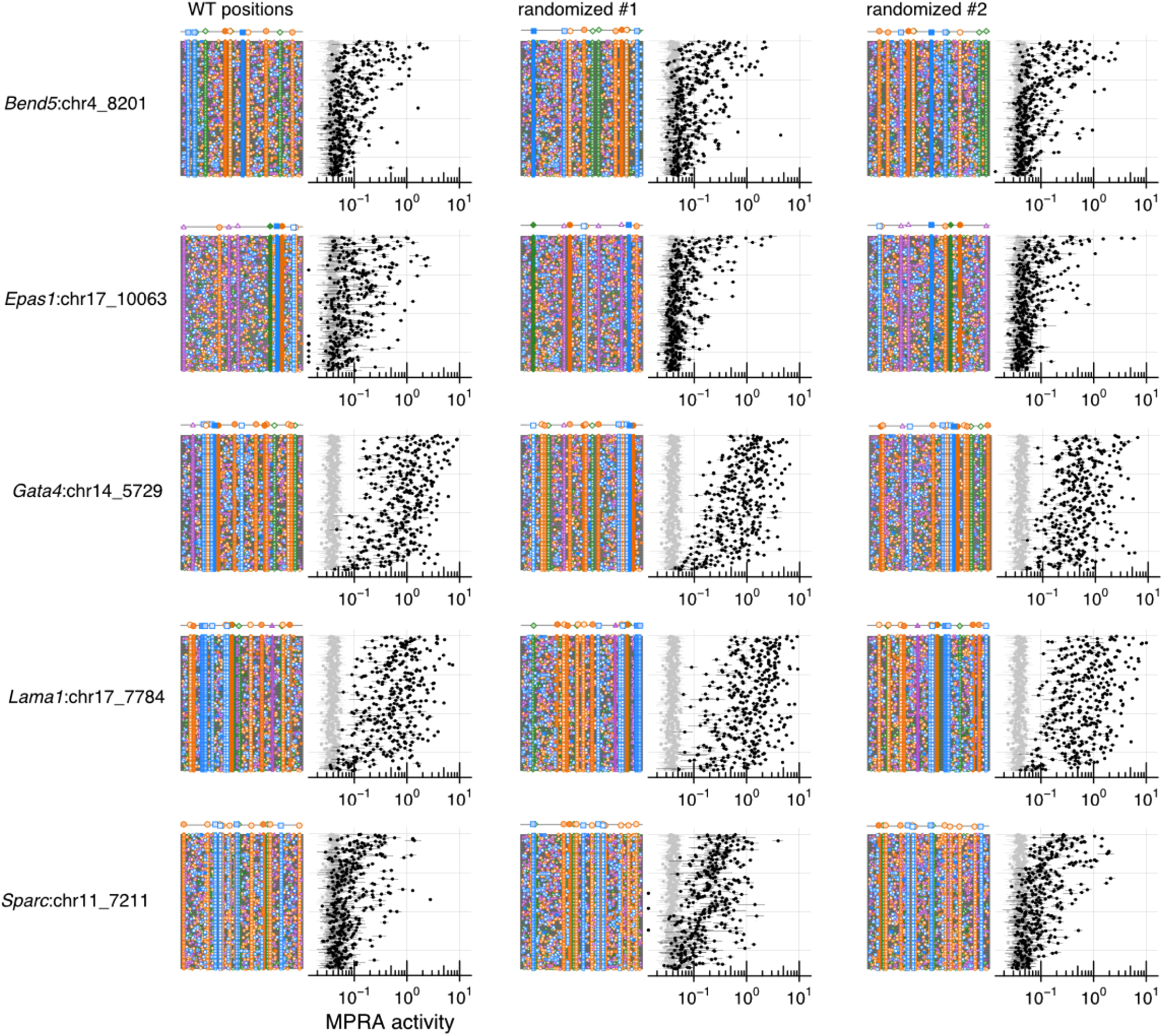
TFBS anchors reconstituted across hundreds of ‘neutral’ DNA background sequences. Reconstitution dataset, equivalent of **Fig. 5b-c** (only showing the reconstitution panels), but for all 15 configurations of reconstitutions (5 model CRE TFBS anchors across 3 sets of positions: one WT and two randomized), organized by model CREs (rows) and reconstituted positions (columns). All background sequences are ordered from top to bottom in the same way across panels, as determined by the overall responsiveness across reconstitutions (top: high response, bottom: low response). Only non-autonomously active backgrounds (activity <1.5 minP basal activity) are included for these plots (n=347/400).

**Supplementary Figure 21.**
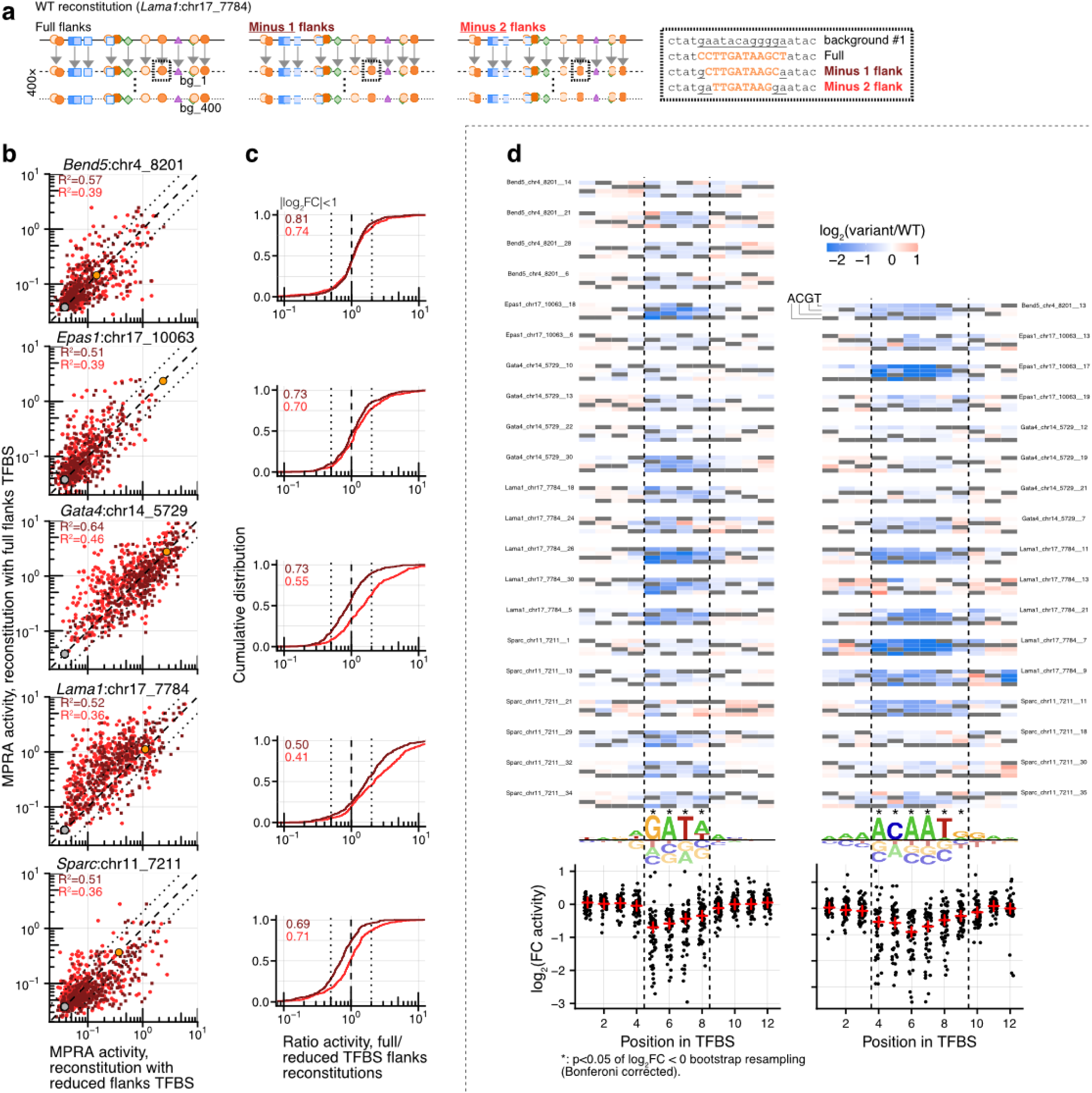
Modest importance of 12-14 bp TFBS flanks for reconstitution activity. Given the importance of bases flanking the core motifs in TF binding^5,89^, a natural explanation for difference in reconstitution activities would be that the different backgrounds lead to small-magnitude modulation in affinity of the anchor sites as a result of changing surrounding sequences. To explicitly test this, reconstitutions with reduced flanks were performed (with WT TFBS anchor positions). **(a)** Schematic of reconstitution with TFBS of reduced flanks. We tested reconstitutions with reduced flanks by removing one (minus 1 flanks) and two (minus 2 flanks) base pairs from both ends of the TFBS (5 model CREs across 400 backgrounds, n = 4000 sequences). The rightmost dashed-line box shows an example Gata4/6 binding site with the full vs reduced flanks sequences deposited in the background. **(b)** Correlation between activity of full reconstitutions (y axis) vs. reduced flanks (x axis, dark red: minus 1, light red: minus 2). R^2^ of log-transformed values are shown in the plot. Dashed and dotted lines show the 1:1 and two-fold deviations respectively. **(c)** Cumulative distribution of ratio of activities between full and reduced. Proportion of reconstitutions with less than 2-fold difference between full and reduced indicated on the plot. **(d)** Correlations are substantial between full and reduced, they remain incomplete, suggesting flanks could indeed substantially contribute. As another line of evidence against this possibility, we aligned our saturation mutagenesis data for the most prevalent singleton-functional TFs (Gata4/6: left, and Sox17: right). Top of panel shows saturation mutagenesis heatmap (log_2_FC vs. WT, as in **Fig. 3**), one sub-panel per TFBS, with CRE and TFBS ID indicated. The bottom of panels shows aggregated log_2_FC (bottom) as a function of position within the TFBS (horizontal jitter for visualization, median shown as red +). Significant effects (log_2_FC<0 in resampling bootstrap at p<0.05 with Bonferroni correction) are indicated by *. Probound^74^ binding model weights are shown as a sequence logo. The empirically determined core region is delimited by vertical dashed lines. Within our saturation mutagenesis data, flanking positions not included in the reduced reconstitutions indeed do not contribute substantially to activity. Consistently with the correlations in background responsiveness across reconstitutions, these point to complex sequence determinants beyond the flanks of TFBS anchors.

**Supplementary Figure 22.**
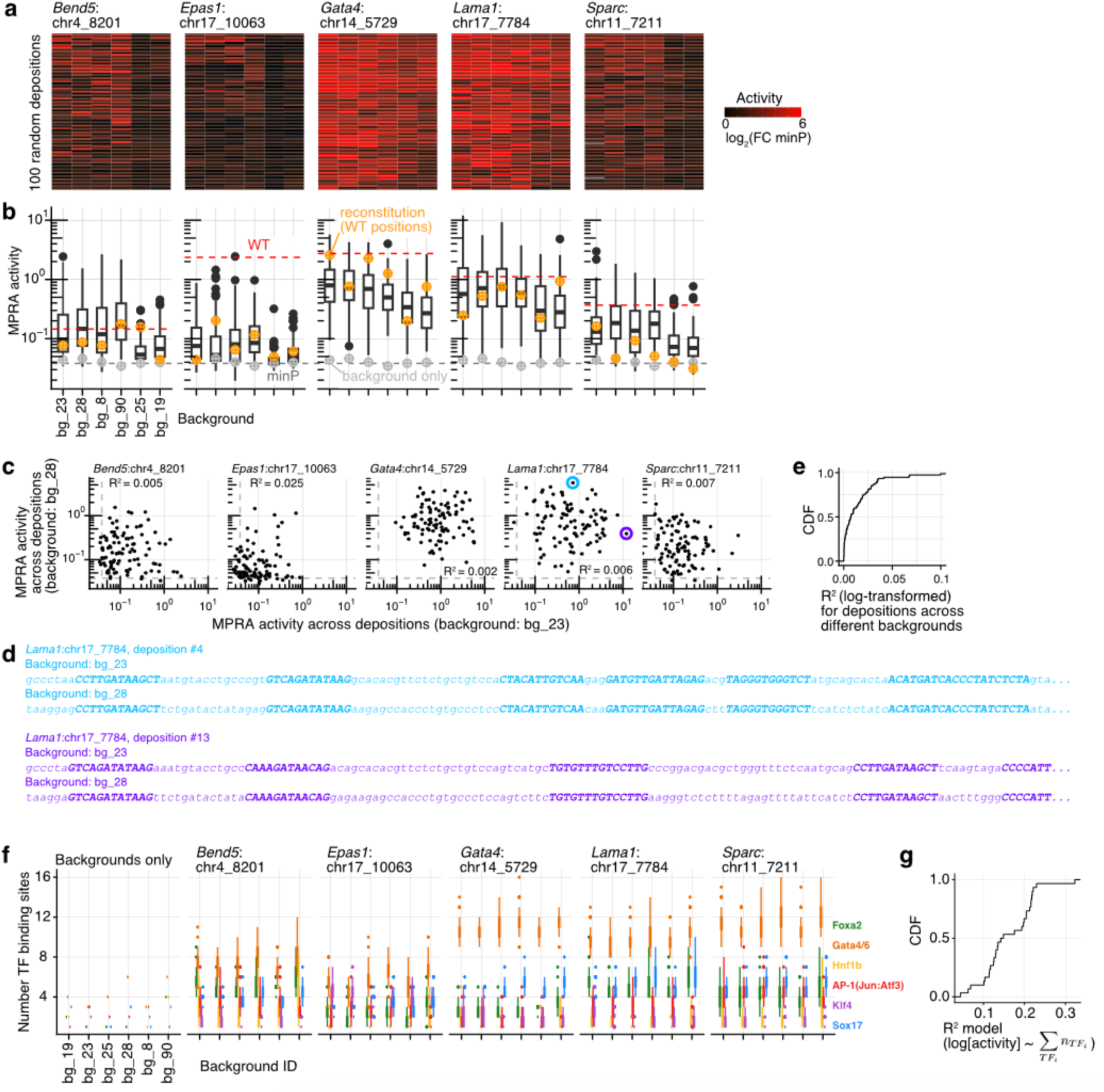
Additional analysis of random TFBS depositions. **(a)** Heatmaps (one per model CRE) of random deposition activities (log_2_FC over minP) over deposition positions (rows, fixed across backgrounds internally for each CRE) and background DNA (columns). Deposition and backgrounds are ordered by activities. **(b)** Boxplot aggregating the data over deposition from panel **a**. minP baseline is shown as gray dashed line and WT CRE activity as a red dashed line. Background-only activity is shown as gray + and WT TFBS anchors reconstituted (equivalent to a random deposition with the WT positions) in the same background as orange +. Note how the WT reconstitutions fall across the full range of activities of random depositions (see **Fig. 5g**). **(c)** Example of correlation (R^2^ on log-transformed) between two background DNA (bg_23 vs. bg_28) across 100 depositions for 5 CREs. Correlations are overall low. Colored dots in *Lama1*:chr17_7784 CRE indicate examples with strongly discordant activities across backgrounds shown in panel **d**. **(d)** First 145 bp of sequences for CREs highlighted in panel **c** (bg_23 and bg_28 with depositions #4 and #13) with concordant coloring. Random depositions are fixed, but background sequences differ. TFBS anchors are shown bolded and in capital letters (background, lowercase). **(e)** Cumulative distribution of R^2^ across pairs of background/CRE pairs (6 choose 2 = 15 pairs with 5 CREs). **(f)** Distribution of mapped number of TFBS (>0.1 normalized affinity) across random depositions stratified by CREs and backgrounds. Boxplot illustrates the slight variability in the number of binding sites across depositions putatively due to difference in chimeric sequences at junctions of TFBS and background. On average, we observe changes between 1.5 to 2.9 binding sites (span of the central 80% range) per CRE per TF over a fixed anchor set and background. **(g)** Simple linear models (one per CRE/background pair) relating these slight TFBS number variations to activity hold limited predictive potential (average R^2^ of <0.2), underscoring that ‘chimeric’ TFBS are probably a minor determinant of activity.

**Supplementary Figure 23.**
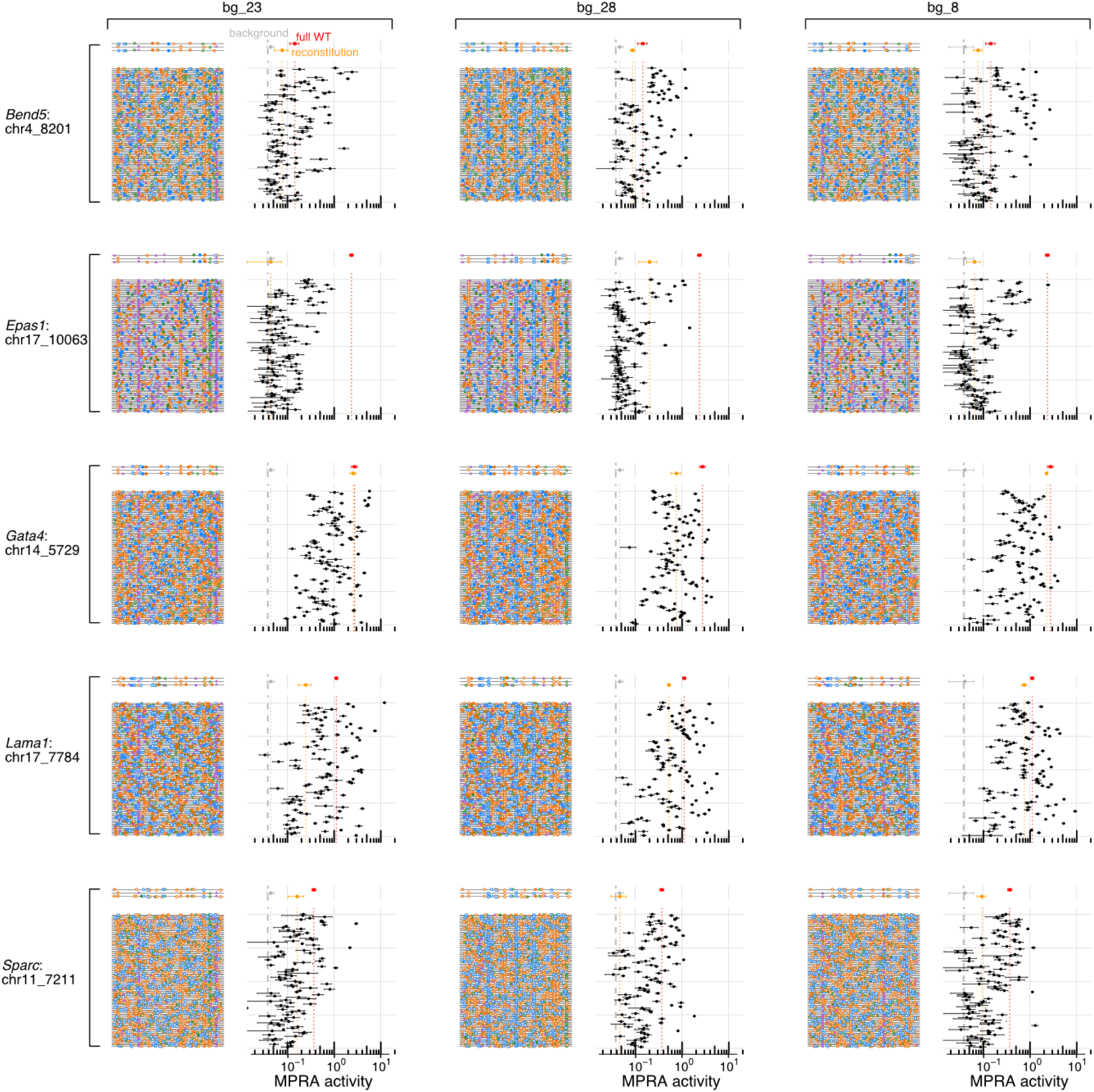
Thousands of random TFBS depositions across 5 CREs ⨉ 6 backgrounds (part 1) Random deposition dataset, analogous to **Fig. 5e-f**, but on the full set of 30 CRE/background pairs. Depositions (fixed across backgrounds for each model CREs) are ordered based on overall activity.

**Supplementary Figure 23.**
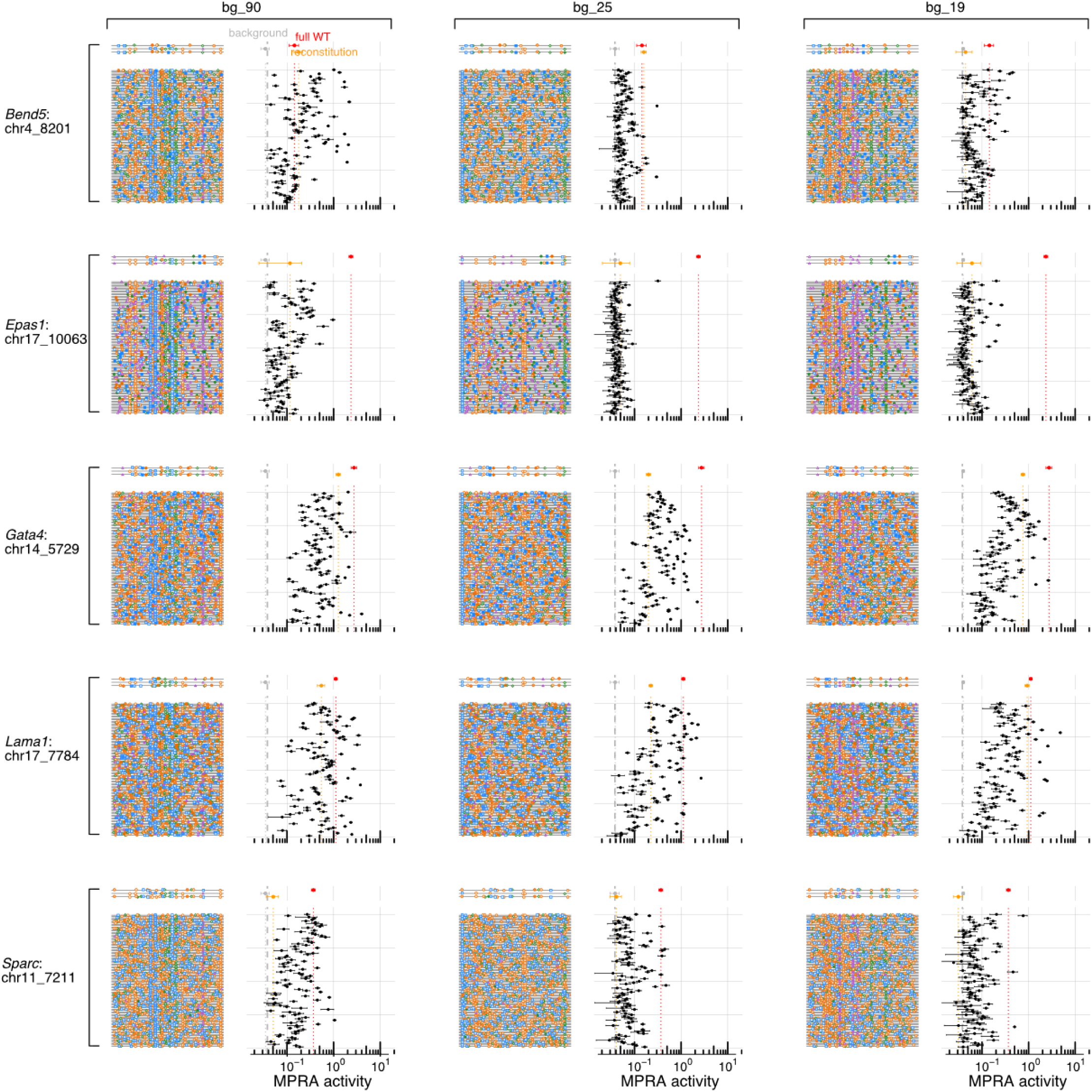
Thousands of random TFBS depositions across 5 CREs ⨉ 6 backgrounds (part 2). Random deposition dataset, analogous to **Fig. 5e-f**, but on the full set of 30 CRE/background pairs. Depositions (fixed across backgrounds for each model CREs) are ordered based on overall activity.

**Supplementary Figure 24.**
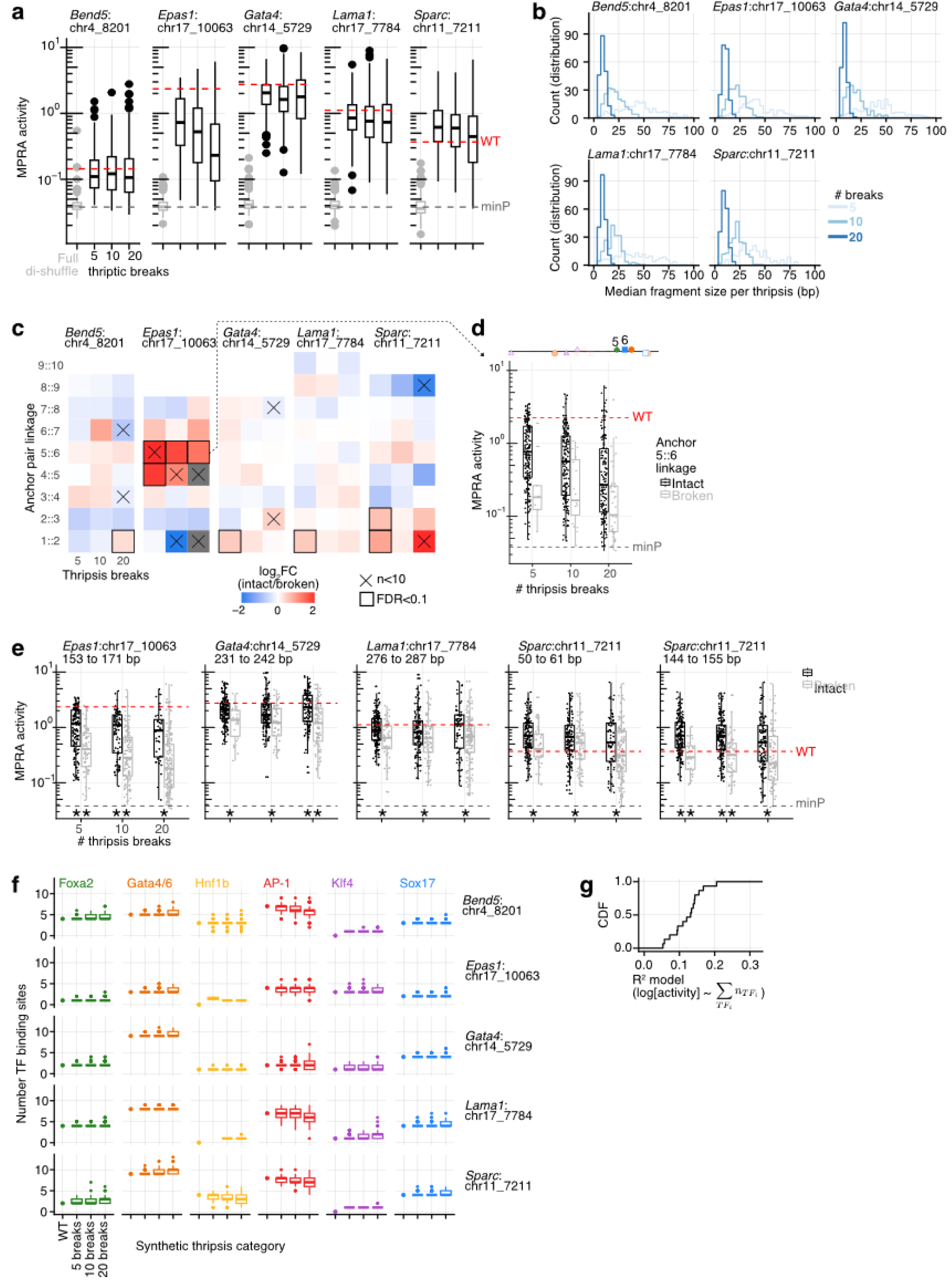
Analysis of synthetic thripsis derivatives. **(a)** Box plot of CRE derivatives MPRA activity stratified by model CRE (panels) and number of thriptic break points (x-axis). For reference, the full CRE di-nucleotide shufflings for each CREs are included (gray box plot), minP-only (gray dashed line) and corresponding model CRE WT (red dashed line). **(b)** Distribution of median fragment length resulting from the synthetic thripsis operation per CRE. The average median fragment size per thripsis ranged from 45 to 8 bp from 5 to 20 break points. This is slightly shorter than the expected *L/(n+1)* because of the anchor-avoiding constraint. **(c)** For each consecutive pairs of TFBS anchors, the thripsis CREs were split between those preserving or not the linkage (i.e., with or without one or more break point between them) and the difference in activity quantified. FC between medians of intact/broken linkage is shown as a heatmap. Linkages with FDR<0.1 (rank-sum p-value with Benjamini-Hochberg correction) are highlighted with dark edges. Linkages with fewer than 10 observations are marked by an ✕ (and those with none, e.g., for anchors that are far apart, are grayed out). Only the TFBS 5::6 linkage in CRE *Epas1*:chr17_10063 is significant across the three thripsis conditions (FDR<0.03) **(d)** Underlying data for the 5::6 linkage data pointed at (dashed line with arrow) in panel **c**. Each point is the activity of a thripsis CRE, split between linkage preserving [black] and linkage abrogating [gray] instances. **(e)** Similar analysis as panels c-d, but now with functionally important AP-1 TFBS (not included in the anchor set, indicated in **Fig. S17b**). Thripsis activities are stratified by whether the indicated region is intact (black) or not (gray). Integrity of the regions are all significantly related to activity (ranksum test with Benjamini-Hochberg correction, *: 0.0005 ≤ FDR < 0.05, **: FDR < 0.0005). **(f)** Quantification of the number of CRM TFBS (columns) per CRE (normalized affinity > 0.1) across different model CREs (rows) organized by different number of thriptic break points shown as a boxplot. These provide information about the likelihood of chimeric junctions creating new TFBS. Quantification for WT model CREs are included as a single point. Overall, median change in the number of TFBS is low (5 breaks: 0.5, 10 breaks: 0.8, 20 breaks: 1.3). (g) These changes in TFBS numbers can be used to predict log-activity with a linear model, but they hold corresponding little explanatory power (on average R^2^<0.13). Shown is the distribution of R^2^ for the 15 situations (5 model CREs by 3 thripsis set).

**Supplementary Figure 25.**
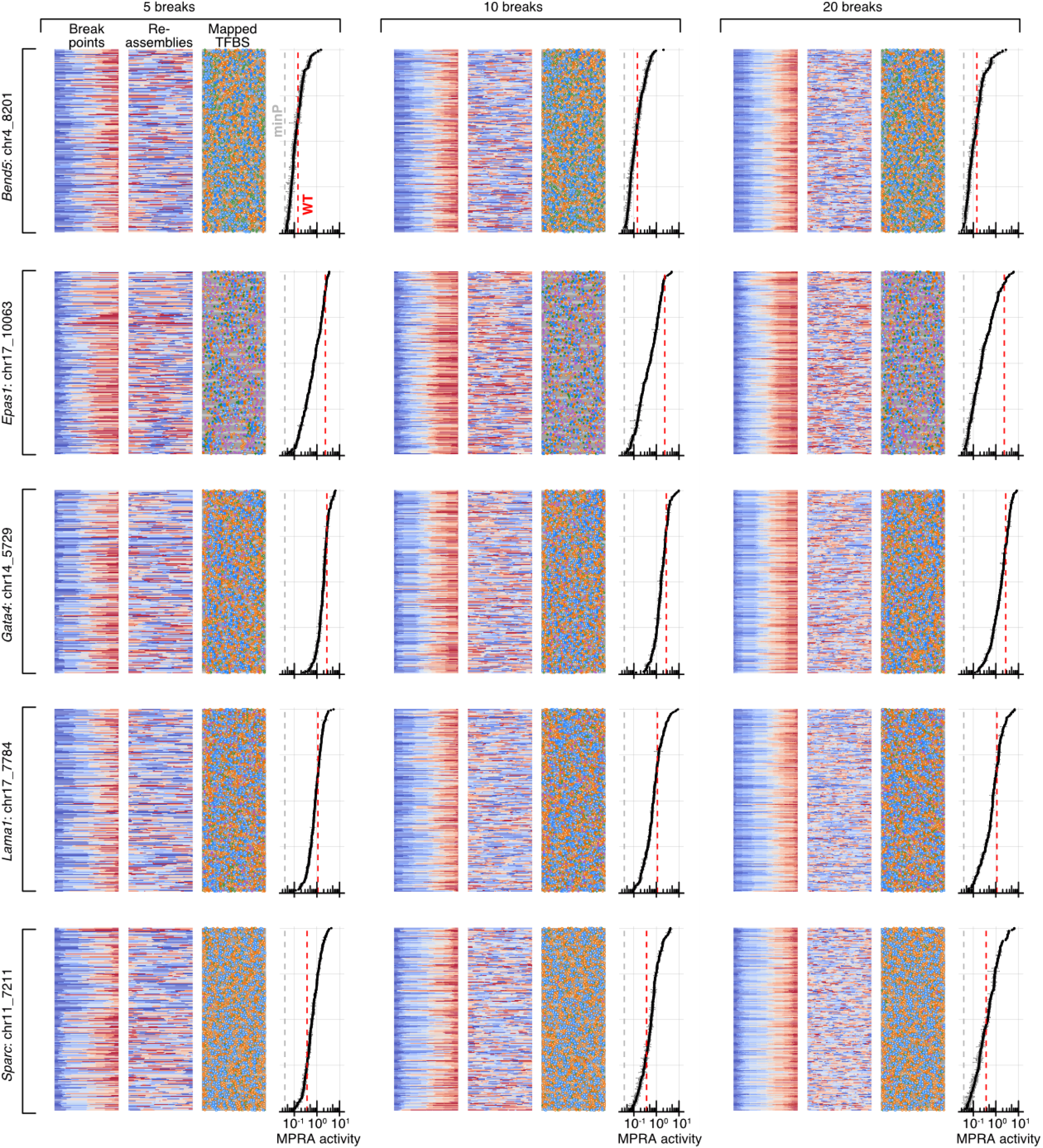
Thousands of synthetic thripsis from five model CREs. Analogous to **Fig. 5i-j**, but for all different numbers of thriptic fragments and starting CREs. Since comparisons were not possible across thripsis (unlike for reconstitutions/random depositions), thripsis operations are ordered vertically by activity.

